# Bipartite left-right sided endocrine system: processing of contralateral effects of brain injury

**DOI:** 10.1101/2023.06.28.546857

**Authors:** Hiroyuki Watanabe, Yaromir Kobikov, Olga Nosova, Daniil Sarkisyan, Vladimir Galatenko, Liliana Carvalho, Gisela H. Maia, Nikolay Lukoyanov, Igor Lavrov, Mathias Hallberg, Jens Schouenborg, Mengliang Zhang, Georgy Bakalkin

## Abstract

The crossed descending neural tracts set a basis for contralateral effects of brain injury. In addition, the left-right side-specific effects of the unilateral brain lesions may be mediated by neurohormones through the humoral pathway as discovered in animals with disabled descending motor tracts. We here examined if counterparts of the endocrine system that convey signals from the left and right brain injuries differ in neural and molecular mechanisms. In rats with completely transected cervical spinal cords a unilateral injury of the hindlimb sensorimotor cortex produced hindlimb postural asymmetry with contralateral hindlimb flexion, a proxy for neurological deficit. The effects of the left and right side brain lesions were differently inhibited by antagonists of the δ-, κ- and µ-opioid receptors suggesting differential neuroendocrine control of the left-right side-specific hormonal signaling. Bilateral deafferentation of the lumbar spinal cord eliminated hormone-mediated effects of the left-side brain injury but not the right-side lesion suggesting their afferent and efferent mechanisms, respectively. Analysis of gene-gene co-expression patterns identified the left and right side-specific gene regulatory networks that were coordinated across the hypothalamus and lumbar spinal cord through the humoral pathway. The coordination was ipsilateral and perturbed by brain injury. These findings suggest that the neuroendocrine system that conveys left-right side-specific hormonal messages from injured brain is bipartite, contributes to contralateral neurological deficits through asymmetric neural mechanisms, and enables ipsilateral coordination of molecular processes across neural areas along the neuraxis.

**GRAPHICAL ABSTRACT:** 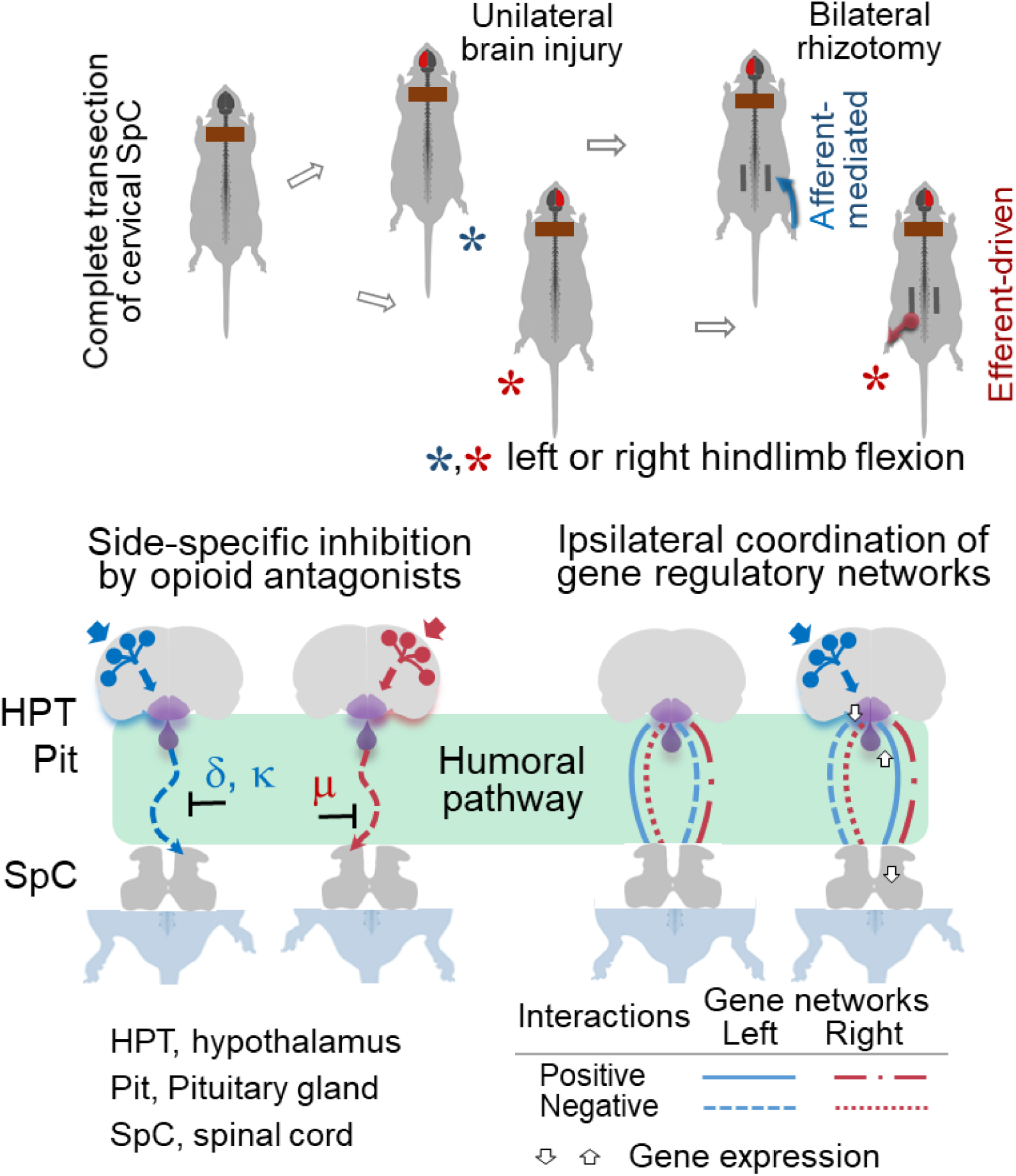

## INTRODUCTION

The central neurology concept known as *cross association* states that each cerebral hemisphere is functionally connected to the contralateral side of the body through the decussating neural tracts (Bakalkin, 2022; Lemon, 2008; Louis, 1994; Purves, Augustine, & Fitzpatrick, 2001; Simon, 2018; Smith, Paton, Chakrabarty, & Ichiyama, 2017). Traumatic brain injury and stroke in patients, and brain lesions in animal experiments cause postural and sensorimotor deficits that are generally contralateral and include asymmetric posture and reflexes (Dewald, Beer, Given, McGuire, & Rymer, 1999; Serrao et al., 2012; Spaich, Hinge, Arendt-Nielsen, & Andersen, 2006; Wilson et al., 2017; Zhang et al., 2020). After a unilateral injury to the hindlimb sensorimotor cortex, animals exhibit the hindlimb postural asymmetry (HL-PA) with contralesional limb flexion, and asymmetry of reflexes with greater activity on the contra *vs*. ipsilesional side (Wolpaw, 2012; Zhang et al., 2020). A cause of the contralateral effects of brain lesions has been considered as solely neuroanatomical – based on the decussation of the descending neural pathways (Bakalkin, 2022; Schutta, Abu-Amero, & Bosley, 2010; Vulliemoz, Raineteau, & Jabaudon, 2005).

Emerging evidence indicates that, in addition to neural mechanisms, the contralateral effects of brain lesions are mediated through the humoral pathway by neurohormones that produce either the left or right side-specific effects (Bakalkin, 2022; Lukoyanov et al., 2021; Watanabe et al., 2021; Watanabe et al., 2020). The humoral signaling was identified in animals whose descending neural tracts were disabled by complete transection of the spinal cord and then the brain was injured. Strikingly, rats with transected thoracic spinal cord and a unilateral injury of the hindlimb sensorimotor cortex developed contralateral hindlimb flexion, and asymmetry in hindlimb withdrawal reflexes and gene expression patterns in the lumbar spinal cord. The left hindlimb was flexed after the right-side brain injury, while injury of the left hemisphere induced the right hindlimb flexion. Hypophysectomy abolished these effects, whereas serum from animals with brain injury induced HL-PA in animals with intact brain. Arg-vasopressin and β-endorphin were identified as molecules that mediate the effects of the left-sided brain injury. They are produced in the hypothalamic-pituitary system, and evoke HL-PA with right hindlimb flexion in animals with intact brain (Lukoyanov et al., 2021). Thus the left-right side-specific neuroendocrine signals may bypass descending neural tracts and convey information on the side of brain injury. These neurohormones could be a part of a general mechanism that spans the nervous system or the entire body after their release from the hypothalamus or pituitary gland, and thus enables differential neuroendocrine control of the left and right body sides.

How this topographic left-right side-specific neuroendocrine system (T-NES) is organized and functions is still an enigma. Three stages may be envisaged; the encoding of signals from the left and right hemispheres into the left and right side-specific neurohormones, respectively; their release from the hypothalamus and pituitary into the blood; and the decoding of these hormonal messages into left-right sided responses in the spinal cord and peripheral nervous system (Bakalkin, 2022; Lukoyanov et al., 2021). Different top-down signaling from two anatomically symmetric hemispheres requires bipartite, lateralized and hemisphere (side) specific organization of the T-NES.

We reasoned that the T-NES consists of two counterparts that differentially process and convey the left and right side-specific messages, and whose activities are balanced in intact rats whereas may be impaired by a unilateral impact. These two parts may be mirror-symmetric in their structure, e.g., in cell type composition and connectivity, or may differ in their internal architecture and exploit different endocrine, neural and molecular mechanisms to produce symmetric physiological outcomes.

In this study, we addressed these hypotheses with aim to characterize the T-NES counterparts and to reveal their lateralized features. The HL-PA, a proxy for neurological deficit with binary, left or right sided outcomes including directional asymmetry in posture and motor functions, was used to characterize and compare the left and right T-NES counterparts. To analyze the T-NES, neural pathways between the brain and lumbar spinal cord were disabled by complete spinal cord transection. The left and right sided unilateral injury of the hindlimb sensorimotor cortex was applied to evoke signaling through the left and right T-NES counterparts and, by this virtue, to separately analyze their features. The cortex was injured by ablation to restrict a damaged area to the hindlimb sensorimotor cortex and examine specific changes in hindlimb motor functions and lumbar spinal circuits. In this biologically relevant acute injury model pathological factors that may interfere with the T-NES functions, e.g., neuroinflammation, widespread damage to neurons, axons and blood vessels that are produced by traumatic brain injury and stroke (Fu, Liu, Anrather, & Shi, 2015; Ng & Lee, 2019) were largely excluded.

Analysis of signaling from the sensorimotor cortex injured on the left or right side demonstrated that the T-NES is binary and functionally asymmetric; the left or right side T-NES counterparts differently target the contralateral afferent and efferent processes controlling hindlimb functions. Experiments with opioid receptor antagonists confirmed that the T-NES is bipartite, and that the “left” and “right” messages are side-specifically controlled by the δ-, κ- and µ-receptors. Analysis of gene expression as the readout suggested that UBI affects the hypothalamus and pituitary gland, that the side-specific molecular processes are coordinated between the hypothalamus and the lumbar spinal cord by the T-NES, and that this coordination is ipsilateral and impaired by a unilateral brain lesion.

## RESULTS

### The unilateral brain injury (UBI)-induced HL-PA in rats with transected cervical spinal cord

Previous study suggested that the asymmetric effects of the unilateral brain injury on hindlimb posture and reflexes in animals with completely transected thoracic spinal cords are mediated through the humoral pathway (Lukoyanov et al., 2021). Theoretically, the top-down asymmetric signaling from a unilaterally injured brain to the hindlimb muscles may be mediated through the preganglionic sympathetic neurons that are located in the superior thoracic segments projecting to the paravertebral sympathetic chain, and further through their postganglionic fibers to the hindlimbs (Hotta, Iimura, Watanabe, & Shigemoto, 2021; Lee, Lois, Troupe, Wilson, & Yates, 2007; McCall, Miller, & Yates, 2017). To explore this possibility, the hindlimb responses to UBI were analyzed in the rats with complete spinal cord transection performed at the level rostral to the preganglionic sympathetic neurons. A 3-4-mm segment of the C6-C7 spinal cord was excised, and then the hindlimb representation area of the sensorimotor cortex was unilaterally ablated by suction and HL-PA was analyzed.

The lesion sites extended 4.8 – 5.6 mm rostrocaudally and 2.6 – 3.1 mm mediolaterally. The lesion was 1.0 – 1.5 mm in depth and did not affect the white matter below the cortex (**Figure 1—figure supplement 1**). The lesion volumes of the cortices (mean value ± SD) without correcting for tissue shrinkage due to fixation were similar in the left (7.9 ± 2.3 mm^3^, n = 5) and right (7.8 ± 2.5 mm^3^, n = 5) UBI rats (P = 0.92; two tailed t-test).

**Figure 1.**
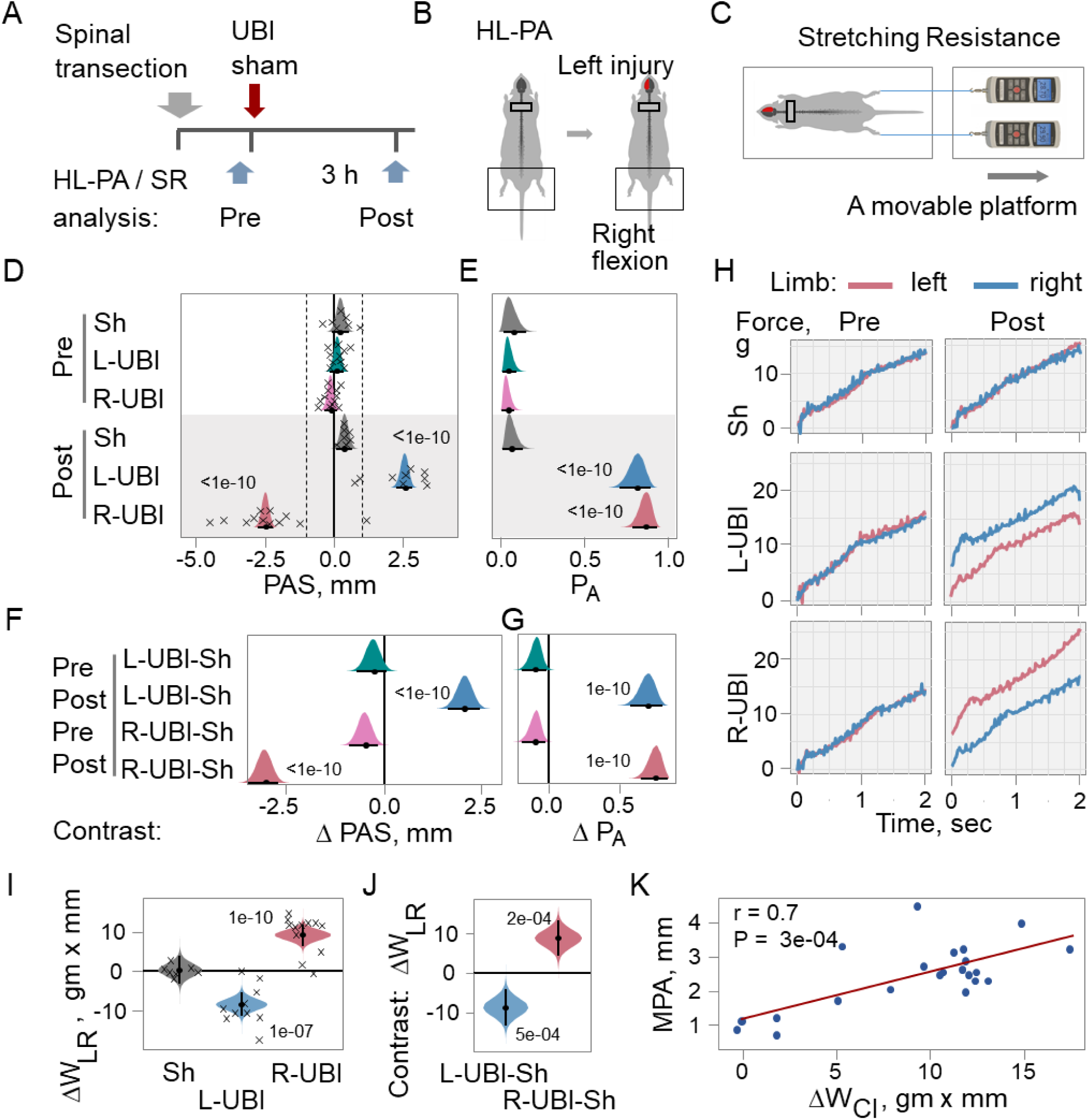
Hindlimb asymmetry in posture (HL-PA) and stretching resistance (SR) induced by the unilateral ablation of the hindlimb representation area of sensorimotor cortex (UBI) in rats with completely transected spinal cord. (**A**) Experimental design. The spinal cord was transected at the C6-7 level that was followed by left UBI (L-UBI; n = 10), right UBI (R-UBI; n = 12), or sham surgery (Sh; n = 7). The asymmetries were analyzed before (Pre) and 3 h after (Post) UBI or sham surgery. (**B**) The UBI-induced HL-PA was manifested as flexion of the left or right hindlimb. (**C**) The stretching resistance was analyzed as the amount of mechanical work W to stretch a hindlimb, calculated as integral of stretching force over 0-10 mm distance. Stretching force was measured by the micromanipulator-controlled force meter device consisted of two digital force gauges fixed on a movable platform. Two silk threads were hooked to the force gauges, their other ends were glued to the leg nails, and the legs were placed in symmetric position and stretched at 5 mm/sec speed. (**D**) The postural asymmetry size (PAS) in millimeters (mm), and (**E**) the probability to develop HL-PA (P_A_) above 1 mm threshold (denoted in **D** by vertical dotted lines). (**F,G**) Differences (contrasts) between the UBI and sham groups in the PAS and P_A_ before (Pre) and after (Post) UBI or sham surgery. (**H**) Representative traces of the stretching force recorded from the left and right hindlimbs of rats with transected spinal cord before the UBI and sham operation and after injury. (**I**) Differences in stretching force calculated between the left and right hindlimbs as ΔW_LR_ = (W_Left_ - W_Right_) in gm × mm analyzed after UBI or sham surgery (Post). (**J**) Differences (contrast) between the UBI and sham surgery groups in ΔW_LR_ analyzed 3 h after UBI or sham surgery (Post). (**K**) Pearson correlation between the postural asymmetry magnitude (MPA) and differences in the work between the contra- and ipsilesional hindlimbs ΔW_CI_ = (W_Contra_ – W_Ipsi_) in gm × mm. Data are presented for L-UBI and R-UBI groups analyzed 3 hours after brain surgery. The PAS, P_A_, ΔW_LR_ and contrasts are plotted as median (black circles), 95% HPDC intervals (black lines), and posterior density (colored distribution) from Bayesian regression. Negative and positive PAS values are assigned to rats with the left and right hindlimb flexion, respectively. Significant effects on asymmetry and differences between the groups: 95% HPDC intervals did not include zero, and adjusted P-values were ≤ 0.05. Adjusted P is shown for differences identified by Bayesian regression. Crosses in **D** and **I** denote the PAS and ΔW_LR_ values for individual rats, respectively. **Source data:** The EXCEL source data file “masterfile-210807.xlsx” and source data folder “/HL-PA/data/SF/”.

#### Analysis of HL-PA

HL-PA was analyzed before (designated as Pre) and 3 hours after the UBI or sham surgery (Post) (**Figure 1A,B**), by both the hands-on and hands-off methods of hindlimb stretching followed by photographic and / or visual recording of the asymmetry in animals under pentobarbital anesthesia (Lukoyanov et al., 2021; Zhang et al., 2020). Data obtained by these two methods are well correlated (**Figure 1—figure supplement 2**). The HL-PA data on **Figure 1** and throughout the paper are presented for the hands-off assay. HL-PA was characterized by i) the postural asymmetry size (PAS) in mm; ii) the magnitude of postural asymmetry (MPA) in mm; and iii) the probability to develop HL-PA (P_A_). In the P_A_ calculations, the rats with the MPA > 1 mm were defined as asymmetric; the 1 mm MPA was 94^th^ percentile in rats before UBI or sham surgery and after sham surgery.

In rats with transected cervical spinal cords, UBI induced HL-PA (**Figure 1D,E**). The PAS and P_A_ in rats with left and right UBI were at least 7-fold greater than in rats before UBI or sham surgery, or after sham surgery. The P_A_ and MPA did not differ between the left- and right-side UBI groups. Differences in the PAS and P_A_ between the UBI sham surgery groups were strong and highly significant (**Figure 1F,G**). The response to the injury was developed on the contralesional side; the left or right UBI induced the right and left hindlimb flexion, respectively. The PAS and P_A_ in the UBI rats with transected cervical spinal cords were similar to those that were previously reported for the UBI animals with intact and transected spinal cords (Lukoyanov et al., 2021; Watanabe et al., 2021).

#### Hindlimb stretching resistance: effects of UBI

We next examined the UBI effects in rats with completely transected cervical spinal cord on biomechanical properties of the contra- and ipsilesional hindlimbs. The passive hindlimb musculo-articular resistance to stretching was assessed in the anesthetized rats before and 3 hours after UBI or sham surgery. The asymmetry in the resistance was assessed as i) the difference in the work between the left- and right hindlimbs ΔW_LR_ = (W_Left_ – W_Right_) where W_Left_ and W_Right_ were the W applied to stretch the left and right hindlimbs (**Figure 1I**); and ii) the left / right asymmetry index for the work AI_LR_ = log_2_ (W_Left_ / W_Right_) (**Figure 1—figure supplement 3**). The ΔW and AI may differently depend on the stretching distance, and therefore were both analyzed. Difference in the work between contra-(C) and ipsilesional (I) hindlimbs were compared by computing ΔW_CI_ = (W_C_– W_I_) (**Figure 1K**) and AI_CI_ = log_2_ (W_C_ / W_I_) (**Figure 2—figure supplement 1**).

**Figure 2.**
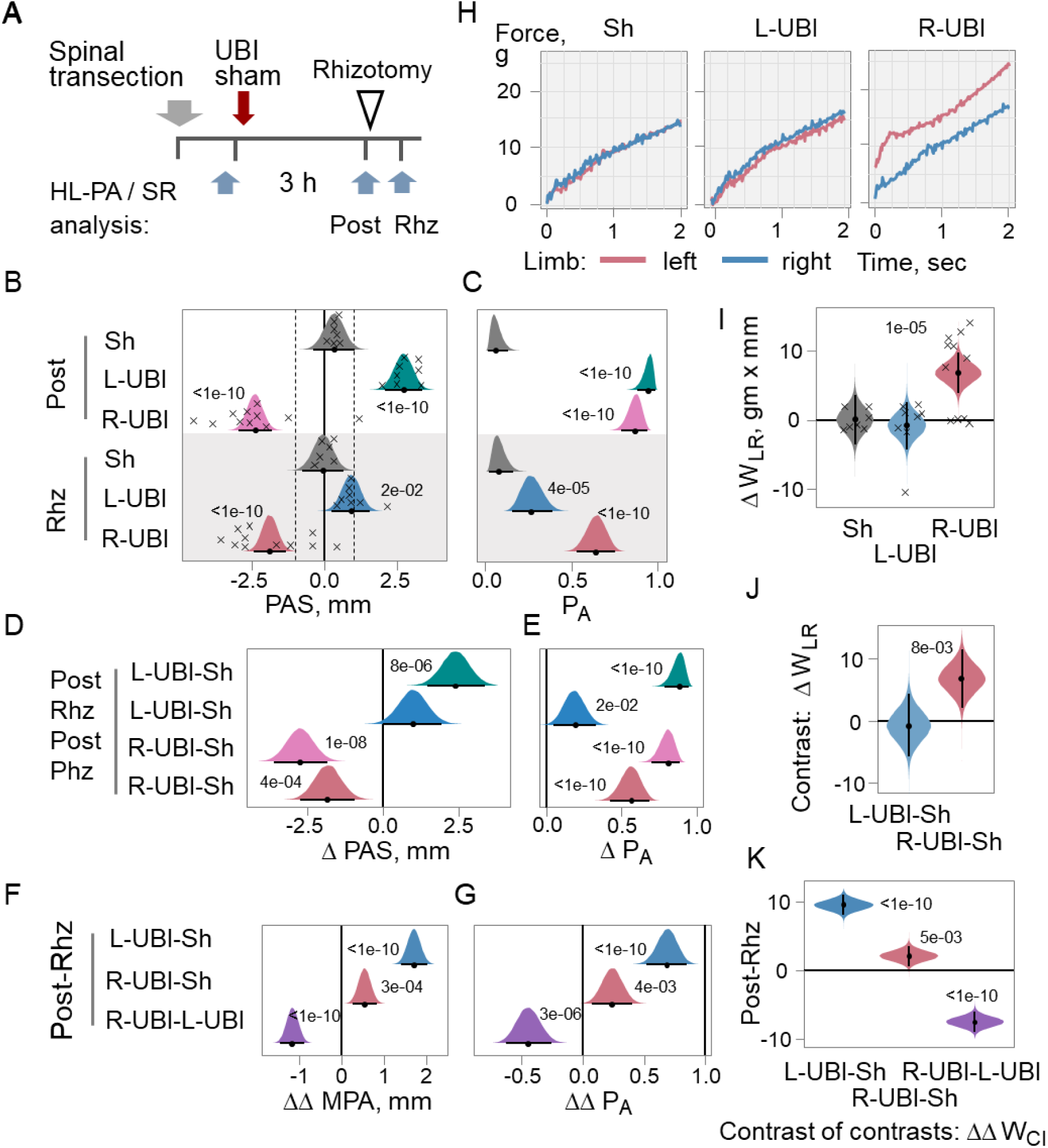
Effects of bilateral deafferentation of lumbar spinal cord on HL-PA and hindlimb asymmetry in stretching resistance (SR) induced by L-UBI and R-UBI in rats with completely transected spinal cord. (**A**) Experimental design. The spinal cord was transected at the C6-7 level that was followed by L-UBI (n = 8), R-UBI (n = 11), or sham surgery (Sh; n = 7). The asymmetries were analyzed three hours after UBI or sham surgery (Post) and in the same rats after bilateral rhizotomy performed from the L1 to S2 spinal levels (designated as Rhz). (**B**) The HL-PA size (PAS) in millimeters (mm) and (**C**) the probability to develop HL-PA (P_A_) above 1 mm threshold (denoted by dotted vertical lines). (**D,E**) Contrast between the UBI and sham surgery groups in the PAS and P_A_ before (Post) and after rhizotomy (Rhz). (**F,G**) The effects of rhizotomy on differences in the MPA (or P_A_) between L-UBI, R-UBI and sham surgery (Sh) analyzed as contrast of contrasts between i) L-UBI and sham surgery: ΔΔMPA (or ΔΔP_A_) = [(L-UBI _Post_ – Sh _Post_) – (L-UBI _Rhz_ – Sh _Rhz_)]; ii) R-UBI and sham surgery: ΔΔMPA (or ΔΔP_A_) = [(R-UBI _Post_ – Sh _Post_) – (R-UBI _Rhz_ – Sh _Rhz_)]; and iii) R-UBI and L-UBI sham surgery: ΔΔMPA (or ΔΔP_A_) = [(R-UBI _Post_ – L-UBI _Post_) – (R-UBI _Rhz_ – L-UBI _Rhz_)]. (**H**) Representative traces of the stretching force recorded from the left and right hindlimbs of rats with transected spinal cord and UBI or sham surgery after rhizotomy. (**I**) Differences in stretching force calculated between the left and right hindlimbs as ΔW_LR_ analyzed after rhizotomy in spinalized rats with UBI or sham surgery. (**J**) Difference (contrast) between the UBI and sham surgery groups in ΔW_LR_ analyzed 3 h after rhizotomy. (**K**) The effects of rhizotomy on differences in ΔW_CI_ = (W_Contra_ – W_Ipsi_) analyzed as contrast of contrasts ΔΔ between i) L-UBI and sham surgery: [(L-UBI _Post_ – Sh _Post_) – (L-UBI _Rhz_ – Sh _Rhz_)]; ii) R-UBI and sham surgery: [(R-UBI _Post_ – Sh _Post_) – (R-UBI _Rhz_ – Sh _Rhz_)]; and iii) R-UBI and L-UBI: [(R-UBI _Post_ – L-UBI _Post_) – (R-UBI _Rhz_ – L-UBI _Rhz_)]. The PAS, P_A_, MPA, ΔW_LR_, ΔW_CI_ and contrasts are plotted as median (black circles), 95% HPDC intervals (black lines), and posterior density (colored distribution) from Bayesian regression. Effects on asymmetry and differences between the groups: 95% HPDC intervals did not include zero, and adjusted P-values were ≤ 0.05. Adjusted P is shown for differences identified by Bayesian regression. Crosses in (**B**) and (**I**) denote the PAS and ΔW_LR_ values for individual rats, respectively. **Source data:** The EXCEL source data file “masterfile-210807.xlsx” and source data folder “/HL-PA/data/SF/”.

Representative traces of the stretching force recorded from the left and right hindlimbs of rats with transected cervical spinal cord before and 3 h after UBI or sham operation are shown on **Figure 1H**. Force to stretch hindlimbs increased during stretching starting from zero. No differences in the stretching force were evident between the contra and ipsilesional hindlimbs in rats analyzed before sham surgery and UBI, and those after sham surgery (**Figure 1H,I**). The left and right UBI induced marked increase in the stretching resistance of the contralesional limb. Differences in the work applied to stretch the left and right hindlimbs ΔW_LR_ and AI_LR_ were highly significant after left and right UBI while sham surgery did not produce the asymmetry (**Figure 1I**; **Figure 1—figure supplement 3B**). Both the left and right UBI groups significantly differed from the sham group in the ΔW_LR_ (**Figure 1J**) and AI_LR_ (**Figure 1—figure supplement 3C**). The ΔW_CI_ strongly correlated with the MPA (**Figure 1K**; ΔW_CI_ *vs*. the MPA). Thus in rats with completely transected cervical spinal cord both the left and right UBI produced robust and highly significant increase in the musculo-articular resistance to stretching of the contralesional vs. ipsilesional hindlimbs. We concluded that the UBI-induced asymmetries were not mediated by the sympathetic system or descending neural tracts, and that the endocrine pathway may be only an option for the left-right side-specific signaling from injured brain in these experiments.

### Effects of bilateral deafferentation of lumbar spinal segments on HL-PA induced by the L-UBI or R-UBI

#### HL-PA formation

HL-PA may be developed due to activation of spinal reflexes or changes in the efferent spinal circuits (Zhang et al., 2020). We next sought to determine whether afferent somatosensory input is required for persistence of HL-PA induced by the UBI through a humoral pathway. The left or right UBI was performed in rats with completely transected cervical spinal cord, and the effects of bilateral rhizotomy of the dorsal roots from the L1 to S2 levels on HL-PA were analyzed (**Figure 2**). In rats with left UBI, the PAS and P_A_ were strongly, 3.0- and 3.5-fold, respectively, reduced after rhizotomy (**Figure 2B,C**). In contrast, in the right-side UBI rats, the PAS and P_A_ demonstrated only small, approximately 1.3-fold decrease after deafferentation. No signs of the asymmetry were revealed in the sham surgery rats after rhizotomy.

Contrast in both the PAS and P_A_ was strong and highly significant between the UBI groups and sham surgery group before the rhizotomy, and between the right UBI and sham surgery group after rhizotomy (**Figure 2D,E**). In opposite, the left UBI – sham surgery contrast was negligible in rats analyzed after deafferentation. Relative impact of rhizotomy (before vs. after it) on the effects of left and right UBI (UBI vs. sham surgery) was analyzed as contrast of contrasts (**Figure 2F,G**). Contrast in each the MPA and P_A_ between the animal groups (left UBI vs. sham surgery; right UBI vs. sham surgery; and left UBI vs. right UBI) was compared between two time points that were before vs. after rhizotomy. Contrast was high and significant for comparison of the left UBI group with sham surgery group, whereas it was much smaller when the right UBI rats were compared with sham rats. Contrast of contrasts in both the MPA and P_A_ for the left UBI vs. right UBI was strong and significant.

#### Stretching resistance analysis

The stretching resistance of the contra- and ipsilesional hindlimbs in rats with transected cervical spinal cords was analyzed before and after bilateral rhizotomy that was performed 3 hours after UBI or sham surgery (**Figure 2H-K**; **Figure 2—figure supplement 1**). In rats with left UBI, rhizotomy abolished the differences between the hindlimbs in the resistance. In contrast, the asymmetry was only slightly decreased after rhizotomy in rats with the right UBI. No rhizotomy effects on the left-right differences were evident in rats with sham surgery. Contrast in both the ΔW_LR_ and ΔAI_LR_ between UBI groups and sham surgery group was strong and highly significant before rhizotomy, while after it no differences in the left UBI group were evident (**Figure 2J**; **Figure 2—figure supplement 1**). Contrasts between the right UBI – sham surgery groups remained strong and significant after rhizotomy.

Impact of rhizotomy (contrast: before vs. after it) on the effects of left and right UBI (contrast: UBI vs. sham surgery) was compared as contrast of contrasts in both the ΔW_CI_ and ΔAI_CI_ (**Figure 2K**; **Figure 2—figure supplement 1**). Contrast of the left UBI group vs. sham group was high and significant, while that of the right UBI rats vs. sham group was noticeably smaller. Contrast of contrasts for the left UBI vs. right UBI was strong and highly significant.

Thus, the HL-PA and stretching resistance data corroborate, and demonstrate that the effects of the left-side and right-side UBI mediated through humoral pathway are differentially sensitive to bilateral deafferentation. The effects of the left UBI may depend on afferent somatosensory input, while those of the right-side UBI on activity of efferent motoneurons signaling to hindlimb muscles.

### Effect of opioid antagonists on hindlimb asymmetry in posture and stretching resistance induced by the left and right UBI

We previously demonstrated that naloxone, a nonselective opioid antagonist, blocked the asymmetric effects of the left UBI on hindlimb posture in rats with transected spinal cords, and that the selective opioid antagonists differentially inhibited formation of HL-PA after the left and right UBI in rats with intact spinal cords (Lukoyanov et al., 2021; Watanabe et al., 2021). We here examined if the UBI effects mediated through the humoral pathway are controlled by the opioid system, and if this control is side- and receptor subtype-specific.

The effects of the µ-, δ- and κ-opioid antagonists β-Funaltrexamine (FNA), naltrindole (NTI) and nor-Binaltorphimine (BNI), respectively, and naloxone on the asymmetry in hindlimb posture and stretching resistance were compared between rats with the left and right-side UBI (**Figure 3**; **Figure 3—figure supplement 1**). BNI and FNA are long acting antagonists that selectively block these receptor subtypes, and require approximately 24 h after administration to do so (Horan, Taylor, Yamamura, & Porreca, 1992; Patkar et al., 2013; Petrillo et al., 2003; Rutten, Schroder, Christoph, Koch, & Tzschentke, 2018). These antagonists were administered to the rats 24 h before the surgeries. NTI and naloxone were administered 3-4 h after UBI to rats that displayed HL-PA with the MPA > 1.5 mm before the injection, and then HL-PA was analyzed 1 h later.

**Figure 3.**
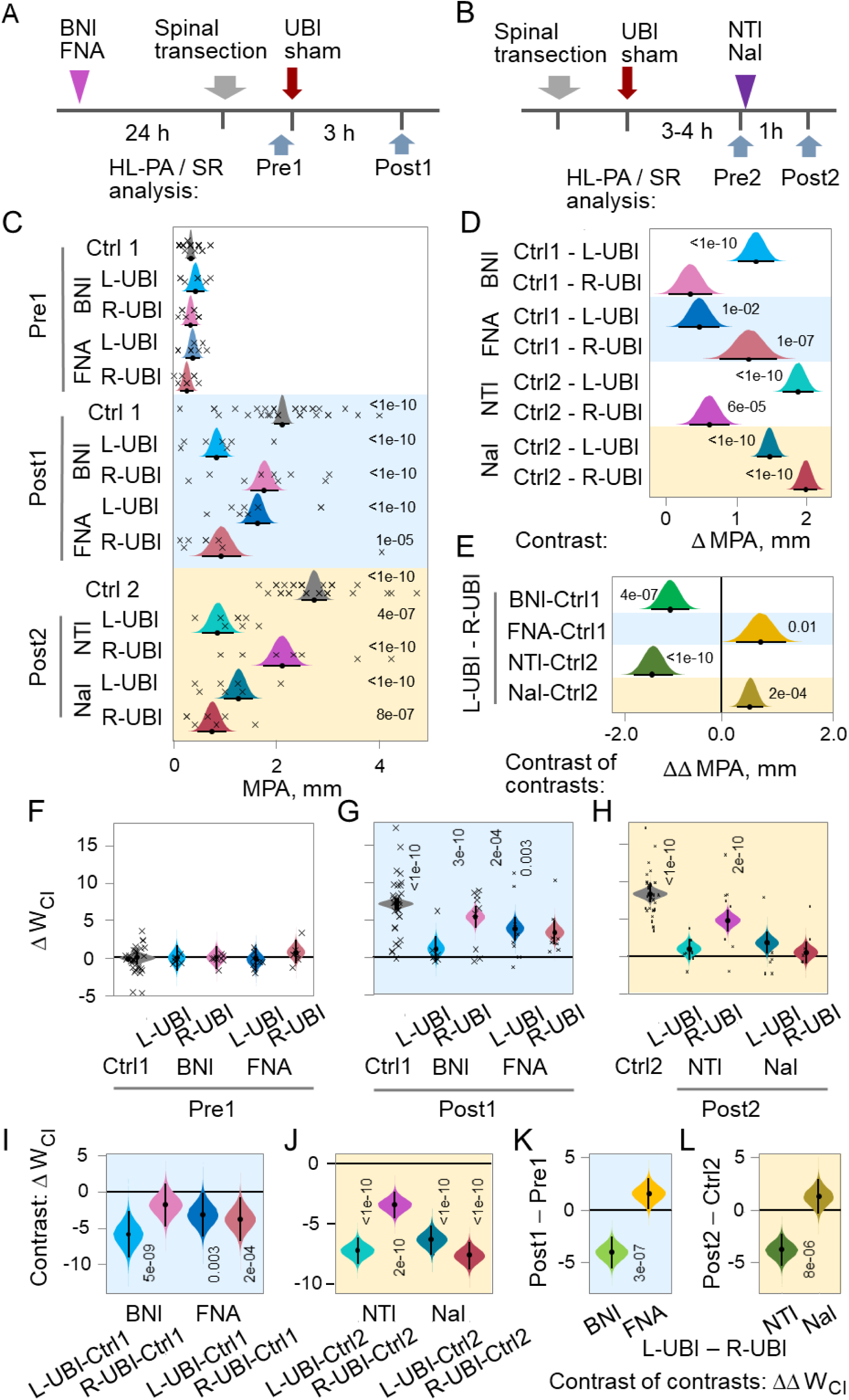
Effects of nor-Binaltorphimine (BNI), β-Funaltrexamine (FNA) and naltrindole (NTI), the selective κ-, µ- and δ-opioid antagonists, respectively, and naloxone (Nal), the general opioid antagonist, on HL-PA and hindlimb asymmetry in stretching resistance (SR) induced by L-UBI and R-UBI in rats with completely transected spinal cord. (**A**) Experimental design 1. The spinal cord was transected at the C6-7 level and then the brain was injured. BNI and FNA were administered 24 h before the surgeries. The asymmetries were analyzed after the transection before the UBI (designated as Pre1), and then 3 h after the UBI (Post1). Control group consisted of rats with transected spinal cords and UBI that were not treated with drugs (Ctrl 1; n = 30). (**B**) Experimental design 2. The asymmetries were assessed 3-4 h after UBI in rats with transected spinal cords (Pre2). The rats with the MPA exceeding 1 mm were treated with NTI, Nal or saline and 1 h later the asymmetries were analyzed (Post2). Control group Ctrl2 consisted of rats with MPA > 1.5 mm that were not treated with saline or drugs and analyzed 3-4 h after UBI (Pre1; n = 24), and rats treated with saline 3 h after UBI and analyzed 1 h later at Post2 time point (n = 7). No significant differences in the MPA between saline treated and untreated groups were revealed, and they were combined into the Ctrl2 group (n = 31). Both the Ctrl1 control group for BNI and FNA, and Ctrl2 control group for NTI and Nal, were composed of rats with the L-UBI and R-UBI. No statistically significant differences in both the MPA and asymmetry in stretching resistance ΔW_CI_ between the L-UBI and R-UBI subgroups of each control group were revealed, and these subgroups were combined into respective control group for statistical analysis. (**C**) The antagonist effects on the MPA. (**D**) Contrasts in the MPA between the respective control groups and the groups treated with antagonists. (**E**) Difference in the effects of antagonists on the MPA between the L-UBI and R-UBI groups analyzed as contrast of contrasts: ΔΔMPA = [(L-UBI _Post1_ – Ctrl1) – (R-UBI _Post1_ – Ctrl1)] for BNI and FNA; and ΔΔMPA = [(L-UBI _Post2_ – Ctrl2) – (R-UBI _Post2_ – Ctrl2)] for NTI and Nal. (**F-H**) Differences in stretching force between the contra- and ipsilesional hindlimbs ΔW_CI_ in gm × mm. (**I,J**) Contrasts in the ΔW_CI_ between the UBI groups treated with the antagonists and respective control groups. (**K,L**) Difference in the effects of antagonists on the ΔW_CI_ between the L-UBI and R-UBI groups analyzed as contrast of contrasts; **K**: ΔΔW_CI_ = [(L-UBI _Post1_ – Pre1) – (R-UBI _Post1_ – Pre1)] and **L**: ΔΔW_CI_ = [(L-UBI _Post2_ – Ctrl2) – (R-UBI _Post2_ – Ctrl2)]. Crosses denote the MPA and ΔW_CI_ values for individual rats. The MPA, ΔW_CI_ and contrasts are plotted as median (black circles), 95% HPDC intervals (black lines), and posterior density (colored distribution) from Bayesian regression. Effects on asymmetry and differences between the groups: 95% HPDC intervals did not include zero, and adjusted P-values were ≤ 0.05. Adjusted P is shown for differences identified by Bayesian regression. The number of rats in the groups is given in Figure 3**—figure supplement 1**. **Source data:** The EXCEL source data file “SDU-RDPA-Stat_v2.xlsx” and source data folder /HL-PA-opioid-antagonists/data/SF/”.

In the left UBI group, a marked decrease in the MPA was induced by NTI and BNI (3.2- and 2.5-fold, respectively) while no substantial effects of FNA on HL-PA were evident (**Figure 3C,D**). In contrast, in the right-side UBI group administration of FNA, but not NTI and BNI, resulted in substantial MPA reduction (2.3-fold). Naloxone inhibited the effects of both the left- and right side injuries (2.2 and 3.7-fold, respectively). The effects of the antagonists were significantly different between the left and right UBI groups (**Figure 3E**). NTI and BNI preferentially inhibited formation of the right hindlimb flexion, whereas in contrast, FNA and naloxone stronger inhibited flexion on the left side.

Administration of BNI and NTI strongly decreased the contra-ipsilesional hindlimb asymmetry in stretching resistance ΔW_CI_ after the left UBI, while their effects were minor in rats with the right UBI (**Figure 3F-L**). The FNA effects were less pronounced (**Figure 3G,I,K**). Naloxone substantially reduced the ΔW_CI_ in rats either with the left or right UBI (**Figure 3H,J,L**). Analysis of contrast of contrasts revealed significant differences between the left and right UBI groups in the effects of BNI and NTI (**Figure 3K,L**). The effect of FNA on the ΔW_CI_ after the right UBI slightly exceed that after the left side injury (**Figure 3K**). Thus, the HL-PA and stretching resistance data are in general agreement; the effects of the left and right UBI were differently inhibited by the opioid antagonists suggesting that the left and right T-NES counterparts are differentially controlled by opioid mechanisms.

### Gene expression and co-expression patterns: the UBI effects

Signals from the injured hemisphere may be encoded into left-right side-specific hormonal messages in the hypothalamic-pituitary system (Lukoyanov et al., 2021). After being released these messages target the lumbar spinal cord that may result in a coordination of gene expression across these neuroendocrine and motor regions. The prerequisite for the left-right side-specific encoding of neurohormonal messages may be an asymmetrical organization of hypothalamic neurosecretory circuits including their gene expression profiles, and the side-specific responsiveness of these circuits to a unilateral impact. We previously reported that the “decoding” lumbar spinal cord is characterized by asymmetric gene expression, and that the UBI produced the ipsi-contralesional side-specific changes in gene expression and gene-gene co-expression patterns in rats with complete spinal cord transection (Kononenko et al., 2017; Lukoyanov et al., 2021; Watanabe et al., 2021).

We here examined if the UBI targets the hypothalamus and pituitary gland as the “encoding” areas by analysis of gene expression as a readout; if the UBI effects are the ipsi- or contralesional; and if expression of neurohormonal and neuroplasticity-related genes is lateralized in this area. To reveal regulatory humoral interactions between the “encoding” hypothalamus and “decoding” spinal cord, we then characterized gene-gene co-expression patterns between these regions in rats with complete spinal cord transection along with their perturbations produced by UBI.

#### The hypothalamic-pituitary system

We reasoned that the “encoding” system that mediates the neuroendocrine UBI effects in the hypothalamus involves genes of the releasing and inhibitory hormones (*Crh, Ghrh, Gnrh1, Sst* and *Trh*); neuropeptides and their receptors genes (*Avp, Avpr1a, Nts, Penk, Pdyn, Pomc, Oprm1, Oprk1, Oprd1* and *Oxt*; **Figure 4—figure supplement 1-3**), along with neuroplasticity-related genes coding for regulators of axonal sprouting, synapse formation, neuronal survival and neuroinflammation (*Arc*, *Bdnf*, *Dlg4*, *Homer-1*, *Gap43*, *Syt4* and *Tgfb1*), transcriptional regulators of synaptic plasticity (*cFos*, *Egr1* and *Nfkbia*), and essential components of the glutamate system critical for neuroplasticity (*GluR1*, *Grin2a* and *Grin2b*) (**Figure 4—figure supplement 4**; for detailed description, see “Materials and methods”).

**Figure 4.**
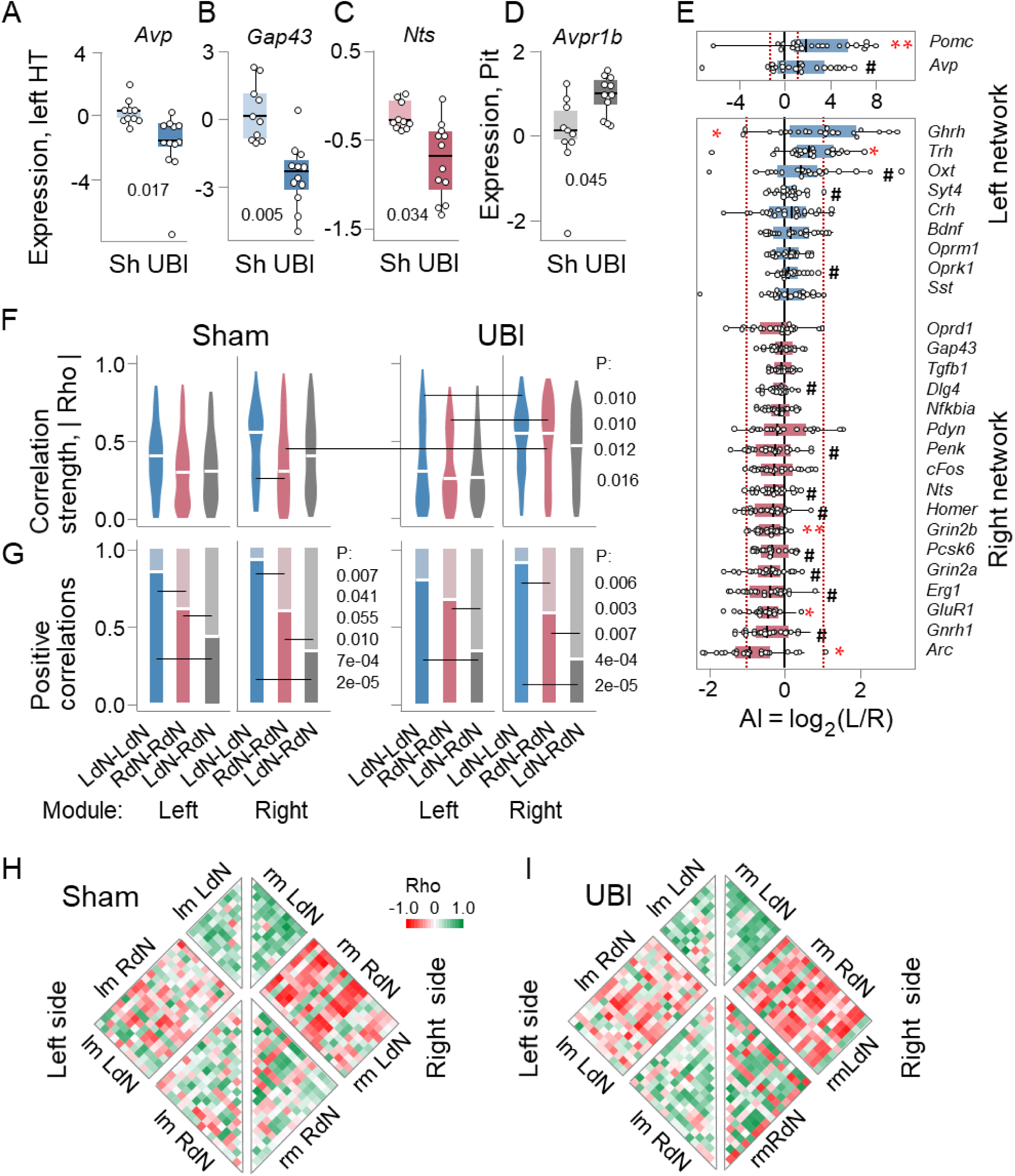
The UBI effects on gene expression patterns in the hypothalamus and pituitary gland. Analysis of the left (LdN) and right (RdN) side dominant gene regulatory networks in the hypothalamus. Gene expression was analyzed in the left and right hypothalamus (28 genes) (**A-C**) and the pituitary gland (17 genes) (**D**) collected 3 h after left sham surgery (Sh: n = 11 rats) or left UBI (n = 12 rats). The data expression levels in the log_2_ scale and the Bonferroni adjusted P values determined by Mann–Whitney test are shown. (**E**) The asymmetry index AI = log_2_[L/R], where L and R are the median expression levels in the left and right hypothalamus. Sham surgery and UBI groups were combined (n = 22) because there were no differences in the AI between these groups. Wilcoxon signed-rank test followed by Bonferroni multiple testing correction: *, P adj < 0.05; **, P adj < 0.01; #, P ≤ 0.05 (not adjusted). Light blue boxes denote genes with AI > 0, and pink boxes with AI < 0 that were defined as the LdN and RdN. In (**A**-**E**), data are presented as boxplots with median and hinges representing the first and third quartiles, and whiskers extending from the hinge to the highest/lowest value that lies within the 1.5 interquartile range of the hinge. (**F,G**) Patterns of intra-modular correlations within each the LdN (LdN-LdN) and RdN (RdN-RdN) and between them (LdN-RdN). (**F**) The coordination strength (absolute Rho value averaged across pairwise correlations) and (**G**) the proportion of positive correlations for the left and right modules of the sham surgery and UBI groups. The correlation patterns were compared within each the left and right module, and between UBI and sham surgery groups (for details, see Figure 4**—figure supplement 10**). (**H,I**) Heatmaps for Spearman’s rank coefficients for pairwise gene-gene correlations in the left- and right modules of sham surgery and UBI groups. P values were determined by permutation testing with Benjamini-Hochberg family-wise multiple test correction, and are shown for the median AI of the combined sham surgery and UBI group. P values computed for the variant 1 are shown for the contrasts that are significant (P ≤ 0.05) either after the correction for all three categorization variants, or for two of them while in the third variant P was < 0.05 and < 0.10 before and after the correction, respectively (Figure 4**—figure supplement 9**). **Source data:** The EXCEL source data files “Hypoth_SO_UBI.xlsx; RD Hypophis_ Master file.xlsx; Table III-S6 23 05 10.xlsx; raw_groups.xlsx”.

Expression of the *Avp* (fold change (FC)=3.46x), *Gap43* (FC= 1.18x) and *Nts* (FC=1.32x) genes were significantly affected by the left UBI in the left hypothalamus (**Figure 4A-C**). The UBI induced effects were nominally significant for the *Crh* (FC=1.76x), *Sst* (FC=1.59x), *Bdnf* (FC=1.25x), *Syt4* (FC=1.22x), *Pomc* (FC=2.49x) and *Ghrh* (FC=1.63x) genes (**Figure 4—figure supplement 5**). The expression levels of these genes were lower in the left hypothalamus in the UBI group compared to sham surgery group. No significant and nominally significant differences were revealed in the right hypothalamus, while the UBI-induced changes were consistent between the left and right sides in their magnitude. Pearson and Spearman’s rank correlation coefficients between log-scaled FCs induced by UBI in the left and the right hypothalamus were equal to 0.79 (P = 5.4×10^-7^) and 0.48 (P = 0.010), respectively (**Figure 4—figure supplement 6**). Fitting the data with a linear model with an arbitrary intercept (logFC_right_ ≍ *a* logFC_left_ + *b*) resulted in estimates of *a* = 0.64 (95% confidence interval [0.45, 0.83]) that was significantly smaller than 1. Consistently, absolute values of the FC significantly differed between the sides (Wilcoxon signed rank test: P = 0.018). Thus the effects of the left UBI on the left side were significantly greater than on the right side.

No significant differences in AI = log_2_[L/R], where L and R were expression levels in the left and right hypothalamus) were identified between UBI and sham groups, and these group were combined for analysis of lateralization. Comparison of the AI with zero identified three genes (*Pomc*, P = 0.009; *Trh*, P = 0.014; and *Ghrh*, P = 0.014) with higher expression in the left hypothalamus while other three genes (*Grin2b*, P = 0.002; *GluR1*, P = 0.010; and *Arc*, P = 0.010) showed higher expression on the right side (**Figure 4E**). Lateralization was nominally significant for the *Avp*, *Oprk1, Syt4* and *Oxt* genes that showed higher expression in the left hypothalamus, and for the *Grin2a, Homer, Nts, Erg1, Pcsk6, Penk, Gnrh1* and *Dlg4* genes demonstrated higher expression on the right side.

In the pituitary gland, the expression of the hormonal (*Fshb, Cga, Gh1, Lhb, Prl* and *Tshb*), opioid peptides and their receptors (*Oprm1, Oprd1, Oprk1*, *Pdyn, Penk,* and *Pomc*), oxytocin (*Oxt*), Arg-vasopressin and its receptors (*Avp, Avpr1a, Avpr1b* and *Avpr2*) (**Figure 4—figure supplements 2,3,7**) genes were compared between the UBI and sham surgery groups. The expression levels of the *Avpr1b* gene were significantly higher (FC=1.84x) (**Figure 4D**), while those of the *Oxt* (FC = 1.17x) and *Tshb* (FC = 1.63x) genes (**Figure 4—figure supplement 8**) were higher with nominal significance in the UBI group.

Thus in the hypothalamus expression of a subset of the neurohormonal, neuropeptide and neuroplasticity-associated genes was lateralized and affected by UBI on the ipsilesional side. Among genes responded to UBI in the neuroendocrine system were *Avp*, *Avpr1b* and *Pomc* that give rise to Arg-vasopressin, the vasopressin receptor V1B and β-endorphin that mediate the effects of the left UBI on HL-PA through humoral pathway.

#### Gene-gene co-expression within and between the hypothalamus and spinal cord

Analysis of gene-gene co-expression patterns uncover regulatory interactions between tissues and brain areas (Antonucci et al., 2019; Dobrin et al., 2009; Erola, Bjorkegren, & Michoel, 2020; Gerring, Gamazon, Derks, & Major Depressive Disorder Working Group of the Psychiatric Genomics, 2019; Zhang et al., 2020). We here evaluated if such patterns are coordinated across the hypothalamus and spinal cord and their sides, and if this coordination is mediated through humoral pathway and affected by the UBI. To take into account the lateralization factor, we separately analyzed genes with higher expression either on the left or right side, and defined them as the left dominant network (LdN) or right dominant network (RdN) genes. We examined if gene-gene co-expression patterns differ between these networks and for both network between the sides in the hypothalamus and spinal cord; if the patterns are coordinated between these regions; and if the coordination is ipsi- or contralateral and perturbed by a unilateral brain lesion. Gene-gene co-expression was analyzed by pairwise Spearman correlations. The coordination strength and the directions (signs) of interactions in the gene-gene co-expression patterns were assessed as mean of the absolute value of the correlation coefficient Rho and the proportion of positive correlations, respectively.

##### Categorization of genes into the left and right networks

In the hypothalamus and spinal cord separately, genes were categorized into the LdN and RdN based on their AI that was either above or below zero, respectively. Genes expressed on the left or right sides constituted the left (lm) and right (rm) modules of these networks, respectively. To avoid a bias, the categorization was performed in three variants that then were used in the correlation analysis. In the variant 1, the LdN and RdN genes were defined by their median AI in the combined sham surgery and UBI group; in the variant 2, by their median AI in sham surgery group only; and in the variant 3, by their mean AI in the combined sham surgery and UBI group (**Figures 4E** and **5A**; **Figure 4—figure supplement 9; Figure 5—figure supplement 1**). A P value between correlation patterns was determined by the permutation test with Benjamini-Hochberg family-wise multiple test correction. The contrast was defined as significant using a stringent criterion; namely if the P value was ≤ 0.05 for i) all three variants after correction, or for ii) any two of them while for the third variant it was < 0.05 and < 0.10 before and after the correction, respectively. In the hypothalamus, all genes showed stable patterns between the LdN and RdN in three categorization variants besides *Mor* that wobbled. In the spinal cord, five LdN genes and eight RdN genes showed stable patterns across the three variants, while seven genes wobbled between the sides.

**Fig. 5.**
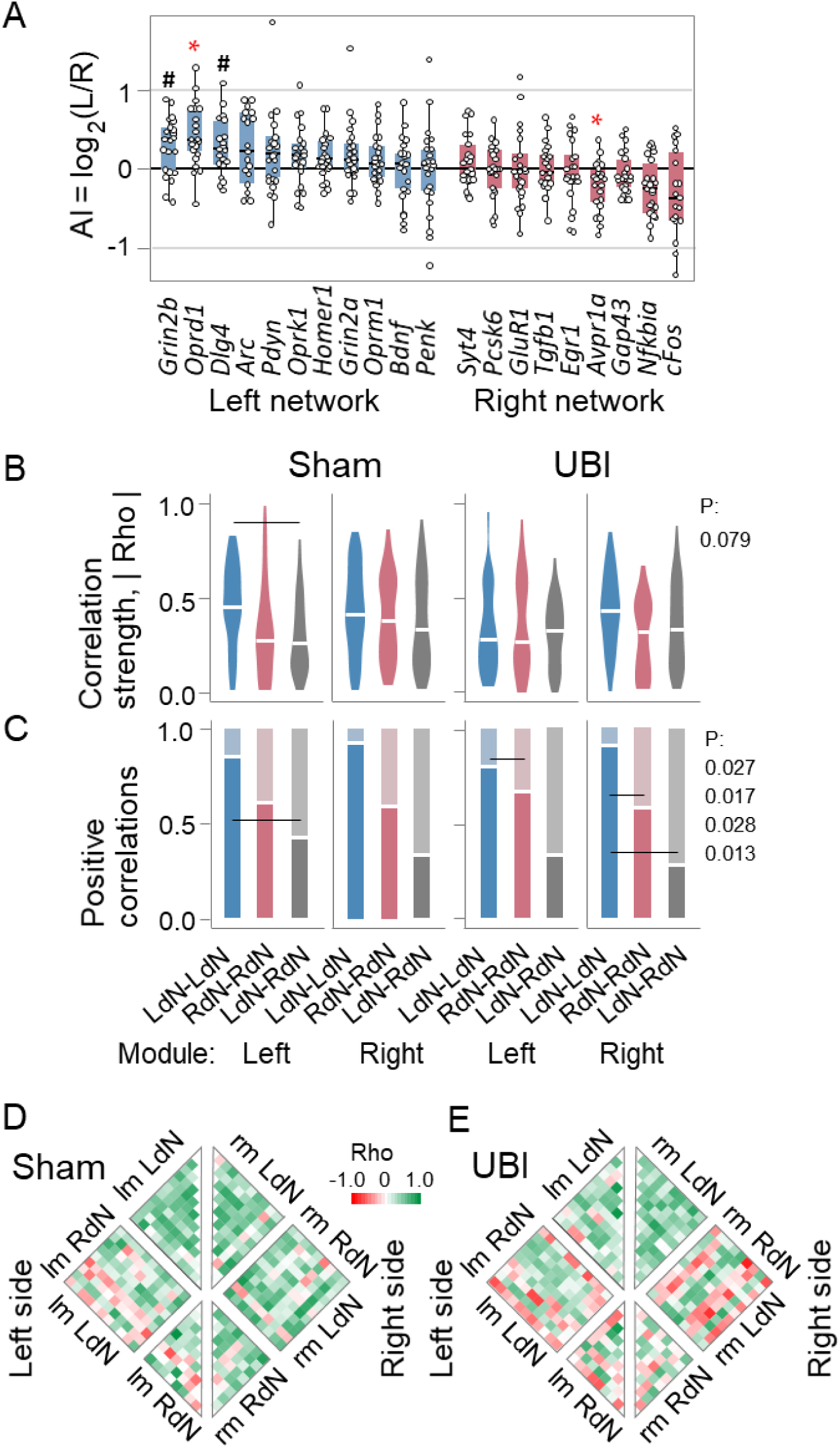
Gene co-expression patterns in the left and right lumbar spinal cord of the sham surgery and UBI rats. Analysis of the LdN and RdN gene networks. Expression of the neurohormonal and neuroplasticity-related genes was analyzed in the left and right halves of the spinal cord isolated 3 h after the left sham surgery (n = 11) or left UBI (n = 12) that were performed in rats with complete spinal cord transection. (**A**) Gene clustering by the AI = log_2_[L/R], where L and R are the median expression levels in the left and right spinal cord into the LdN (AI > 0) and RdN (AI < 0). The median AI is shown for the combined sham surgery and the left UBI group because there were no differences in the AI between these groups (n = 22); for details, see **Figure 5—figure supplement 1**). Wilcoxon signed-rank test followed by Bonferroni multiple testing correction: *, P adj < 0.05; **, P adj < 0.01; #, P ≤ 0.05 (not adjusted). Data are presented as boxplots with median and hinges representing the first and third quartiles, and whiskers extending from the hinge to the highest/lowest value that lies within the 1.5 interquartile range of the hinge. (**B,C**) Patterns of intra-modular correlations within each the LdN (LdN-LdN) and RdN (RdN-RdN) and between them (LdN-RdN). (**B**) The coordination strength (absolute Rho value averaged across pairwise correlations) and (**C**) the proportion of positive correlations for the left and right modules of the sham surgery and UBI groups. The correlation patterns were compared within each the left and right module, and between UBI and sham surgery groups. (**H,I**) Heatmaps for Spearman’s rank coefficients for pairwise gene-gene correlations in the left- and right modules of sham surgery and UBI groups. P values were determined by permutation testing with Benjamini-Hochberg family-wise multiple test correction, and are shown for the median AI of the combined sham surgery and UBI group. P values computed for the variant 1 are shown for the contrasts that are significant (P ≤ 0.05) either after the correction for all three categorization variants, or for two of them while in the third variant P was < 0.05 and < 0.10 before and after the correction, respectively (**Figure 5— figure supplement 1**). **Source data:** The EXCEL source data file “SpinalC_SO_UBI_Ctrl_RD_DD.xlsx; Table III-S6 23 05 10.xlsx; raw_groups.xlsx”.

##### Correlation patterns in the hypothalamus and spinal cord

We compared the LdN and RdN in their coordination strength and the proportion of positive correlations for intra-modular correlations in both the left (lmLdN-lmLdN, lmRdN-lmRdN and lmLdN-lmRdN) and right (rmLdN-rmLdN, rmRdN-rmRdN and rmLdN-rmRdN) modules; these intra-modular correlations between the modules; and inter-modular correlations lmLdN-rmLdN and lmRdN-rmRdN (**Figures 4** and **5**, **Figure 4—Supplements 10, 11, Figure 5—Supplements 1,2**). The patterns were compared between sham surgery and UBI groups.

In the hypothalamus, the permutation test revealed significant differences in the coordination strength between the LdN and RdN (LdN-LdN > RdN-RdN) in the right module of sham surgery rats, and between the left and right modules for the both networks (lmLdN-lmLdN < rmLdN-rmLdN; lmRdN-lmRdN < rmRdN-rmRdN) in the UBI rats (**Figure 4C**). UBI resulted in elevation of the RdN-RdN strength on the contralesional side.

Robust and significant differences in the proportion of positive correlations in the hypothalamus were evident between correlations that were internal for the LdN and RdN (sham surgery: lmLdN-lmLdN > lmRdN-lmRdN; rmLdN-rmLdN > rmRdN-rmRdN; UBI: rmLdN-rmLdN > rmRdN-rmRdN), and between these and mixed (LdN-RdN) correlations (sham surgery: lmLdN-lmLdN > lmLdN-lmRdN; lmRdN-lmRdN > lmLdN-lmRdN; rmLdN-rmLdN > rmLdN-rmRdN; rmRdN-rmRdN > rmLdN-rmRdN; UBI: lmLdN-lmLdN > lmLdN-lmRdN; lmRdN-lmRdN > lmLdN-lmRdN; rmLdN-rmLdN > rmLdN-rmRdN; rmRdN-rmRdN > rmLdN-rmRdN) (**Figure 4G**). Furthermore, the proportion was significantly larger in the inter-modular LdN (lmLdN-rmLdN) vs. RdN (lmRdN-rmRdN) correlations (**Figure 4G** and **Figure 4—figure supplement 11**).

In the spinal cord, differences in the coordination strength were significant between the LdN and mixed correlation patterns in the left module of sham surgery rats (LdN-LdN > LdN-RdN) (**Figure 5B**). The pattern of differences in the proportion of positive correlations was generally similar with that in the hypothalamus, however less contrasts were significant (**Figure 5C**). The proportion in the LdN was larger than that in the RdN (UBI: lmLdN-lmLdN > lmRdN-lmRdN; rmLdN-rmLdN > rmRdN-rmRdN), and the mixed correlations (sham surgery: lmLdN-lmLdN > lmLdN-lmRdN; UBI: rmLdN-rmLdN > rmLdN-rmRdN). Furthermore, the proportion was larger in the inter-modular lmLdN-rmLdN correlations in the sham surgery group vs. UBI group (**Figure 5—figure supplement 2**).

In summary, the LdN and RdN were strongly and significantly different between each other in the coordination strength and the proportion of positive correlations both in the hypothalamus and spinal cord. The UBI produced contrasting effects on the left and right hypothalamic modules in the coordination strength that was elevated for both gene networks on the contralesional side. In the significant contrasts, both the coordination and the proportion were higher for the LdN compared to the RdN. Strikingly, correlations were largely positive within each network, and in contrast, were mostly negative between them that suggests positive regulatory interactions among the genes in each network and negative regulations between the networks. These differences are clearly seen on heatmaps (**Figures 4H,I and 5D,E**).

##### Coordination of gene expression between the hypothalamus and spinal cord

We next examined if the LdN and RdN are coordinated between the hypothalamus and the lumbar spinal cord through the humoral pathway, and if this crosstalk is perturbed by the unilateral impact (**Figure 6**, **Figure 6—figure supplement 1**). First, the coordination was analysed as the ipsilateral correlations between the left halves (L-Ip) and between the right halves (R-Ip) of these regions for each the LdN and RdN separately (**Figure 6**). The coordination strength in the LdN for the R-Ip pattern was significantly greater than that for i) the LdN for the L-Ip pattern and ii) the RdN R-Ip pattern in sham surgery group, and iii) the LdN R-Ip pattern in the UBI group (**Figure 6B**). The proportion of positive correlations for both the LdN and RdN in sham surgery group was strongly asymmetric: it was much higher on the left *vs.* right side for the LdN, and, in opposite, was much higher on the right *vs.* left side for the RdN. At the same time, the proportion and coordination strength were quite similar between the L-Ip-LdN and R-Ip-RdN patterns, and between the L-Ip-RdN and R-Ip-LdN patterns (**Figure 6B,C**).

**Fig. 6.**
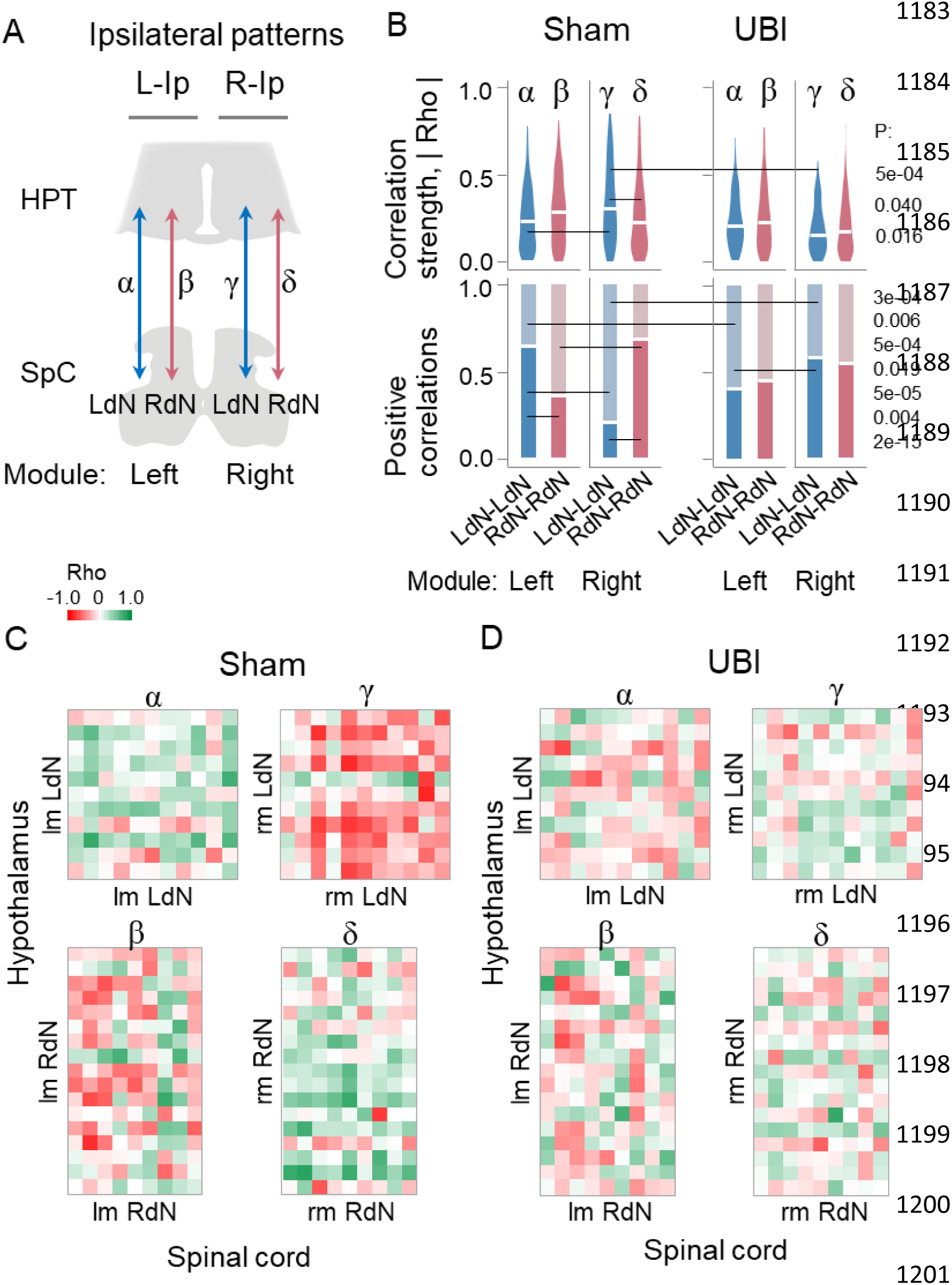
Ipsilateral coordination of the LdN and RdN between the hypothalamus and lumbar spinal cord. The effects of UBI. Experimental design is described in **Figs. 5** and **6**. (**A**) Analyzed patterns of the ipsilateral pairwise gene-gene Spearman rank correlations between the hypothalamus (HPT) and spinal cord (SpC) on the left side (L-Ip: α and β) and right side (R-Ip: γ and δ). (**B**) The coordination strength and the proportion of positive correlations for the correlation patterns depicted in (**A**). The correlation patterns were compared between the LdN and RdN (α *vs*. β; γ *vs*. δ); each of them between the left and right modules (α *vs*. γ; β *vs*. δ), and all the patterns between UBI and sham surgery groups. (**C**,**D**) Heatmaps for Spearman’s rank coefficients for pairwise gene-gene correlations in sham surgery and UBI groups. P values computed for the variant 1 are shown for the contrasts that are significant (P ≤ 0.05) either after the correction for all three categorization variants, or for two of them while in the third variant P was < 0.05 and < 0.10 before and after the correction, respectively (**Figure 4—figure supplement 9 ; Figure 5—figure supplement 1**). **Source data:** The EXCEL source data file “Hypoth_SO_UBI.xlsx; SpinalC_SO_UBI_Ctrl_RD_DD.xlsx; Table III-S6 23 05 10.xlsx; raw_groups.xlsx”.

The UBI impaired the hypothalamic-spinal cord crosstalk (**Figure 6B,D**). Notably, only the LdN was affected by lesion of the left somatosensory cortex. The left UBI resulted in strong decrease of the coordination strength and elevation of the proportion in the R-Ip LdN patterns (on the contralesional, right side); while in the L-Ip LdN patterns the proportion was strongly decreased. Furthermore, in the UBI group the proportion was higher for the R-Ip LdN pattern than for the L-Ip LdN pattern. Thus, the proportion of positive correlations was higher than random value for the L-Ip pattern and lower it for the R-Ip pattern in sham surgery rats, while this pattern was reversed after the UBI.

Analysis of the contralateral (diagonal) correlations between the left hypothalamus and right spinal cord, and between the right hypothalamus and left spinal cord for each the LdN and RdN did not reveal significant patterns and UBI effects (**Figure 6—figure supplement 1**). In conclusion, the correlation analysis revealed the robust side-specific ipsilateral patterns in the coordination of gene expression between the hypothalamus and spinal cord that differed between the LdN and RdN, and strong perturbations in these patterns by the unilateral ablation injury in animals with transected spinal cords. The findings suggest the functional link between these two regions that is the left-right side-specific and mediated by the endocrine signaling.

## DISCUSSION

### The left-right side-specific humoral signaling from the brain to spinal cord: an alternative to neural pathways

Formation of HL-PA and asymmetry in hindlimb reflexes in rats with transected spinal cords was the basis for the hypothesis that the contralateral effects of unilateral brain lesions are mediated through humoral pathway (Bakalkin, 2022; Lukoyanov et al., 2021). However, signaling from the brain to the lumbar spinal cord through the paravertebral sympathetic chain was not excluded because the spinal transection was performed at the T2-T3 level and left intact the neural connections between the brain and the superior preganglionic neurons. Activity of the sympathetic preganglionic neurons located in the upper thoracic and lumbar segments is coordinated at a supraspinal (medullary) level (Farmer et al., 2019) and muscle sympathetic nerve activity is controlled by central commands (Boulton, Taylor, Green, & Macefield, 2021). Furthermore, the spinal somato-sympathetic nerve reflexes may contribute to the maintenance of muscle contractile force both before and after spinal cord transection (Hotta et al., 2021). In the present study, the supraspinal part of the CNS was fully disconnected from the preganglionic spinal neurons by spinal cord transection at the C6-C7 level that was rostral to the thoracic preganglionic sympathetic neurons. Despite of the transection, the UBI induced asymmetric hindlimb responses. Thus the hypothetical mechanism of the brain injury-induced HL-PA formation mediated through the paravertebral sympathetic ganglia and descending neural pathways was ruled out that provided an unequivocal evidence for the role of the left-right side specific endocrine signaling.

### The bipartite T-NES: intrinsic neurohormonal and neural asymmetry

The UBI-induced signaling is binary, either left or right sided. This could determine the bipartite structure of the T-NES that by encoding and decoding hormonal messages may selectively propagate the effects of either left or right brain lesion (**Figure 7A,B**). The bipartite structure is supported by the findings that the left and right side-specific T-NES functions are differentially affected by selective opioid antagonists. The δ- and κ-antagonists NTI and BNI, respectively, inhibited HL-PA after the left UBI but not after lesion to the right hemisphere, whereas µ antagonist FNA interfered with the effects of the right side injury (**Figure 7A,B**). To note, the opioid antagonists differently blocked the effects of the left and right brain injury in rats with intact spinal cords (Watanabe et al., 2021). The preferential side of inhibition by the δ- and µ-antagonists was the same in rats with intact and completely transected spinal cords. However, the side affected by the κ-antagonist BNI was different between these animal groups. Thus the opioid mechanisms through different receptor subtypes may control a flexibility of outcome and enable synergy or antagonism of the neural and endocrine signaling pathways.

**Fig. 7.**
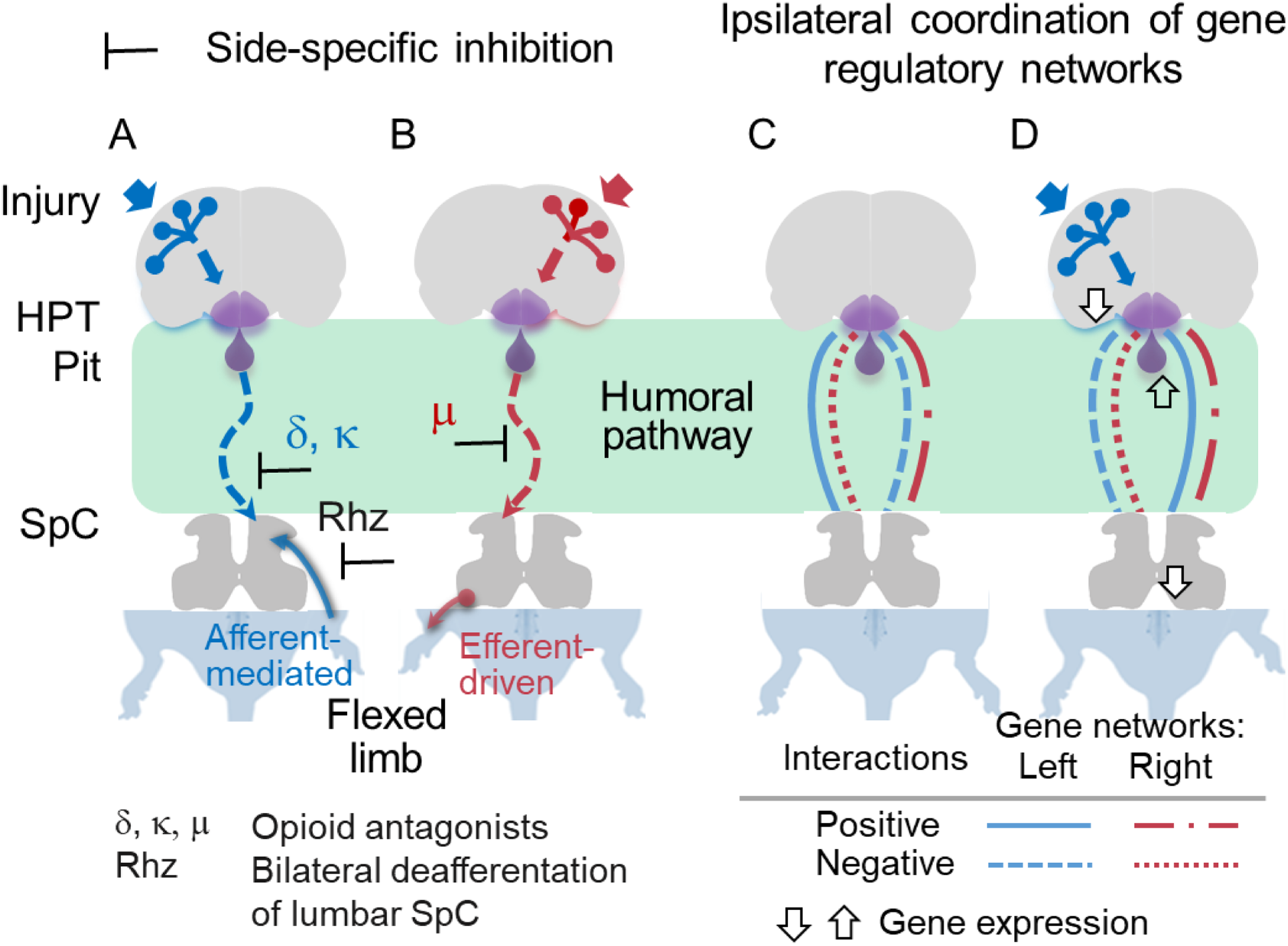
A model of the bipartite asymmetric T-NES that enables the left-right side-specific signaling from the brain to the lumbar spinal cord (SpC) through the humoral pathway. The left (**A**) and right (**B**) T-NES counterparts mediate the contralateral effects of the left-side and right-side brain injury. A unilateral brain lesion stimulates the release of side-specific neurohormones from the hypothalamus (HPT) and pituitary gland (Pit) into the blood, they bind to neuronal receptors that are lateralized in the spinal cord (Kononenko et al., 2017; Watanabe et al., 2021) or peripheral neuronal endings, and induce contralateral responses, e.g., hindlimb flexion. The δ- and κ-opioid receptors may control signaling from the brain injured on the left side, whereas the µ-opioid receptors that after the right UBI. The endogenous opioid peptides could differentially convey signals from the left and right hemispheres through the humoral pathway or control their processing in the hypothalamus or spinal cord. The left and right T-NES counterparts are not mirror symmetric to each other in their neural mechanisms, while produce overall symmetric functional responses, e.g., flexion of the right and left hindlimbs, respectively. The effects of the left T-NES but not right T-NES may require an afferent input and depend on spinal reflexes. The right T-NES effects may develop through activation of efferent neurocircuits or changes in neuromuscular system. (**C,D**) The T-NES-mediated ipsilateral crosstalk between the hypothalamic and lumbar spinal cord gene-regulatory networks, and its reorganization in response to a unilateral brain lesion. Ipsilateral interactions between the hypothalamus and spinal cord are depicted for the left and right gene regulatory networks as positive or negative according to the proportion of positive correlations in each network, that is higher or lower than 0.5, respectively. Arrows show the direction of changes in gene expression induced by the left UBI. The ipsilateral correlations significantly differ between the left and right networks on each body side and for both networks between the sides. Only interactions of the left networks were significantly perturbed by the left UBI. The patterns of interactions were similar or almost mirror symmetric (allo-symmetric) for the left networks on the left side and the right networks on the right side, and for the right networks on the left side and the left networks on the right side. The diagonal (contralateral) inter-regional interactions were not significant.

Translation of the T-NES humoral messages into the left-right side-specific hindlimb responses may occur in lumbar neural circuits or at peripheral nerve endings. We previously demonstrated that opioid peptides, synthetic opioids and Arg-vasopressin may induce HL-PA after their systemic or intrathecal administration into rats with intact brain but completely transected spinal cords (Bakalkin, Iarygin, Trushina, Titov, & Smirnov, 1980; Bakalkin & Kobylyansky, 1989; Bakalkin, Kobylyansky, Nagornaya, Yarygin, & Titov, 1986; Chazov et al., 1981; Lukoyanov et al., 2021; Watanabe et al., 2020). The striking finding was that the side of the flexed limb depended on the compound administered. Met-enkephalin, the μ/δ-opioid ligand and dynorphin, bremazocine and U-50,488, κ-opioid agonists, induced flexion of the left hindlimb. In contrast, the δ-agonist Leu-enkephalin, β-endorphin and Arg-vasopressin cause the right limb to flex (Bakalkin et al., 1980; Chazov et al., 1981; Klement’ev, Molokoedov, Bushuev, Danilovskii, & Sepetov, 1986; Lukoyanov et al., 2021). After brain injury, these neurohormones may be released from the endocrine glands into the bloodstream and induce side-specific effects through their receptors lateralized in the spinal cord or peripheral terminals of afferent and efferent neurons. In the cervical and lumbar spinal cord, the expression of the opioid receptors is lateralized to the left, and the proportions of their subtypes and their co-expression patterns differ between the left and right sides (Kononenko et al., 2017; Watanabe et al., 2021). Activation of lateralized receptors by endogenous neurohormones may be the basis of the decoding process in the bipartite T-NES.

Bilateral deafferentation of the lumbar segments in animals with transected cervical spinal cords did not interfere with HL-PA formed after the right-side UBI suggesting that spinal reflexes are not involved and that the asymmetry persisted due to activation of efferent neurons or changes in neuromuscular system. In contrast, bilateral lumbar rhizotomy abolished HL-PA formed after the left UBI suggesting that its maintenance requires an afferent input and depends on spinal reflexes. Thus the left and right T-NES counterparts may act through different neural mechanisms and nonetheless enable overall mirror-symmetric outcomes that are the right and left hindlimb flexion, respectively (**Figure 7A,B**).

The asymmetric spinal processing of the UBI effects is in agreement with reported spinal cord asymmetries (de Kovel et al., 2017; Deliagina, Orlovsky, Selverston, & Arshavsky, 2000; Hultborn & Malmsten, 1983a, 1983b; Kononenko et al., 2017; Malmsten, 1983; Nathan, Smith, & Deacon, 1990; Ocklenburg et al., 2017; Zhang et al., 2020). Three-quarters of cervical spinal cords are asymmetric with a larger right side (Nathan et al., 1990). Mono- and polysynaptic spinal reflexes showed rightward lateralization (Hultborn & Malmsten, 1983a, 1983b; Malmsten, 1983; Zhang et al., 2020). Lateralized signals from injured brain that target efferent spinal neural circuits may be clinically relevant.

Patients with stroke and cerebral palsy often do not relax their muscles – they are tonically constricted without any voluntary command. This phenomenon is defined as “stretch- and effort unrelated sustained involuntary muscle activity following central motor lesions” and called spastic dystonia (Gracies, 2005; Lorentzen, Pradines, Gracies, & Bo Nielsen, 2018; Marinelli et al., 2017). It is regarded as a form of efferent muscle hyperactivity (Baude, Nielsen, & Gracies, 2018; Gracies, 2005) and may have a central mechanism that does not depend on afferent input in contrast to spasticity based on exacerbated reflex excitability (Sheean & McGuire, 2009). Thus HL-PA may model spastic dystonia that may affect more frequently the left than right hindlimb.

### Lateralized regulatory crosstalk between the hypothalamus and spinal cord

We previously reported that gene expression patterns in the spinal cord are lateralized and affected by UBI through humoral pathway with clear differences between the contra- and ipsilesional sides (Kononenko et al., 2017; Lukoyanov et al., 2021; Zhang et al., 2020). Expression of the neurohormonal and neuroplasticity-related genes was also different between the left and right sides in the hypothalamus, and affected by a unilateral cortical lesion in this area and pituitary gland (**Figure 4**). The left UBI decreased the gene expression levels in the ipsilesional hypothalamus with no significant changes on the right side. In the pituitary gland, the UBI resulted in elevation of expression of *Avpr1b* that encodes the Arg-vasopressin V1B receptor. Arg-vasopressin may induce HL-PA with flexion of the right limb in rats with intact brain and mediate the effects of the left UBI on the hindlimb posture (Lukoyanov et al., 2021). These effects were blocked by SSR-149415, the selective antagonist of the V1B receptor that is mainly expressed in the anterior pituitary (Roper, O’Carroll, Young, & Lolait, 2011). It was hypothesized that Arg-vasopressin acting through the V1B receptor on the pituitary corticotropes stimulates the release of the proopiomelanocortin-derived β-endorphin that produces HL-PA with the right hindlimb flexion (Lukoyanov et al., 2021). Changes in the V1B receptor expression suggests plasticity in the Arg-vasopressin system that signals from the injured brain.

#### The left-right side-specific gene regulatory networks

Analysis of gene-gene co-expression patterns suggests that the neurohormonal, neuropeptide and neuroplasticity-related genes form the left and right gene regulatory networks in both the hypothalamus and spinal cord; that these networks are coordinated across these two regions and their sides; and that the UBI perturbs this coordination. In both regions, pairwise gene-gene correlations internal for the LdN and RdN were generally positive, while those between the networks were mostly negative suggesting their antagonism. The hypothalamus - spinal cord correlation patterns were strikingly different between the LdN and RdN (**Figure 6B,C**). In the proportion of positive correlations, the ipsilateral hypothalamic LdN – spinal cord LdN gene-gene co-expression pattern displayed marked left – right asymmetry (L-Ip > R-Ip) (**Figures 6** and **7C**). Similarly, the pattern of ipsilateral inter-area correlations for the RdNs was also asymmetric while the direction was opposite (L-Ip < R-Ip). At the same time, the patterns were almost mirror-symmetric between the LdNs on the left side and the RdNs on the right side (LdN-L-Ip ≍ RdN-R-Ip), and between the RdNs on the left side and the LdNs on the right side (RdN-L-Ip ≍ LdN-R-Ip) (**Figures 6** and **7C**). These ensembles may be defined as “allo-symmetric”. In contrast to the ipsilateral patterns, the diagonal (contralateral) inter-area correlation did not significantly differ between the LdNs and RdNs, and were not affected by the UBI (**Figure 6— figure supplement 1**). Asymmetry and “allo-symmetry” of the LdN and RdN were characteristics of the control group and were impaired by the unilateral brain lesion, e.g., the direction of the left-right LdN asymmetry was reversed after the left UBI.

#### Functional implications

Formation of the LdN and RdN by neurohormonal, neuropeptide and neuroplasticity-related genes, along with the asymmetry and “allo-symmetry” of their patterns were revealed in animals with transected spinal cords suggesting that they were established and remodeled by the T-NES (**Figure 7C,D**). The functional role of the LdN and RdN may be to amplify the inherently weak lateralized effects of individual neurohormones and strengthen the left and right side-specific regulations. This mechanism could operate within and between the left and right halves of CNS areas, and across CNS regions and their sides along the neuraxis.

The LdN *Avp* and *Pomc* genes give rise to Arg-vasopressin and β-endorphin that induce right hindlimb flexion in animals with intact brain and may mediate the effects of the left-side UBI (Klement’ev et al., 1986; Lukoyanov et al., 2021). In contrast, Met-enkephalin and dynorphin derived from *Penk* and *Pdyn*, constituents of the RdN, produce flexion of the left hindlimb (Bakalkin et al., 1980; Bakalkin & Kobylyansky, 1989; Chazov et al., 1981; Watanabe et al., 2020). Hypothalamic neurohormones oxytocin and TRH, whose genes are constituents of the LdN, and gonadotropin-releasing hormone, transcribed from the RdN *Ghrh* gene, produce lateralized responses and may regulate lateralized brain functions. The oxytocin receptors mediate the effects of this peptide released from the hypothalamus on pup retrieval behavior through activation of the auditory cortex on the left but not right side (Marlin, Mitre, D’Amour J, Chao, & Froemke, 2015). In human brain, the opioid, Arg-vasopressin and oxytocin systems are lateralized and could mediate lateralized responses (Kantonen et al., 2020; Soriano, Daniels, Prinsen, & Alaerts, 2020; Watanabe et al., 2015; Zink et al., 2011). TRH showed a substantial left side predominance in the hypothalamic nuclei and produced an asymmetric behavior (Borson-Chazot et al., 1986; Chepurnov, 1994; Efimova, Chepurnova, & Chepurnov, 1989). Gonadotropin-releasing hormone is asymmetrically expressed in the hypothalamus and involved in circadian regulation of reproductive functions by acting on the right side (Bakalkin et al., 1984; Cruz, Flores, & Dominguez, 2014; de la Iglesia, Meyer, & Schwartz, 2003; Moran, Cruz, & Dominquez, 1994).

An intriguing possibility is if neurohormones and neuropeptides with asymmetric actions are organized into the left and right sided functional networks that control the entire left and right hemispheres, respectively. These networks may differ between the hemispheres and their integral activities are balanced between the left and right sides. This hypothesis was addressed by analysis of the effects of peptide pools prepared from the left and right brain hemispheres in the HL-PA model that was used to analyze the binary, the left *vs*. right sided responses (Bakalkin, Iarygin, Kobylianskiĭ, Samovilova, & Klement’ev, 1981; Bakalkin, Pivovarov, Kobylyansky, Yarygin, & Akparov, 1989; Kryzhanovskii, Lutsenko, Karganov, & Beliaev, 1984; Vartanian, Shatik, Tokarev, & Klement’ev, 1989). The “left” and “right” extracts administered intrathecally caused formation of HL-PA in rats with intact brain and transected spinal cords. Notably the direction of the asymmetry depended on whether an extract was prepared from the left or right hemisphere. The “left” extract induced flexion of left hindlimb, while the right hindlimb was flexed after administration of peptides from the right hemisphere. No asymmetry was formed after administration of the total peptide pool prepared from the whole brain. Thus, peptides with side specific actions were lateralized in the brain, and the integral activity of the “left” and “right” peptide factors was balanced between the hemispheres. Biochemical analysis demonstrated that factors inducing HL-PA were multiple short peptides that remain to be identified at molecular level (Bakalkin, Pivovarov, Kobylyansky, Yarygin, & Akparov, 1989).

### Limitations

The T-NES was identified in anaesthetized animals with transected spinal cords in acute experiments that lasted for 3 – 5 hours after UBI. A role of this phenomenon in the enduring left-right side-specific regulation of biological and pathophysiological processes requires further investigation. Neural pathways that mediate signaling from the injured brain area to the hypothalamic-pituitary system, central and peripheral targets of the neuroendocrine system mediating the effects of brain injury, and afferent, central and efferent mechanisms of HL-PA remain to be investigated.

Categorization of genes into the LdN and RdN was based on direction of their lateralization but not on significance or range of the asymmetry. Nonetheless robust differences between the LdN and RdN in their intra- and inter-area correlation patterns were revealed suggesting that a substantial part of these genes was properly assigned to a respective network. The analyzed set of the neurohormonal and neuroplasticity-related genes allowed to reveal the LdN and RdN patterns, however it was relatively small. Further transcriptome-wide analysis could reveal a complete structure of the left-right side-specific gene regulatory networks.

## Conclusions

Our previous and present findings indicate that, in addition to neural pathways, the contralateral effects of brain lesions are mediated by the novel topographic neuroendocrine mechanism. The discovered T-NES may bypass descending neural tracts, convey information on brain injury and its side to the spinal cord through the bloodstream, and induce contralateral functional deficits (Bakalkin, 2022; Lukoyanov et al., 2021; Watanabe et al., 2021; Watanabe et al., 2020).

The T-NES is bipartite and consists of the left and right counterparts that mediate the effects of the left-side and right-side brain injury, respectively (**Figure 7**). These counterparts enable overall mirror-symmetric functional responses, e.g., flexion of the right and left hindlimbs, respectively, however, the underlying neural and neurohormonal mechanisms are different. The left T-NES-induced HL-PA formation requires an afferent input while the effects of right T-NES may develop through activation of efferent neurocircuits. Antagonists of the δ-, κ- and µ-opioid receptors differently inhibited neurohormonal signaling from the left and right hemispheres suggesting that the endogenous opioid peptides convey the “left” and “right” signals through the humoral pathway or differentially control their processing in the hypothalamus or spinal cord.

Analysis of gene-gene co-expression patterns revealed the left-right side-specific gene regulatory networks and their ipsilateral coordination across the hypothalamus and spinal cord (**Figure 7**).

The ipsilateral interactions differed between the left and right gene networks on each body side and for both networks between the sides. The left UBI perturbed these patterns by affecting the LdN. The findings suggest the side-specific ipsilateral endocrine crosstalk between the hypothalamus and lumbar spinal cord that coordinates molecular processes between these regions and is reorganized in the response to a unilateral brain lesion.

Functional specialization of the left and right hemispheres is an organizing principle of the brain (Concha, Bianco, & Wilson, 2012; Duboc, Dufourcq, Blader, & Roussigne, 2015; Gunturkun, Strockens, & Ocklenburg, 2020; MacNeilage, Rogers, & Vallortigara, 2009). Lateralized processes may be regulated by the side-specific neurohormonal mechanisms that operate either on the left or right side (Allen, Bobnar, & Kolber, 2021; Bakalkin, 2022; Deliagina et al., 2000; Hussain et al., 2012; Kawakami et al., 2003; Kononenko et al., 2017; Kononenko et al., 2018; Marlin et al., 2015; Nation et al., 2018; Phelps, Navratilova, Dickenson, Porreca, & Bannister, 2019; Watanabe et al., 2015; Watanabe et al., 2020; Zink et al., 2011). Our findings suggest that the lateralized neuroendocrine system has a more general role. The fundamental features of the bilaterian body are its symmetric organization and functioning that require a robust control of the balance between the left and right sided processes. We hypothesize that the bipartite T-NES is a part of this left-right side-specific control mechanism. The T-NES may be based on the lateralized neurohormonal networks, and may operate locally (e.g., within the brain and spinal cord areas), or along the neuraxis by signaling from the left and right hemispheres to the ipsilateral or contralateral body sides. Unilateral brain or body lesion could shift this balance to the left or to the right, depending on the injury side, and thereby impair the left-right side-specific neurohormonal control leading to asymmetric functional impairments. From a clinical perspective, it is essential to weight up the contribution of neural and endocrine pathways to protracted neurological deficits secondary to TBI and stroke including hemiparesis and hemiplegia, and to develop pharmacological means to reinstate the impaired neurohormonal balance.

## Supporting information

SOURCE FILE for Figures 1,2

SOURCE FILE for Figure 3

SOURCE FILE for Figures 4,6

SOURCE FILE for Figure 4D

SOURCE FILE for Figures 4 - 6

SOURCE FILE for Figures 4 - 6

SOURCE FILE for Figures 5,6

## Author contribution

H.W., O.N., L.C., G.H.M., N.L. and M.Z., performed injury, behavioral, morphological and molecular analysis. Y.K., V.G. and D.S. performed statistical analyses. H.W., N.L., I.L., M.H., J.S., M.Z. and G.B. planned the experiments, processed and discussed the data, and participated in manuscript preparation. J.S., M.Z. and G.B. conceived and supervised the project. G.B. wrote the manuscript. All authors worked with and commented on the manuscript.

## Acknowledgements

We are grateful to Dr. Michael Ossipov for discussion and manuscript processing, and Ms. Karen Rich for assistance with histochemical analysis.

## MATERIALS AND METHODS

### Animals

Adult male Sprague Dawley rats (Taconic, Denmark) weighing 190-410 g were used in the study. The animals received food and water ad libitum, and were kept in a 12-h day-night cycle (light on from 10:00 p.m. to 10:00 a.m.) at a constant environmental temperature of 21°C (humidity: 65%) and randomly assigned to their respective experimental groups. Approval for animal experiments was obtained from the Malmö/Lund ethical committee on animal experiments (No.: M7-16. Experiments were performed from 9:00 a.m. to 8:00 p.m. After the experiments were completed, the animals were given a lethal dose of pentobarbital.

### Spinal cord transection

The animals were anesthetized with sodium pentobarbital anesthesia (intraperitoneal, I.P.; 60 mg/kg body weight, as an initial dose and then 6 mg/kg every hour). Core temperature of the animals was controlled using a feedback-regulated heating system.

The experimental design included rats with UBI which was preceded by a complete spinal cord transection. Anaesthetized animals were mounted onto the stereotaxic frame and the skin of the back was incised along the midline at the level of the superior thoracic vertebrae. After the back muscles were retracted to the sides, a laminectomy was performed at the C6 and C7 vertebrae. A 3-4-mm spinal cord segment between the two vertebrae was dissected and removed (Lukoyanov et al., 2021). The completeness of the transection was confirmed by (i) inspecting the cord during the operation to ensure that no spared fibers bridged the transection site and that the rostral and caudal stumps of the spinal cord were completely retracted; and (ii) examining the spinal cord in all animals after termination of the experiment.

### Brain surgery

The head of the rats mounted onto the stereotaxic frame was fixed in a position in which the bregma and lambda were located at the same horizontal level. After local injection of lidocaine (Xylocaine, 3.5 mg/ml) with adrenaline (2.2 μg/ml), the scalp was cut open and a piece of the parietal bone located 0.5 – 4.0 mm posterior to the bregma and 1.8 – 3.8 mm lateral to the midline (Paxinos & Watson, 2007) was removed. The part of the cerebral cortex located below the opening that includes the hind-limb representation area of the sensorimotor cortex was aspirated with a glass pipette (tip diameter 0.5 mm) connected to an electrical suction machine (Craft Duo-Vec Suction unit, Rocket Medical Plc, UK). Care was taken to avoid damaging the white matter below the cortex. After the ablation, bleeding was stopped with a piece of Spongostone and the bone opening was covered with a piece of TissuDura (Baxter, Germany). For sham operations, animals underwent the same anesthesia and surgical procedures, but the cortex was not ablated.

After completion of all surgical procedures, the wounds were closed with the 3-0 suture (AgnTho’s, Sweden) and the rat was kept under an infrared radiation lamp to maintain body temperature during monitoring of postural asymmetry and during stretching force analysis.

### Dorsal rhizotomy

A bilateral dorsal rhizotomy was performed in rats with complete transection of the cervical spinal cords 3 h after the UBI. After laminectomy from the T11 to L3 vertebral level, the dura was opened, and the dorsal roots were cut bilaterally from the L1 to S2 spinal levels with a pair of fine scissors as close to their exit as possible from the spinal column so that the spinal cord was not damaged. After the cutting the dorsal rootlets were flipped to make sure that the rhizotomy was complete. This procedure prevents hindlimb afferent input to the spinal cord as demonstrated elsewhere (Goldberger, 1988; Lavrov et al., 2008; Noguchi, Ohta, Kakinoki, Kaizawa, & Matsuda, 2013; Takahashi & Nakajima, 1996). The HL-PA and stretching resistance were analyzed before UBI, 3 h after UBI before rhizotomy, and 0.5 h after rhizotomy.

### Histological analysis of brain injury

Localization and size of cortical lesions were analyzed in rats with left side (n = 5) and right side (n = 5) UBI 3 – 5.5 hours after the injury. After perfusion with 4% paraformaldehyde the brain was removed and postfixed in the same fixative overnight. Then the brain was soaked in phosphate-buffered saline with 30% sucrose for 48 hours, dissected into blocks which were then sliced into 50 µm sections with a freezing microtome. Every fourth section was stained with toluidine (Nissl stain), and all the stained sections across the lesion site were photographed and the rostrocaudal respective mediolateral extension as well as lesion volume were calculated.

### Analysis of hindlimb postural asymmetry (HL-PA) by the hands-on and hands-off methods

The HL-PA value and the side of the flexed limb were assessed as described elsewhere (Lukoyanov et al., 2021; Watanabe et al., 2020; Zhang et al., 2020). Briefly, the measurements were performed under pentobarbital (40 mg/kg, i.p.). The level of anesthesia was characterized by a barely perceptible corneal reflex and a lack of overall muscle tone. The anesthetized rat was placed in the prone position on the 1-mm grid paper.

In the hands-on analysis, the hip and knee joints were straightened by gently pulling the hindlimbs backwards for 1 cm to reach the same level. Then, the hindlimbs were set free and the magnitude of postural asymmetry was measured in millimeters as the length of the projection of the line connecting symmetric hindlimb distal points (digits 2-4) on the longitudinal axis of the rat. The procedure was repeated six times in immediate succession.

In the hands-off method, silk threads were glued to the nails of the middle three toes of each hindlimb, and their other ends were tied to one of two hooks attached to the movable platform that was operated by a micromanipulator constructed in the laboratory (Lukoyanov et al., 2021). To reduce potential friction between the hindlimbs and the surface with changes in their position during stretching and after releasing them, the bench under the rat was covered with plastic sheet and the movable platform was raised up to form a 10° angle between the threads and the bench surface. Positions of the limbs were adjusted to the same, symmetric level, and stretching was performed for the 1.5 cm distance at a rate of 2 cm/sec. The threads then were relaxed, the limbs were released and the resulting HL-PA was photographed. The procedure was repeated six times in succession, and the HL-PA values for a given rat were used in statistical analyses.

The limb displacing shorter projection was considered as flexed. The HL-PA was measured in mm with negative and positive HL-PA values that were assigned to rats with the left and right hindlimb flexion, respectively. This measure, the postural asymmetry size (PAS) shows the HL-PA value and flexion side. The PAS does not show the proportion of the animals with asymmetry in each group, whether all or a small fraction of animals display the asymmetry; and cannot be used for analysis of rat groups with the similar number of left or right flexion. In the latter case the HL-PA value would be about zero. Therefore, the HL-PA was also assessed by the magnitude of postural asymmetry (MPA) that shows absolute flexion size, and the probability of postural asymmetry (P_A_) that shows the proportion of animals exhibiting HL-PA at the imposed threshold (> 1 mm). The MPA and P_A_ do not show flexion side. These three measures are obviously dependent; however, they are not redundant and for this reason, all are required for characterization of the HL-PA data and presentation.

### Analysis of hindlimb resistance to stretch

Stretching force was analyzed under pentobarbital anesthesia within 3-5 hrs after UBI using the micromanipulator-controlled force meter device constructed in the laboratory (Zhang et al., 2020). Two Mark-10 digital force gauges (model M5-05, Mark-10 Corporation, USA) with a force resolution 50 mg were fixed on a movable platform operated by a micromanipulator. Three 3-0 silk threads were glued to the nails of the middle three toes of each hindlimb, and their another ends were hooked to one of two force gauges. The flexed leg of the rat in prone position was manually stretched to the level of the extended leg; this position was taken as 0 mm point. Then both hind limbs were stretched in strictly caudal direction, moving the platform by micromanipulator at the constant 5 mm/sec speed for 10 mm. No or very little trunk movement was observed at stretching for the first 10 mm, and therefore the data recorded for this distance were included in statistical analysis. The force (in grams) from two gauges was simultaneously recoded with 100 Hz frequency during stretching. Five successive ramp-hold-return stretches were performed as technical replicates. Because the entire hindlimb was stretched the measured resistance was characteristic of the passive musculo-articular resistance integrated for hindlimb joints and muscles (Marsala et al., 2005; Nordez, Casari, & Cornu, 2008; Nordez, Casari, Mariot, & Cornu, 2009). The resistance analyzed could have both neurogenic and mechanical components, but their respective contributions were not distinguished in the experimental design. The resistance was measured as the amount of mechanical work W_Left_ and W_Right_ to stretch the left- and right hindlimbs, where W was stretching force integrated over stretching distance interval from 0 to 10 mm.

### Drug treatment design

nor-Binaltorphimine (BNI; 6 mg/kg) and β-Funaltrexamine (FNA; 3 mg/kg) were administered subcutaneously 1 day before the spinal cord and brain surgeries. Naloxone (10 mg/kg), naltrindole (NTI; 5 mg/kg) and saline were administered intraperitoneally 3-4 h after UBI into rats with the MPA > 1.5 mm, and their effects on HL-PA were 1 h later.

Doses and timeline for naloxone (Norris, Perez-Acosta, Ortega, & Papini, 2009), NTI (Nizhnikov, Pautassi, Truxell, & Spear, 2009; Petrillo et al., 2003; Rutten et al., 2018), BNI (Horan et al., 1992; Patkar et al., 2013; Rutten et al., 2018) and FNA (Petrillo et al., 2003) were robustly established in previous studies to block the respective receptors. The dose for naloxone was chosen to block all three subtypes of opioid receptors. BNI and FNA exert long-lasting antagonistic effects that persist for at least 1 month and are receptor selective from 24 h after administration. The antagonists were purchased from Tocris (Minneapolis, MN). All test compounds were dissolved in saline.

### Analysis of gene expression

Gene expression was analyzed in the pituitary gland, and in the left and right halves of each the hypothalamus and lumbar spinal cord that were isolated from the rats with transected spinal cord 3 h after the left UBI (n = 12) or left sham surgery (n = 11). The tissue samples were snap frozen and stored at −80 °C until assay.

#### Quantitative real-time PCR

Total RNA was purified by using RNeasy Lipid Tissue Mini Kit (Qiagen, Valencia, CA, USA). RNA concentrations were measured with Nanodrop (Nanodrop Technologies, Wilmington, DE, USA). RNA (500 ng) was reverse-transcribed to cDNA with the cDNA iScript kit (Bio-Rad Laboratories, CA, USA) according to manufacturer’s protocol. cDNA samples were aliquoted and stored at -20°C. cDNAs were mixed with PrimePCR™ Probe assay and iTaq Universal Probes supermix (Bio-Rad) for qPCR with a CFX384 Touch™ Real-Time PCR Detection System (Bio-Rad Laboratories, CA, USA) according to manufacturer’s instructions. TagMan assay was performed in 384-well format with TagMan probes that are listed in **Figure 4-figure supplements 1-4,7**.

All procedures were conducted strictly in accordance with the established guidelines for the qRCR based analysis of gene expression, consistent with the minimum information for publication of quantitative real-time PCR experiments guidelines (Bustin et al., 2009; Taylor et al., 2019). The raw qPCR data were obtained by the CFX Maestro™ Software for CFX384 Touch™ Real-Time PCR Detection System (Bio-Rad Laboratories, CA, USA). mRNA levels of genes of interest were normalized to the geometric mean of expression levels of two reference genes *Actb* and *Gapdh*. GeNorm software was used to analyze the gene expression stabilities (M value) of the ten candidate reference genes (*Actb, B2m, Gapdh, Gusb, Hprt, Pgk, Ppia, Rplpo13a, Tbp*, and *Tfrc*). The calculation of M value was based on the pairwise variation between two reference genes. If the M value was less than 1.5, it could be considered a suitable reference gene. The smaller the M value, the higher the stability of gene (https://genorm.cmgg.be/ and (Vandesompele et al., 2002)). The expression stability of candidate reference genes was computed for all sets of samples and identified *Actb* and *Gapdh* as the most stably expressed genes. For all three regions analysed, the gene expression stability (M values) did not exceed 0.5. The optimal number of reference genes was determined by calculating pairwise variation (V value) by geNorm program. The V value for *Actb* and *Gapdh*, the top reference genes, was 0.12 that did not exceed the 0.15 threshold demonstrating that analysis of these two genes is sufficient for normalization.

#### Hormonal, neurohormonal and neuroplasticity-related genes

Genes analyzed in the hypothalamus included the hypothalamic neurohormone and neuropeptide genes (**Figure 4—figure supplement 1**; genes of corticotropin releasing hormone *Crh*, growth hormone releasing hormone *Ghrh*, gonadotropin releasing hormone 1*Gnrh1*, neurotensin *Nts*, somatostatin *Sst* and thyrotropin releasing hormone *Trh*); genes of the endogenous opioid system (**Figure 4—figure supplement 2;** genes of δ-opioid receptor *Oprd1*, κ-opioid receptors *Oprk1*, µ-opioid receptors *Oprm1*, prodynorphin *Pdyn*, proenkephalin *Penk* and proopiomelanocortin *Pomc*) and the oxytocin-vasopressin system (**Figure 4—figure supplement 3**; genes of arginine vasopressin *Avp* and oxytocin *Oxt*); and neuroplasticity-related genes (**Figure 4—figure supplement 4**). The *Avpr1a*, *Avpr1b*, and *Avpr2* genes listed in **Figure 4—figure supplement 3** were expressed at low levels and were excluded from further analysis.

The following genes were selected as neuroplasticity-related including *Arc*, activity-regulated cytoskeletal gene implicated in numerous plasticity paradigms; *Bdnf*, brain-derived neurotrophic factor regulating synaptogenesis; *cFos*, a neuronal activity dependent transcription factor; *Dlg4* gene encoding PSD95 involved in AMPA receptor-mediated synaptic plasticity and post NMDA receptor activation events; *Egr1* regulating transcription of growth factors, DNA damage, and ischemia genes; *Gap-43* coding for growth-associated protein Gap-43 that regulates axonal growth and neural network formation; *GluR1* and *Grin2b* coding for the glutamate ionotropic receptor AMPA Type Subunit 1 and NMDA receptor subunit, respectively, both involved in glutamate signaling and synaptic plasticity; *Grin2a* subunit of the glutamate receptors that regulates formation of neural circuits and their plasticity; *Homer-1* giving rise to homer scaffold protein 1, a component of glutamate signaling involved in nociceptive plasticity; *Pcsk6* gene encoding proprotein convertase subtilisin/kexin type 6 involved in post-translational modification; *Nfkbia* (I-Kappa-B-Alpha) that inhibits NF-kappa-B/REL complexes regulating activity-dependent inhibitory and excitatory neuronal function; *Syt4* (Synaptotagmin 4) playing a role in dendrite formation and synaptic growth and plasticity; and *Tgfb1* that gives rise to transforming growth factor β1 regulating inflammation, expression of neuropeptides and glutamate neurotoxicity (Adkins, Boychuk, Remple, & Kleim, 2006; Anderson & Winterson, 1995; Buisson et al., 2003; Dolan, Hastie, Crossan, & Nolan, 2011; Epstein & Finkbeiner, 2018; Grasselli & Strata, 2013; Harris, Zhang, Piccioli, Perrimon, & Littleton, 2016; Hayashi, Ueyama, Nemoto, Tamaki, & Senba, 2000; Joynes, Janjua, & Grau, 2004; Larsson & Broman, 2008; O’Mahony et al., 2006; Santibanez, Quintanilla, & Bernabeu, 2011; Tappe et al., 2006; Vavrek, Girgis, Tetzlaff, Hiebert, & Fouad, 2006; Won, Incontro, Nicoll, & Roche, 2016; You, Morch, & Arendt-Nielsen, 2004) (**Figure 4—figure supplement 4**).

Genes coding for pituitary hormones (**Figure 4—figure supplement 7**; gene of alpha polypeptide of glycoprotein hormones *Cga*, follicle stimulating hormone, subunit beta *Fshb*, glycoprotein hormones, alpha subunit *Gh1*, luteinizing hormone *Lhb*, prolactin *Prl*, and thyroid stimulating hormone, beta subunit *Tshb*), and neuropeptides and their receptors (**Figure 4— figure supplements 2,3)** were analyzed in the pituitary gland.

In the spinal cord, neuroplasticity-related genes (**Figure 4—figure supplement 4)** along with the neuropeptide and their receptor genes (**Figure 4—figure supplements 2,3**) were analyzed besides the *Avp, Avpr1b*, *Avpr2, Oxt* and *Pomc* genes that were expressed at low levels.

### Statistical Analysis

Experimental data were processed and statistically analyzed after completion of the experiments by the statisticians who were not involved in their execution. No intermediate assessment was performed to avoid any bias in data acquisition.

#### Processing of physiological data

##### Bayesian framework

Predictors and outcomes centered and scaled before we fitted Bayesian regression models via full Bayesian framework by calling *Stan 2.21.7* (Stan Development Team, 2022) from *R* 4.1.3 (R Core Team, 2022) using the *brms* 2.18 (Burkner, 2021) interface. To reduce the influence of outliers, models used Student’s *t* response distribution with identity link function unless explicitly stated otherwise. Models had no intercepts with indexing approach to predictors (McElreath, 2019). Default priors were provided by the *brms* according to Stan recommendations (Gelman, 2019). Intercepts, residual SD and group-level SD were determined\from the weakly informative prior student_t(3, 0, 10). The additional parameter ν of Student’s distribution representing the degrees of freedom was obtained from the wide gamma prior gamma(2, 0.1). Group-level effects were determined from the very weak informative prior normal(0, 10). Four MCMC chains of 40000 iterations were simulated for each model, with a warm-up of 20000 runs to ensure that effective sample size for each estimated parameter exceeded 10000 (Kruschke, 2015) producing stable estimates of 95% highest posterior density continuous intervals (HPDCI). MCMC diagnostics were performed according to the Stan manual. P-values, adjusted using the multivariate t distribution with the same covariance structure as the estimates, were produced by frequentist summary in *emmeans* 1.8.4-1 (Lenth, 2023) together with the medians of the posterior distribution and 95% HPDCI. The asymmetry and contrast between groups were defined as significant if the corresponding 95% HPDCI did not include zero and the adjusted P-value was ≤ 0.05. R scripts are available upon request.

##### HL-PA. Postural asymmetry

The magnitude of postural asymmetry (MPA) was inferred via Bayesian framework using Gaussian response distribution. The probability of HL-PA (P_A_) was inferred via Bayesian framework with Bernoulli response distribution and logit link function.

##### Stretching force

The amount of mechanical work W_Left_ and W_Right_ to stretch the left- and right hind limbs, respectively, was computed by integrating the smoothed stretching force measurements over stretching distance from 0 to 10 mm using loess smoothing computed by *loess* function from R package *stats* with parameters span=0.4 and family=”symmetric”. Asymmetry was assessed both as the left / right asymmetry index AI_LR_ = log_2_ (W_Left_ / W_Right_), and for contra- and ipsilesional hindlimbs AI_CI_ = log_2_ (W_Contra_ / W_Ipsi_) and as the difference in work, respectively ΔW_LR_ = (W_Left_ - W_Right_), ΔW_CI_ = (W_Contra_ – W_Ipsi_). The AI and ΔW were inferred via Bayesian framework by fitting linear multilevel models that included *operation type* (left UBI *vs.* right UBI *vs.* sham) as the factor of interest.

#### Molecular Analysis

##### Expression levels

The Lilliefors and Levene’s tests revealed deviations from normality and differences in the variances between the rat groups, respectively, for the expression levels and the asymmetry index (see below for the definition) of several genes in each rat group. The mRNA levels were compared separately for the pituitary gland, and left and right halves of the hypothalamus between UBI or sham surgery groups using Mann-Whitney test followed by

Bonferroni correction for a number of tests (n = 17 and 56, respectively). Fold-change (FC) was computed as a ratio of median expression levels in the UBI to sham groups.

The asymmetry index (AI = median[log_2_L/R)], where L and R were gene expression levels in the left and right halves of hypothalamus or spinal cord, respectively) was computed for each gene in each area (**Figures 4,5**), and compared between UBI and sham surgery groups using Mann– Whitney test followed by a Bonferroni correction for multiple tests (n = 28 and 20 for the hypothalamus and spinal cord, respectively). Because no significant differences between UBI and sham surgery groups were revealed, the groups were combined, and the pooled data were used for analysis of lateralization of gene expression. One-sample version of non-parametric Wilcoxon signed-rank test was applied to compare the AI with zero followed by Bonferroni multiple testing correction (28 genes for hypothalamus). Data for the spinal cord were acquired and analyzed in our previous study (Lukoyanov et al., 2021).

Linear model fitting and analysis were performed in R using *lm, summary.lm* and *confint* commands. Differences were considered to be significant if the P value corrected for multiple comparisons (P_adjusted_) did not exceed 0.05. Heatmaps of Spearman’s rank correlation coefficient were constructed using data (0,1)-standardized for each gene by subtraction of the median value and division by an inter-quartile range.

##### Gene-gene co-expression patterns

In the hypothalamus and spinal cord separately, genes were categorized into two groups that were defined as the left (LdN; AI > 0) and right (RdN; AI < 0) dominant gene regulatory networks (**Figure 4—figure supplement 9; Figure 5—figure supplement 1**). Three categorization variants were used in the following correlation analysis. Genes were assigned into two groups i) by their median AI in the combined UBI and sham surgery group (variant 1); ii) by their median AI in the sham surgery group only (variant 2); and iii) by their mean AI in the combined UBI and sham surgery group (variant 3). Three variants were separately applied for analysis of correlation patterns in each the hypothalamus and SpC, and between these areas (**Source data:** The EXCEL source data files “Table III-S6 23 05 10.xlsx”).

The correlation structure (gene-gene co-expression pattern) for each area, side, between the sides, and across the areas was examines using the Spearman’s rank correlation coefficient calculated for all gene pairs. The pattern of interactions between genes was characterized by the coordination strength (magnitude of correlations or the absolute value of the correlation coefficient averaged across pairwise correlations) and the proportion of positive correlations. In a pattern, edge weights can take values between 0 and 1 for both the magnitude of the correlation coefficient and the proportion of positive correlations.

Robust and unbiased p values were determined in the absence of distributional assumptions by permutation testing. A permutation procedure was employed to characterize the distribution of each statistical test under the null hypothesis of non-replication and non-preservation. Permutation test (Canty A, 2022) with R=10^6^ bootstrap replicates implemented in R/boot package was used to analyze the data. To generate null distribution, we permuted the data across i) *rat identification numbers (IDs)*, ii) *Treatment (UBI and sham surgery)*; iii) *Module* (left and right) within each individual rat; and iv) *CNS area* (hypothalamus and spinal cord) within each module.

The R/boot.pval package was used to compute the P-value (Thulin, 2021). P-values were adjusted using Benjamini-Hochberg family-wise multiple test correction that was applied separately for each set of 6 sets of the tasks (**Source data:** The EXCEL source data files “Table III-S6 23 05 10.xlsx”). Three sets were designed to compare the coordination strength, and other three sets the proportion of positive correlations. Two sets for each the hypothalamus and spinal cord were constructed for analysis of intra-area correlations in the strength and the proportion, and two sets for analysis of the inter-area correlations between the hypothalamus and spinal cord in the strength and the proportion. Each set included comparisons between UBI and sham surgery groups.

##### Executed tasks

(**Figures 4-6**; **Figure 4—figure supplements 10,11; Figure 5—figure supplement 2; Figure 6—figure supplement 1;** (**Source data:** The EXCEL source data files “Table III-S6 23 05 10.xlsx”; raw_groups.xlsx).

###### Task 1

Pairwise gene-gene correlations within each the left (lmLdN-lmLdN vs. lmRdN-lmRdN vs. lmLdN-lmRdN correlations) and right (rmLdN-rmLdN vs. rmRdN-rmRdN vs. rmLdN-rmRdN correlations) modules were compared in each area separately to assess differences in the intra-modular coordination between the gene networks (**Figures 4,5**).

###### Task 2

Pairwise gene-gene intra-modular correlations internal for each the left and right gene networks were compared between the left and right modules (lmLdN-lmLdN vs. rmLdN-rmLdN; lmRdN-lmRdN vs. rmRdN-rmRdN correlations) in each area separately to assess differences in the intra-modular coordination for each gene network between the left and right modules (**Figures 4,5**).

###### Task 3

Pairwise gene-gene inter-modular correlations internal for each the left and right gene networks, were compared between these networks with each other (lmLdN-rmLdN vs. lmRdN-rmRdN correlations) in each area, to assess differences in the inter-modular coordination between the networks (**Figure 4—figure supplement 11; Figure 5—figure supplement 2**).

###### Tasks 4 and 5

The ipsilateral hypothalamus (HPT) and spinal cord (SpC) correlations made either by the left modules (L-Ip: HPT-lmLdN – SpC-lmLdN and HPT-lmRdN – SpC-lmRdN) or right modules (R-Ip: HPT-rmLdN – SpC-rmLdN and HPT-rmRdN – SpC-rmRdN) were compared (a) between the gene networks (LdN vs. RdN) (Task 4), and (b) for each network, between the left and right modules (Task 5), In other words, this analysis assessed differences between (a) the LdN and RdN in each the left and right ipsilateral correlation pattern (Task 4), and (b) between the left and right ipsilateral correlation pattern for each the LdN and RdN (Task 5) (**Figure 6**).

###### Tasks 6 and 7

Each the contralateral left HPT – right SpC correlations (Contralateral type 1 pattern, or Ct1: HPT-lmLdN – SpC-rmLdN and HPT-lmRdN – SpC-rmRdN), and the contralateral right HPT – left SpC correlations (Contralateral type 2 pattern, or Ct2: HPT-rmLdN – SpC-lmLdN and HPT-rmRdN – SpC-lmRdN) were compared between the LdN and RdN (Task 6), and between the Ct1 and Ct2 for each LdN and RdN (Task 7) (**Figure 6—figure supplement 1**).

In each of seven tasks, all correlation patterns were compared between UBI and sham surgery groups.

### Data and code availability

Data supporting the findings of this study are available within the article, its Supporting Information and on https://github.com/YaromirKo/biostatistics-nms.

## Supplementary figures

**Figure 1—figure supplement 1.**
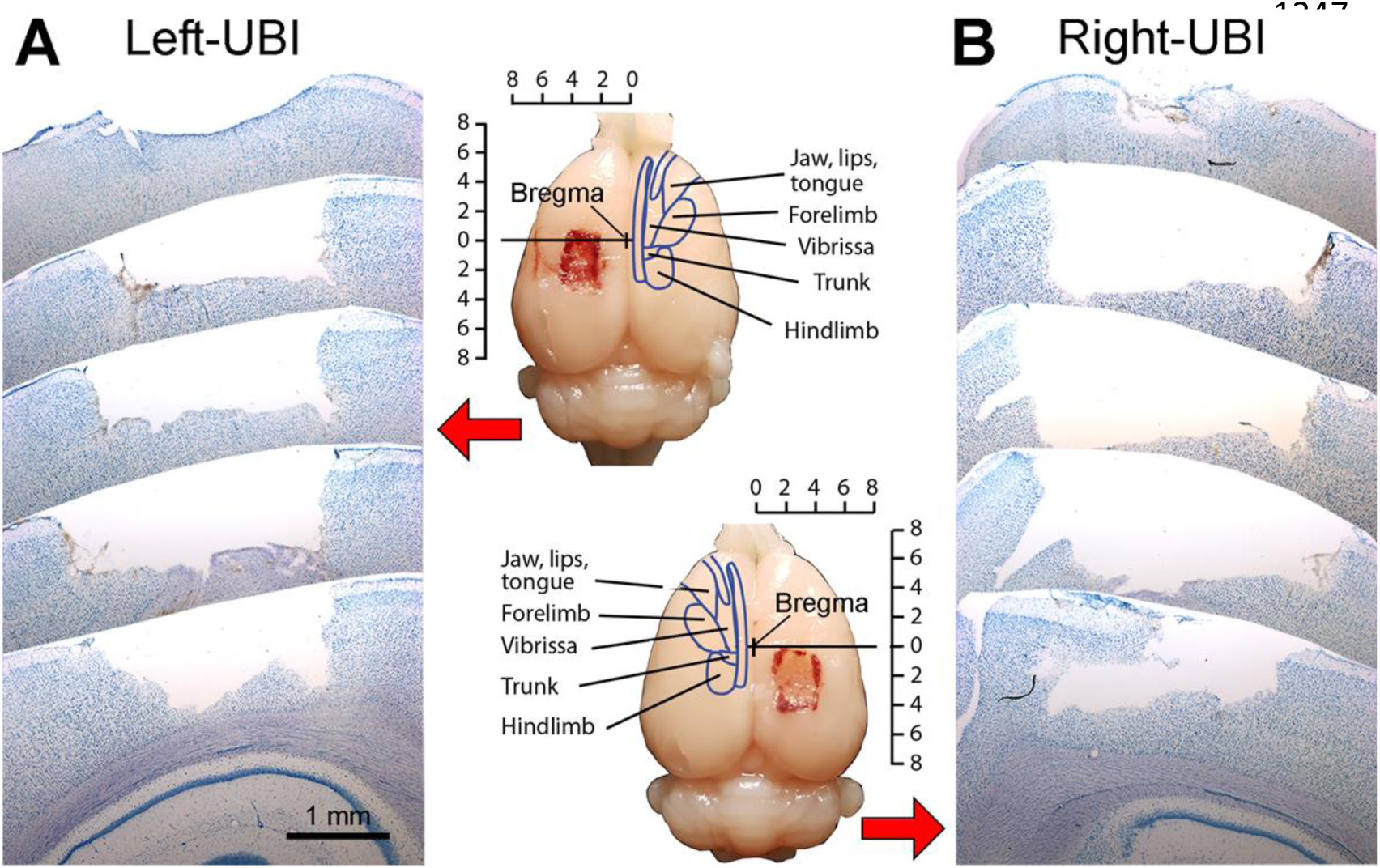
Lesion area in the hindlimb sensorimotor cortex from rats with the left and right unilateral brain injury (UBI) rat. (**A,B**) Five consequent toluidine blue stained cortical sections at an equal distance (1000 μm) across the lesion site in the left and right hemispheres 3 hours after the UBI. Macrographs in the middle show the same brains before sectioning. Delineation on cortex represents somatotopically organized primary motor cortex (modified from (Hall & Lindholm, 1974)). The coordinates are in mm. The scale bar in (**A**) is also valid for (**B**).

**Figure 1—figure supplement 2.**
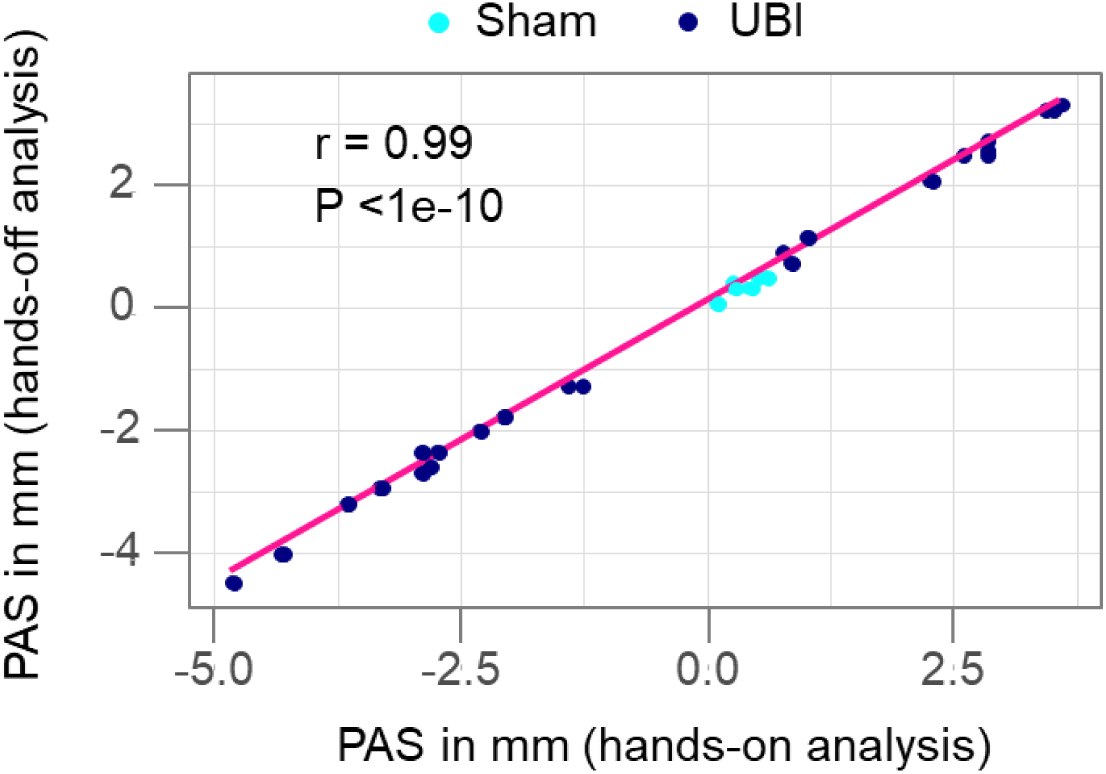
Pearson correlation between the postural asymmetry size (PAS) analyzed by the hands-off and hands-on assay. Data were combined for L-UBI, R-UBI and sham surgery groups of rats with transected spinal cord that were analyzed 3 hours after brain surgery. **Source data:** The EXCEL source data file “masterfile-210807.xlsx ”.

**Figure 1—figure supplement 3.**
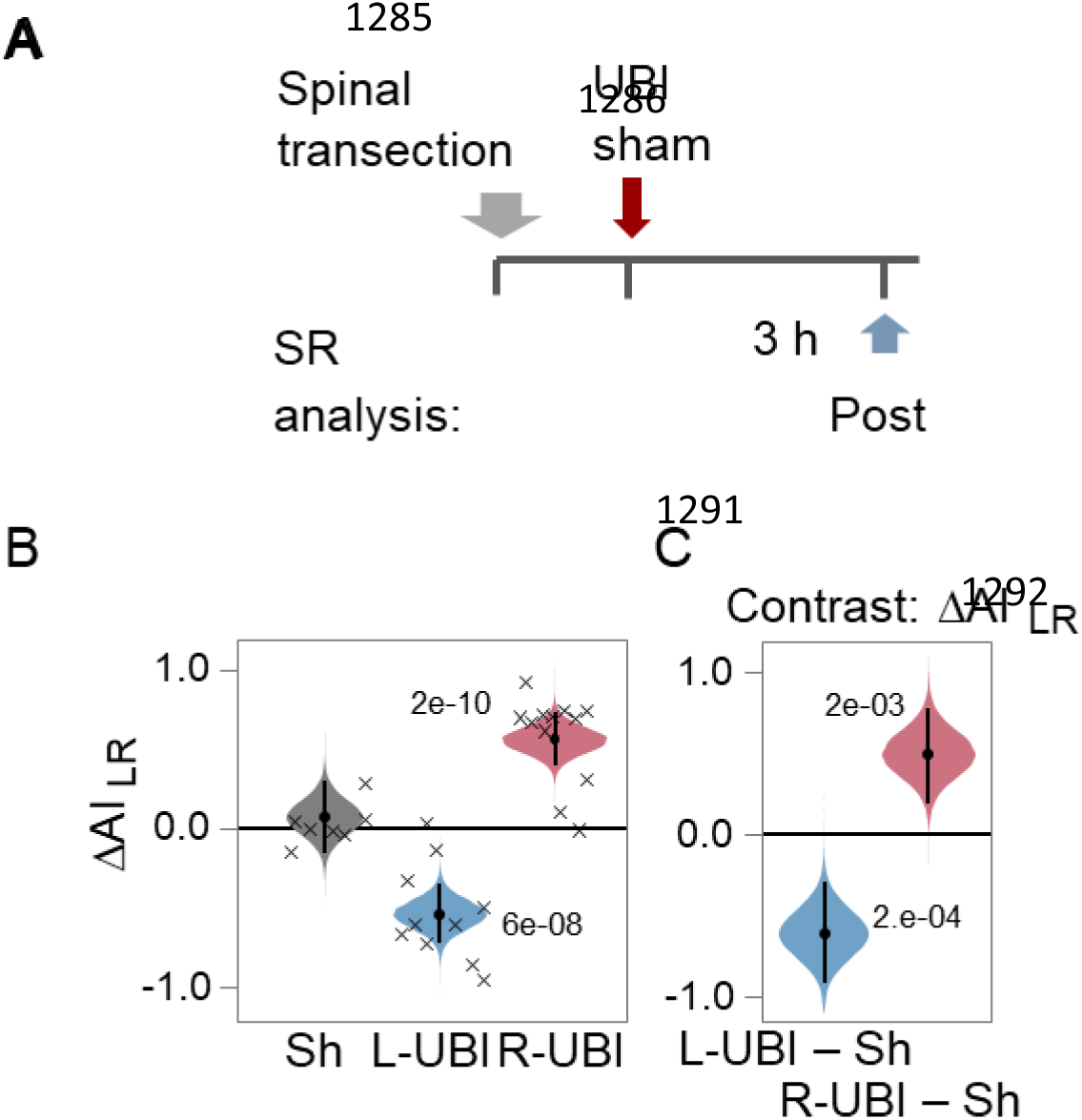
Asymmetry in the hindlimb stretching resistance in rats with completely transected spinal cord: effects of UBI. (**A**) Experimental design. The spinal cord was transected that was followed by L-UBI, R-UBI or sham surgery (Sh). Stretching force was analyzed (**B**,**C**) three hours after UBI or sham surgery (Post; L-UBI, n = 10; R-UBI, n = 12; and sham surgery, n = 7). The UBI effects were analyzed as changes in the asymmetry index for left and right hindlimbs AI_LR_ = log_2_ (W_Left_ / W_Right_). (**C**) Differences (contrast) between the UBI and sham surgery groups. The AI_LR_ and contrast are plotted as median (black circles), 95% HPDC intervals (black lines), and posterior density (colored distribution) from Bayesian regression. Crosses denote the AI_LR_ values for individual rats. Significant effects on the ΔW and the differences between the groups: 95% HPDC intervals did not include zero, and adjusted P-values were ≤ 0.05. Adjusted P is shown for differences identified by Bayesian regression. **Source data:** The EXCEL source data file “masterfile-210807.xlsx” and source data folder “/HL-PA/data/SF/”.

**Table 1.** The number of rats analyzed in experiments with opioid antagonists. Rats with completely transected cervical spinal cord and the left (L-UBI) or right (R-UBI) brain injury were treated with BNI, FNA, NTI, naloxone or saline. Control groups Ctrl1 and Ctrl2 consisted of the rats with transected spinal cords and UBI and were not treated with the antagonists. The Ctrl2 group consisted of rats that were not treated with saline and drugs and were analyzed 3-4 h after UBI, and rats that were treated with saline 3 h after UBI and analyzed 1 h later at Post2 time point (see **Figure 3A**). No significant differences between saline treated and untreated groups in both the MPA and ΔW_CI_ were revealed, and they were combined into the Ctrl2 group. Both the Ctrl1 group that was a control for BNI and FNA, and Ctrl2 group that was a control for NTI and naloxone, were composed of rats with the L-UBI and R-UBI. No statistically significant differences in both the MPA and ΔW_CI_ between the left and right UBI subgroups of each control group were revealed, and the two subgroups were combined into Ctrl1 and Ctrl2 groups, respectively, for statistical analysis.

| Rat group / Treatment | L-UBI rats | R-UBI rats |
| --- | --- | --- |
| Ctrl1 | 17 | 13 |
| BNI | 7 | 9 |
| FNA | 8 | 8 |
| Ctrl2 untreated with drugs or saline | 12 | 12 |
| Ctrl2 treated with saline | 4 | 3 |
| NTI | 8 | 8 |
| Naloxone | 7 | 7 |

**Figure 2—figure supplement 1.**
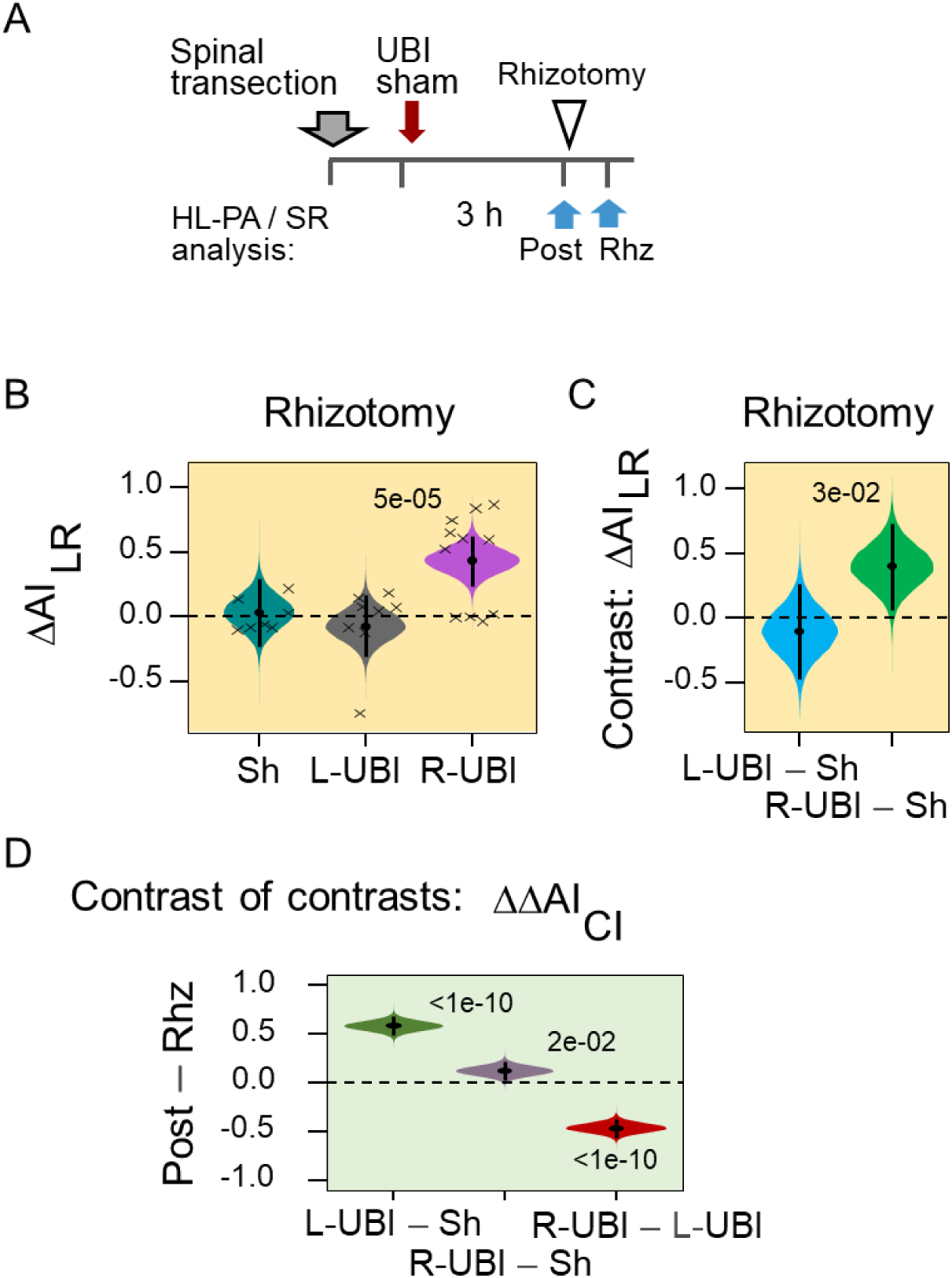
Asymmetry in the UBI-induced hindlimb stretching resistance in rats with completely transected spinal cord: effects of bilateral deafferentation of the lumbar spinal cord. (**A**) Experimental design. The spinal cord was transected that was followed by L-UBI, R-UBI or sham surgery (Sh). Stretching force was analyzed three hours after UBI or sham surgery (Post; L-UBI, n = 10; R-UBI, n = 12; and sham surgery, n = 7); and after bilateral rhizotomy (Rhz) in the subset of rats (L-UBI, n = 8; R-UBI, n = 11; and sham surgery, n = 7). The UBI effects were analyzed as changes in the asymmetry index for left and right hindlimbs AI_LR_ = log_2_ (W_Left_ / W_Right_) (**B,C**), and for contra- and ipsilesional hindlimbs AI_CI_ = log_2_ (W_Contra_ / W_Ipsi_) (**D**). (**B**,**C**) The AI_LR_ and differences (contrast) between the UBI and sham surgery groups after rhizotomy. (**D**) The effects of rhizotomy on differences in AI_CI_ between L-UBI, R-UBI and sham surgery were analyzed as contrast of contrasts i) between L-UBI and sham surgery: [(L-UBI _Post_ – Sh _Post_) – [(L-UBI _Rhz_ – Sh _Rhz_)]; ii) between R-UBI and sham surgery: [(R-UBI _Post_ – Sh _Post_) – [(R-UBI _Rhz_ – Sh _Rhz_)]; and iii) between R-UBI and L-UBI sham surgery: [(R-UBI _Post_ – L-UBI _Post_) – [(R-UBI _Rhz_ – L-UBI _Rhz_)]. Crosses denote the AI_LR_ values for individual rats. The AI_LR_, AI_CL_, contrast and contrast of contrasts are plotted as median (black circles), 95% HPDC intervals (black lines), and posterior density (colored distribution) from Bayesian regression. Significant effects on the ΔW and the differences between the groups: 95% HPDC intervals did not include zero, and adjusted P-values were ≤ 0.05. Adjusted P is shown for differences identified by Bayesian regression. **Source data:** The EXCEL source data file “masterfile-210807.xlsx” and source data folder “/HL-PA/data/SF/”.

**Figure 3—figure supplement 1**

**Figure 4—figure supplement 1**

**Table 2.** Genes coding for hypothalamic releasing and inhibitory hormones and neuropeptide neurotensin, and PCR probes for their analysis (Bio-Rad Laboratories, CA, USA).

| Gene name | Gene symbol | Assay ID | Channel |
| --- | --- | --- | --- |
| *Corticotropin releasing hormone* | *Crh* | qRnoCEP0026204 | FAM |
| *Growth hormone releasing hormone* | *Ghrh* | qRnoCIP0029554 | HEX |
| *Gonadotropin releasing hormone 1* | *Gnrh1* | qRnoCIP0025960 | FAM |
| *Neurotensin* | *Nts* | qRnoCIP0026101 | FAM |
| *Somatostatin* | *Sst* | qRnoCEP0031118 | FAM |
| *Thyrotropin Releasing Hormone* | *Trh* | qRnoCEP0026268 | HEX |

**Figure 4—figure supplement 2**

**Table 3.**
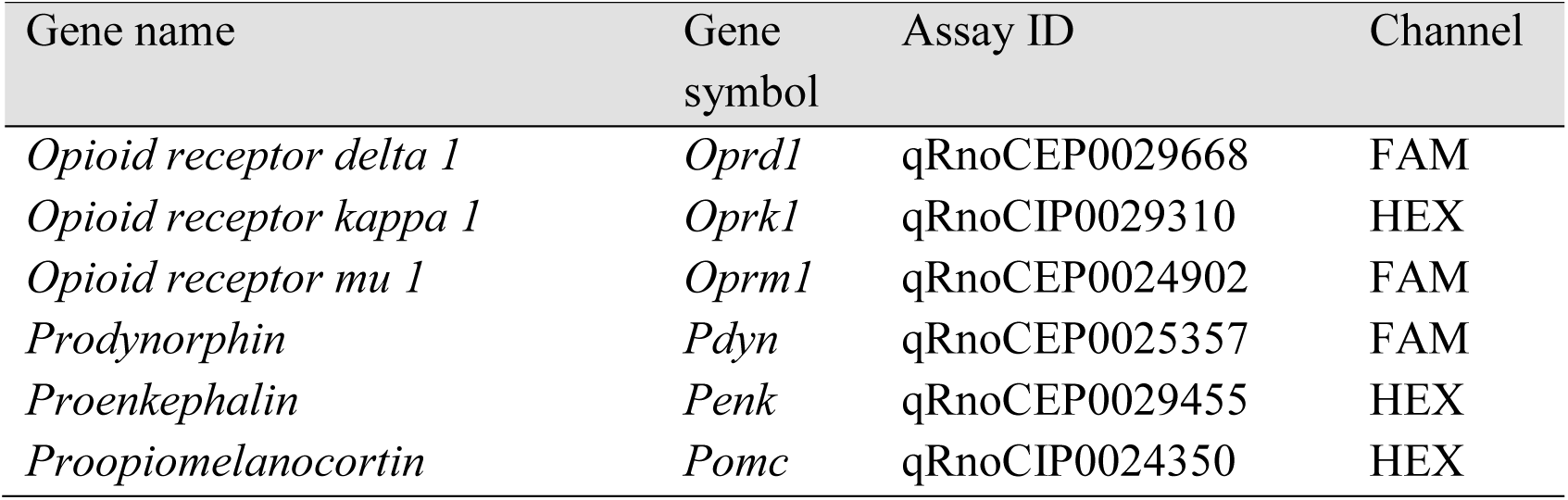
Genes of the endogenous opioid system, and PCR probes for their analysis (Bio-Rad Laboratories, CA, USA).

**Figure 4—figure supplement 3**

**Table 4.** Genes of the oxytocin-vasopressin systems, and PCR probes for their analysis (Bio-Rad Laboratories, CA, USA).

| Gene name | Gene symbol | Structure/ structures analyzed* | Assay ID | Channel |
| --- | --- | --- | --- | --- |
| *Arginine vasopressin* | *Avp* | HPT, PG | qRnoCEP0023611 | FAM |
| *Arginine vasopressin receptor 1A* | *Avpr1a* | PG, SpC | qRnoCEP0023750 | FAM |
| *Arginine vasopressin receptor 1B* | *Avpr1b* | PG | qRnoCEP0037165 | FAM |
| *Arginine vasopressin receptor 2* | *Avpr2* | PG | qRnoCEP0030097 | FAM |
| *Oxytocin* | *Oxt* | HPT, PG | qRnoCEP0031434 | HEX |

\*Structures analyzed: HPT – hypothalamus, PG – pituitary gland, SpC – spinal cord.

**Figure 4—figure supplement 4**

**Table 5.** Neuroplasticity-related genes, and their PCR probes (Bio-Rad Laboratories, CA, USA).

| Gene name | Gene symbol | Assay ID | Channel |
| --- | --- | --- | --- |
| *Activity-regulated cytoskeleton-associated protein* | *Arc* | qRnoCEP0027389 | HEX |
| *Brain-derived neurotrophic factor* | *Bdnf* | qRnoCEP0026843 | HEX |
| *Fos proto-oncogene* | *cFos* | qRnoCEP0024078 | HEX |
| *Discs large MAGUK scaffold protein 4* | *Dlg4* | qRnoCIP0026242 | FAM |
| *Early growth response 1* | *Egr1* | qRnoCEP0022872 | FAM |
| *Growth associated protein 43* | *Gap43* | qRnoCIP0027599 | FAM |
| *Glutamate ionotropic receptor AMPA type subunit 1* | *GluR1* | qRnoCIP0030725 | FAM |
| *Glutamate ionotropic receptor NMDA type subunit 2a* | *Grin2a* | qRnoCIP0025244 | HEX |
| *Glutamate ionotropic receptor NMDA type subunit 2b* | *Grin2b* | qRnoCIP0023973 | HEX |
| *Homer scaffold protein 1* | *Homer-1* | qRnoCEP0023985 | FAM |
| *Proprotein convertase subtilisin/kexin type 6* | *Pcsk6* | qRnoCIP0045340 | FAM |
| *NFKB inhibitor alpha* | *Nfkbia* | qRnoCEP0026759 | HEX |
| *Synaptotagmin 4* | *Syt4* | qRnoCIP0029728 | FAM |
| *Transforming growth factor beta 1* | *Tgfb1* | qRnoCIP0031022 | HEX |

**Figure 4—figure supplement 5.**
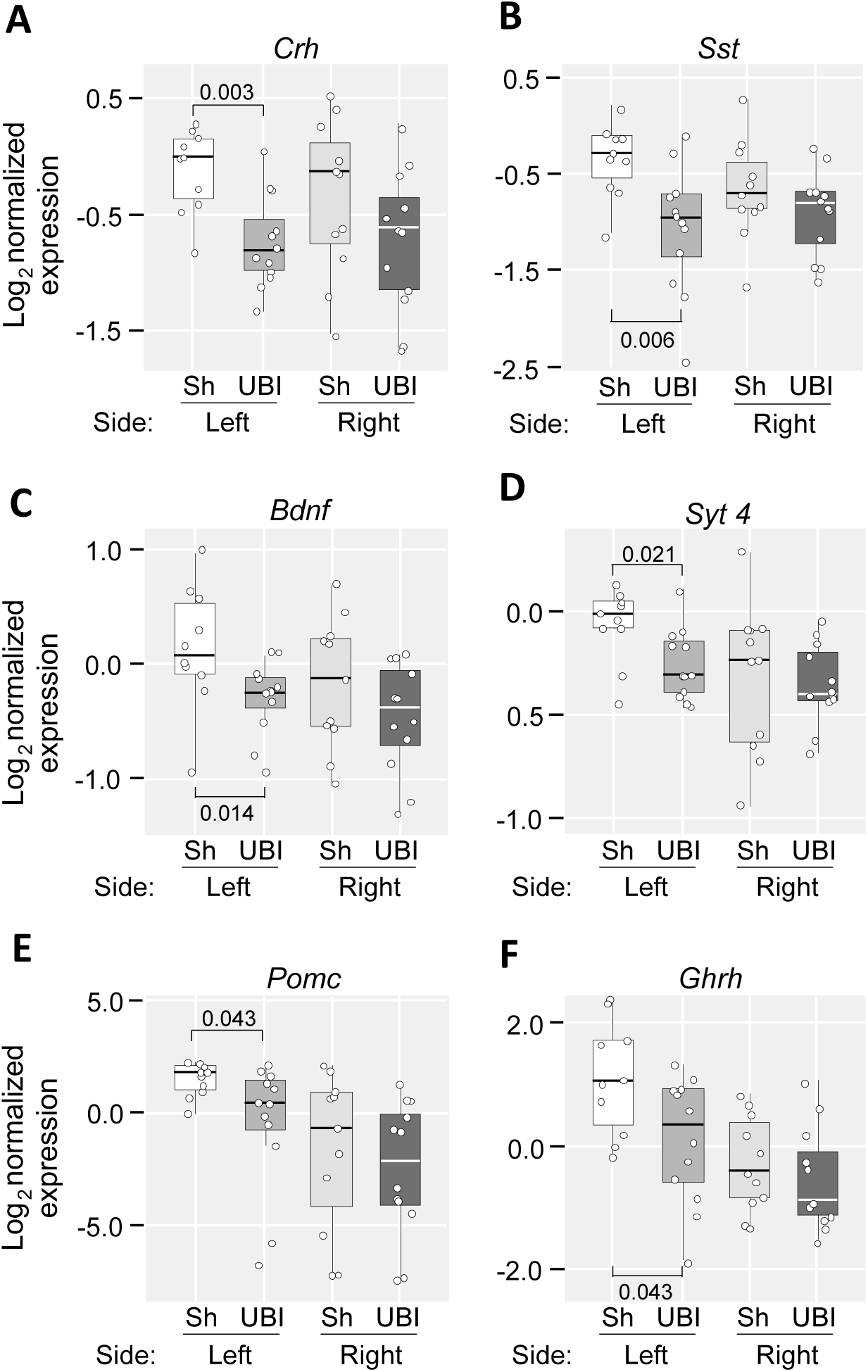
The UBI effects on gene expression in the left and right hypothalamus. Samples were dissected 3 h after left sham surgery (n = 11) or left UBI (n = 12). The expression levels presented in the log_2_ scale as boxplots with median and hinges representing the first and third quartiles, and whiskers extending from the hinge to the highest/lowest value that lies within the 1.5 interquartile range of the hinge. Unadjusted P-values computed using Mann–Whitney test are shown. Fold changes in the left hypothalamus: 1.76x for *Crh* gene, 1.59x for *Sst* gene, 1.25x for *Bdnf* gene, 1.22x for *Syt4* gene, 2.49x for *Pomc* gene, and 1.63x for *Ghrh* gene. **Source data:** The EXCEL source data file “Hypoth_SO_UBI.xlsx”.

**Figure 4—figure supplement 6.**
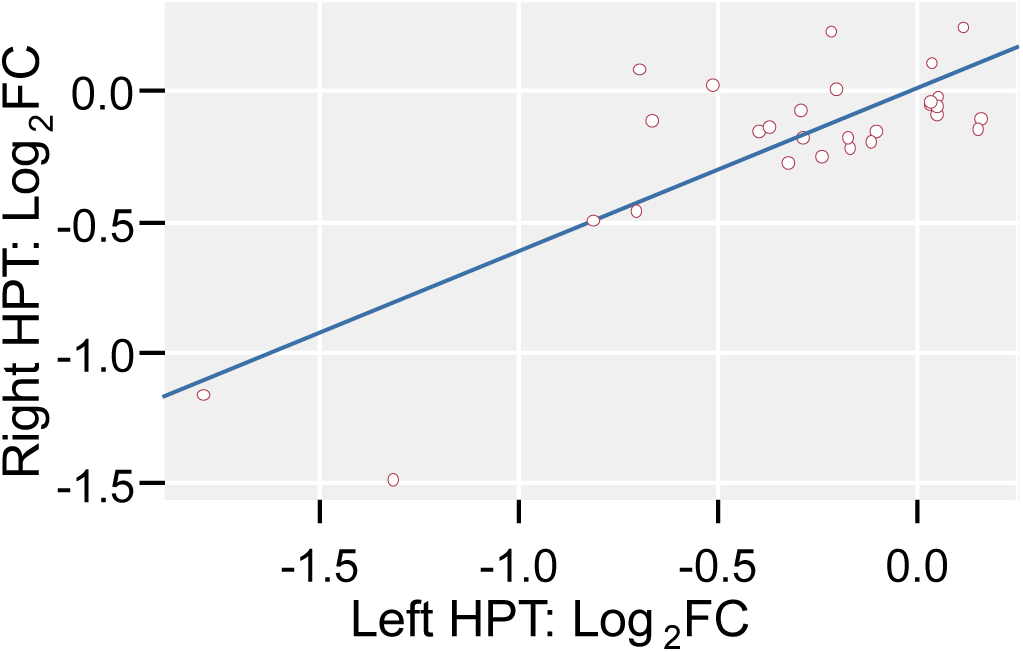
Correlation of the UBI-induced fold changes in the levels of gene expression between the left and right hypothalamus (HPT). Log-scaled fold changes in expression levels of individual genes (log_2_FC, where FC is ratio of median expression levels of UBI and sham groups) are shown. Pearson correlation coefficient: 0.79, P = 5.4×10^-7^; Spearman’s rank correlation coefficient: 0.48, P = 0.010. In a linear model (logFC_right_ ≍ *a* logFC_left_ + *b*), *a* = 0.64 (95% confidence interval [0.45, 0.83]) and *b* = –0.02 (95% confidence interval [-0.12, 0.08]). **Source data:** The EXCEL source data file “Hypoth_SO_UBI.xlsx”.

**Figure 4—figure supplement 7**

**Table 6.**
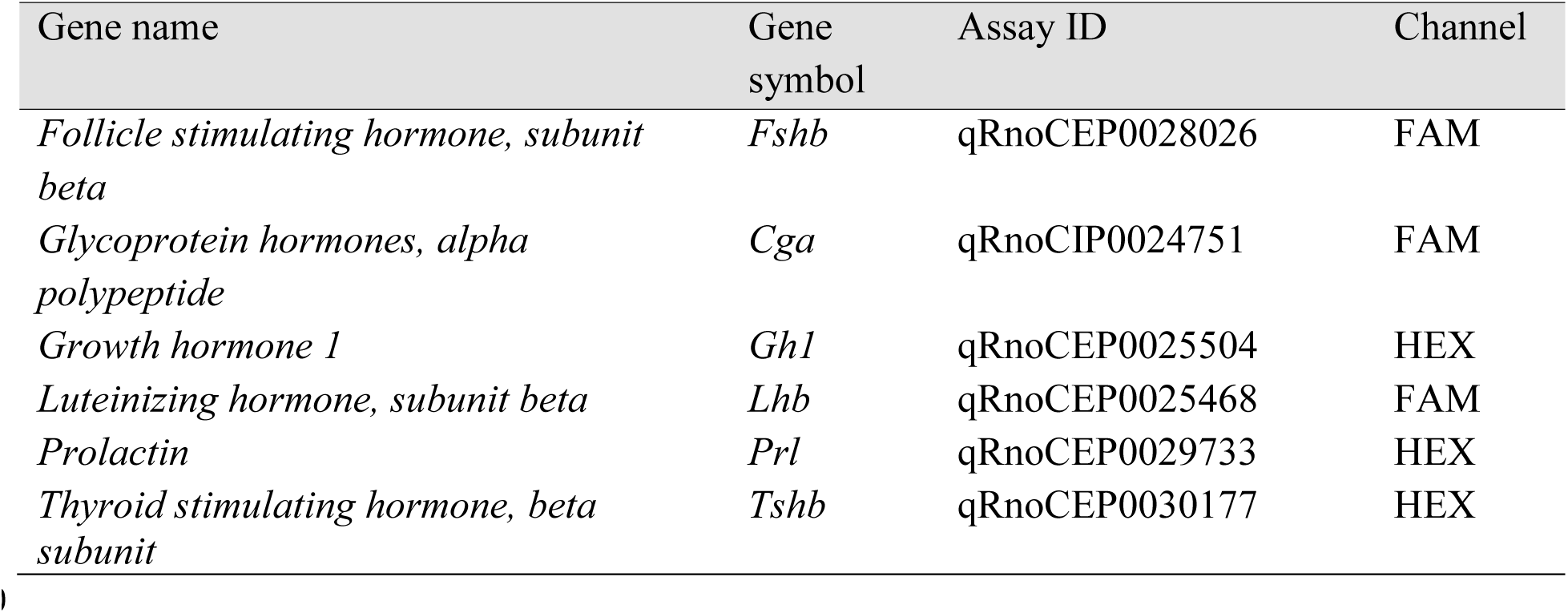
Genes coding for pituitary hormones and PCR probes for their analysis (Bio-Rad Laboratories, CA, USA).

**Figure 4—figure supplement 8.**
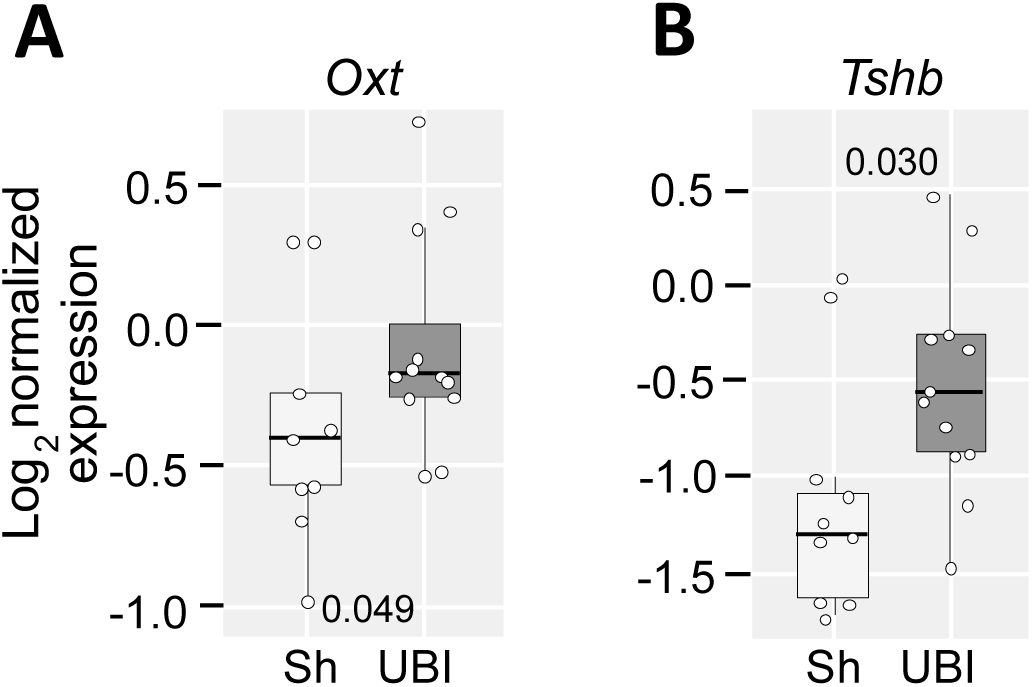
The UBI effects on the levels of gene expression in the pituitary gland. mRNAs were analyzed in the pituitary isolated 3 h after left sham surgery (n = 11) or left UBI (n = 12). The expression levels presented in the log_2_ scale as boxplots with median and hinges representing the first and third quartiles, and whiskers extending from the hinge to the highest/lowest value that lies within the 1.5 interquartile range of the hinge. Unadjusted *P* computed using Mann–Whitney test is shown. FC = 1.17x for *Oxt* gene and FC = 1.63x for *Tshb* gene. **Source data:** The EXCEL source data file “RD Hypophis_ Master file.xlsx”.

**Figure 4—figure supplement 9.**
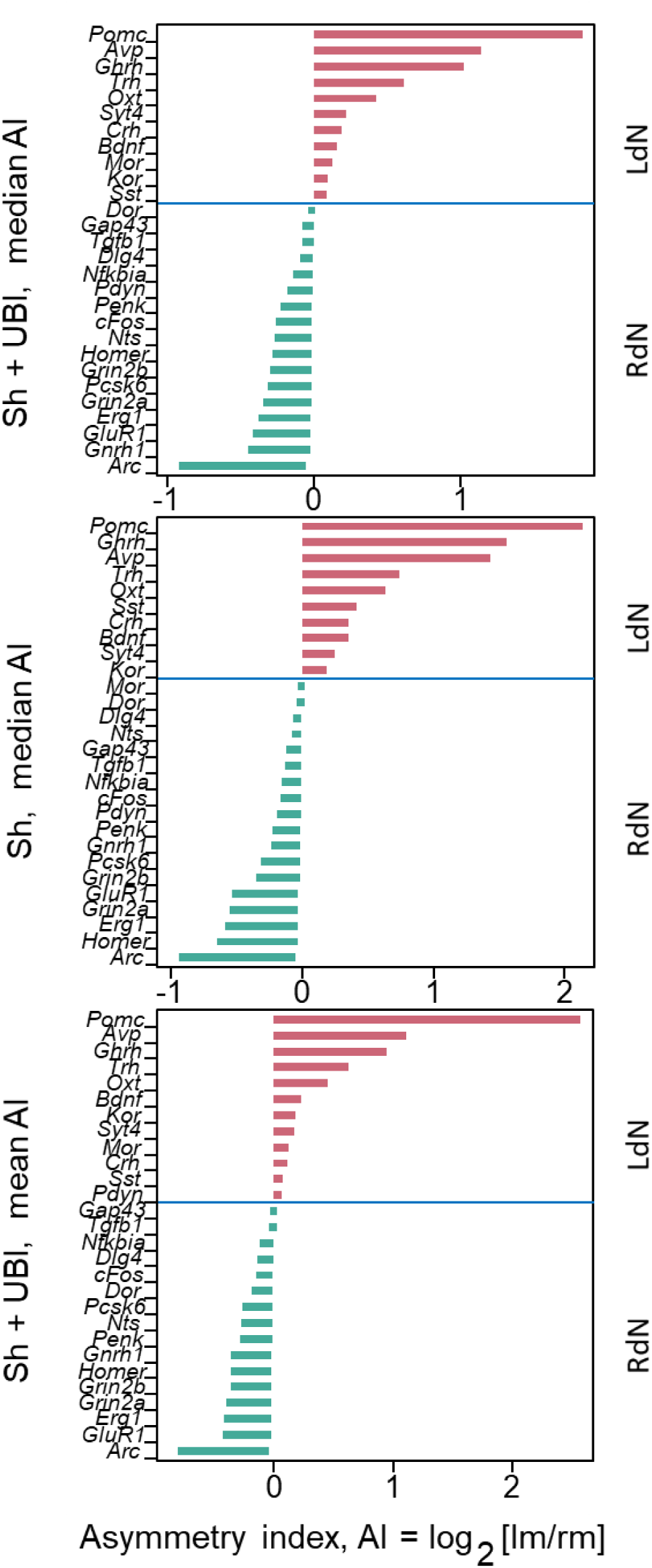
Categorization of genes in the hypothalamus into the left (LdN) and right (RdN) networks using the asymmetry index AI = median[log_2_(lm/rm)], where lm and rm were gene expression modules for left and right halves of the CNS area, respectively. The LdN and RdN are constituted by genes with AI > 0 and AI < 0, respectively. Three categorization variants were designed and used in the following analysis of correlations. Genes were assigned into two groups based on their i) median AI in the combined Sham and UBI group (variant 1); ii) median AI in the sham operation group only (variant 2); and iii) mean AI in the combined sham operation and UBI group (variant 3). Three variants were used in analysis of each the hypothalamus and spinal cord. In the inter hypothalamus –spinal cord analysis, each categorization variant was applied separately for both areas that gave three patterns for the comparisons. **Source data:** The EXCEL source data file “Hypoth_SO_UBI.xlsx; Table III-S6 23 05 10.xlsx; raw_groups.xlsx”.

**Figure 4—figure supplement 10.**
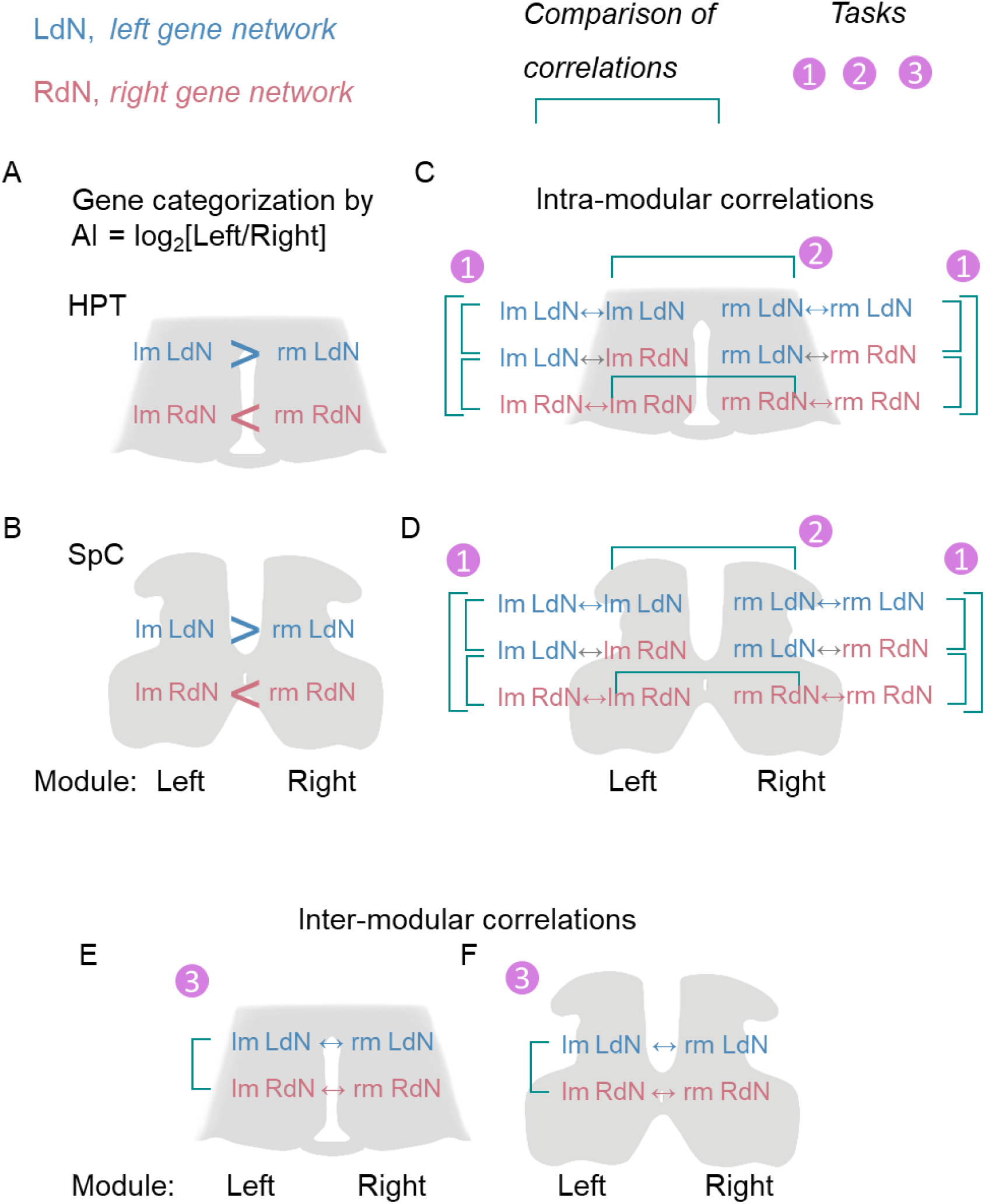
Design of analysis of gene-gene co-expression patterns in both the hypothalamus and spinal cord, and between them (**A,B**) Categorization of genes onto the left (LdN) and right (RdN) networks according to their AI. In each area separately, genes with AI > 0 and AI < 0 were classified into the LdN and RdN, respectively. (**C, D**) Analysis of differences in the intra-modular correlations (LdN-LdN, RdN-RdN, and LdN-RdN) (Task 1) and between the left and right modules in these correlations (Task 2). (**E,F**) Analysis of differences in the inter-modular coordination between the networks (Task 3). Correlations between two variables are shown by double head arrows.

**Figure 4—figure supplement 11.**
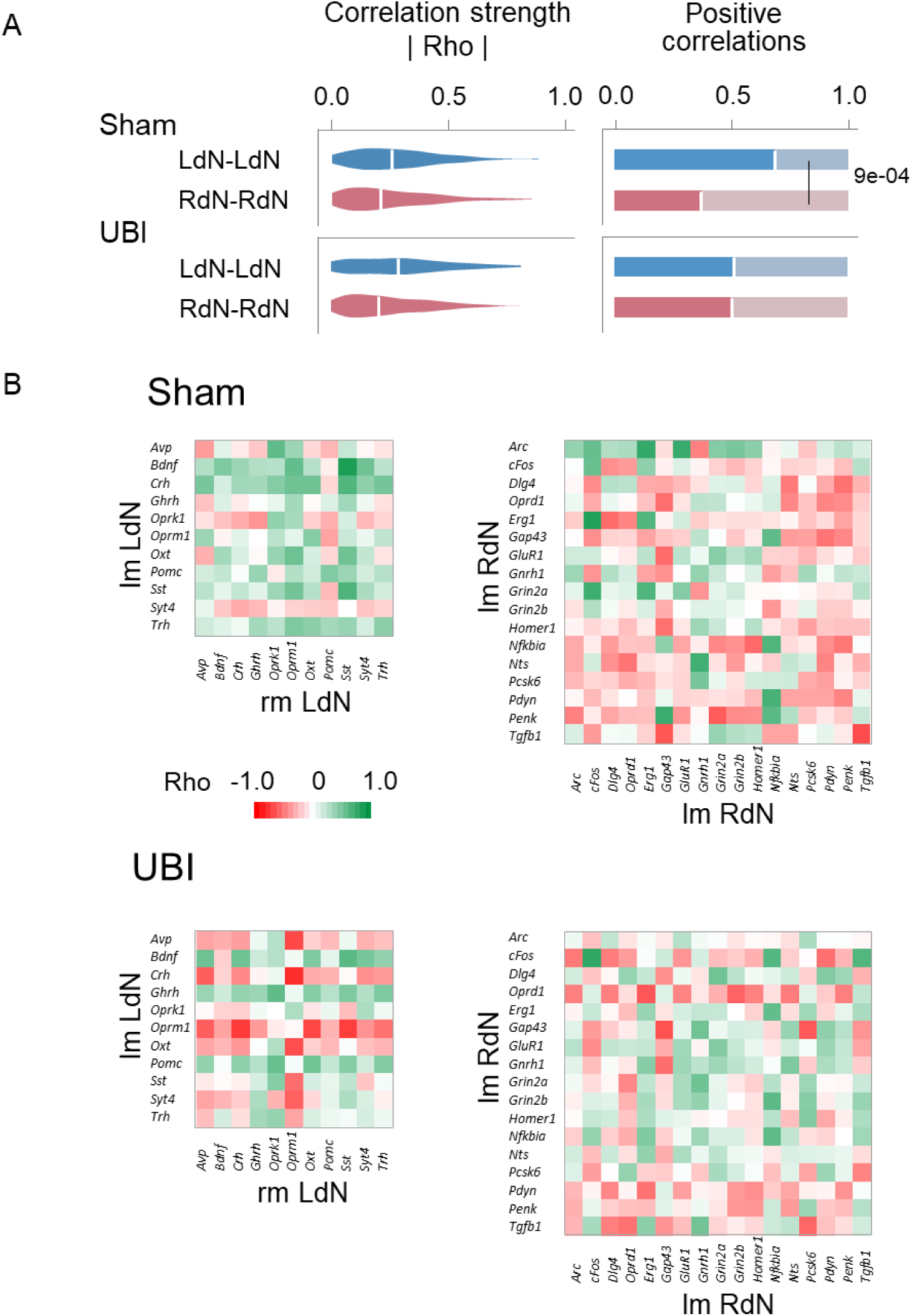
Gene-gene co-expression patterns in the hypothalamus of the sham surgery and UBI rats. (**A**). The coordination strength and the proportion of positive correlations for the inter-modular correlations of the LdN (lmLdN-rmLdN) and RdN (lmRdN-rmRdN). (**B**). Heatmaps for Spearman’s rank coefficients for pairwise gene-gene the inter modular correlations of the LdN (lmLdN-rmLdN) and RdN (lmRdN-rmRdN). Comparison of the patterns between the LdN and RdN, and each of them between sham surgery and UBI groups. P values were determined by permutation testing with Benjamini-Hochberg family-wise multiple test correction, and are shown for the median AI of the combined sham surgery and UBI group. P values shown for the contrasts that are significant (P ≤ 0.05) after the correction for all three categorization variants, or for two of them while in the third variant P was < 0.05 and < 0.10 before and after the correction, respectively. **Source data:** The EXCEL source data file “Hypoth_SO_UBI.xlsx; Table III-S6 23 05 10.xlsx; raw_groups.xlsx”.

**Figure 5—figure supplement 1.**
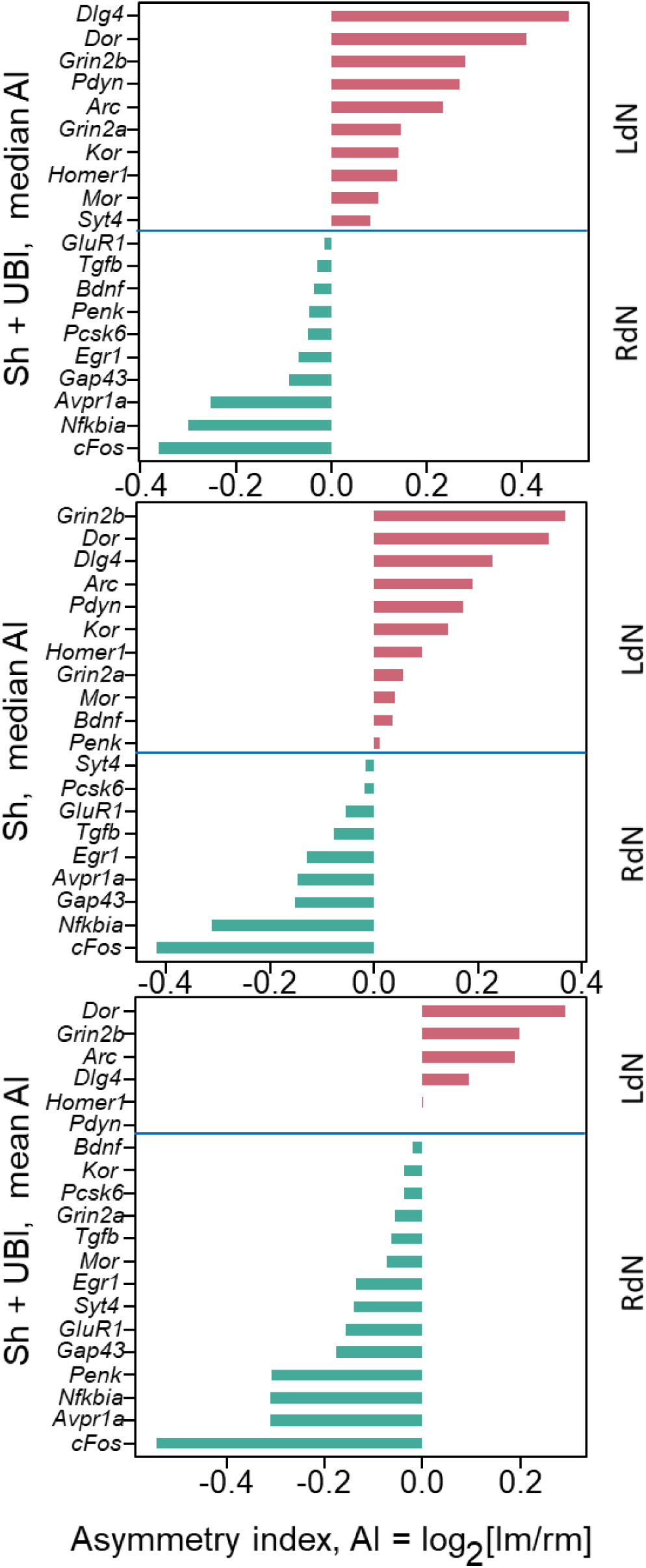
Categorization of genes in the spinal cord into the left (LdN) and right (RdN) networks using the AI. The LdN and RdN are constituted by genes with AI > 0 and AI < 0, respectively. For details, see legend **Figure 4—figure supplement 9**. **Source data:** The EXCEL source data file “SpinalC_SO_UBI_Ctrl_RD_DD.xlsx; Table III-S6 23 05 10.xlsx; raw_groups.xlsx”.

**Figure 5—figure supplement 2.**
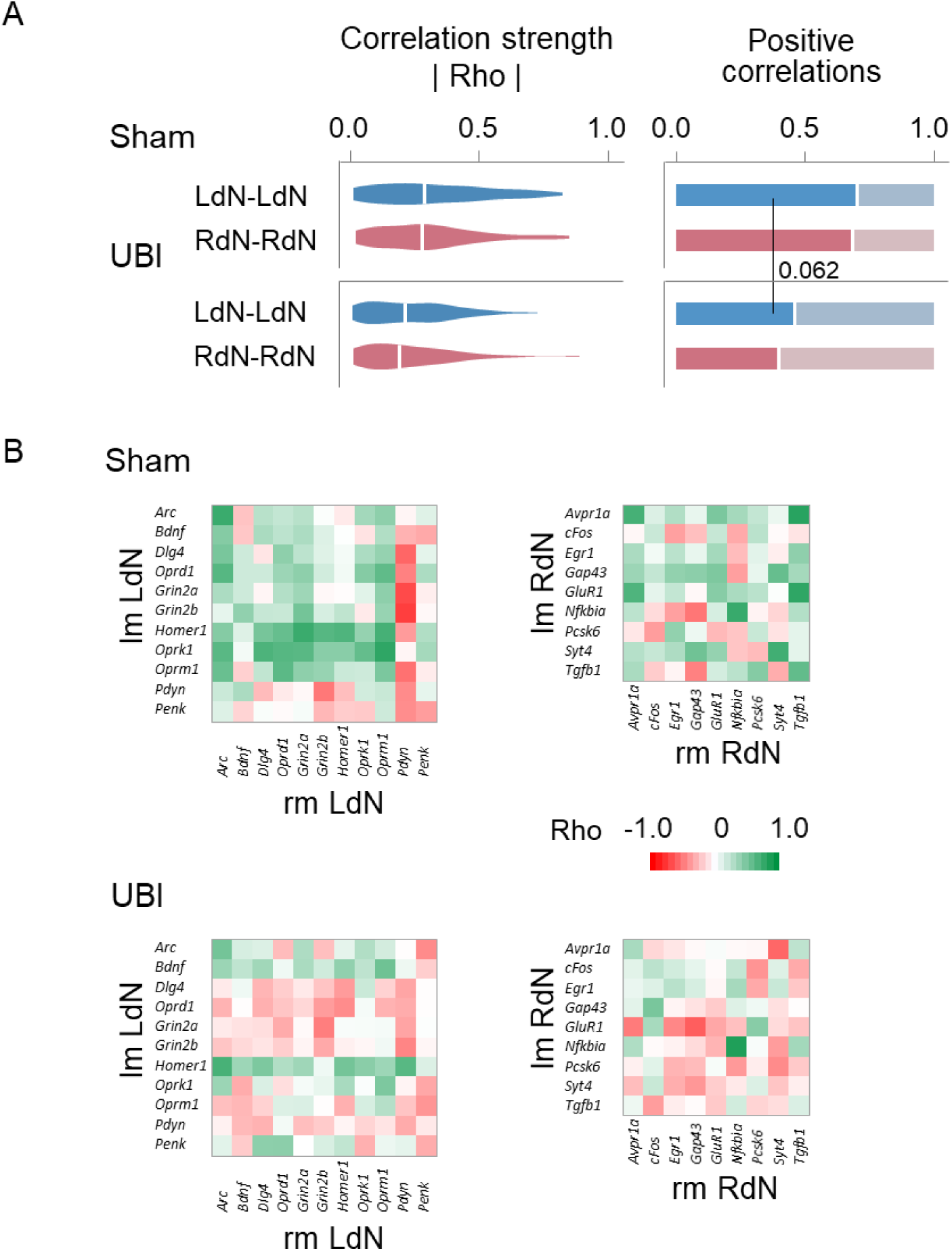
Gene-gene co-expression patterns in the spinal cord of the sham surgery and UBI rats. (**A**). The coordination strength and the proportion of positive correlations for the inter-modular correlations of the LdN (lmLdN-rmLdN) and RdN (lmRdN-rmRdN). (**B**). Heatmaps for Spearman’s rank coefficients for pairwise gene-gene the inter modular correlations of the LdN (lmLdN-rmLdN) and RdN (lmRdN-rmRdN). Comparison of the patterns between the LdN and RdN, and each of them between sham surgery and UBI groups. P values were determined by permutation testing with Benjamini-Hochberg family-wise multiple test correction, and are shown for the median AI of the combined sham surgery and UBI group. P values shown for the contrasts that are significant (P ≤ 0.05) after the correction for all three categorization variants, or for two of them while in the third variant P was < 0.05 and < 0.10 before and after the correction, respectively. **Source data:** The EXCEL source data file “SpinalC_SO_UBI_Ctrl_RD_DD.xlsx; Table III-S6 23 05 10.xlsx; raw_groups.xlsx”.

**Figure 6—figure supplement 1.**
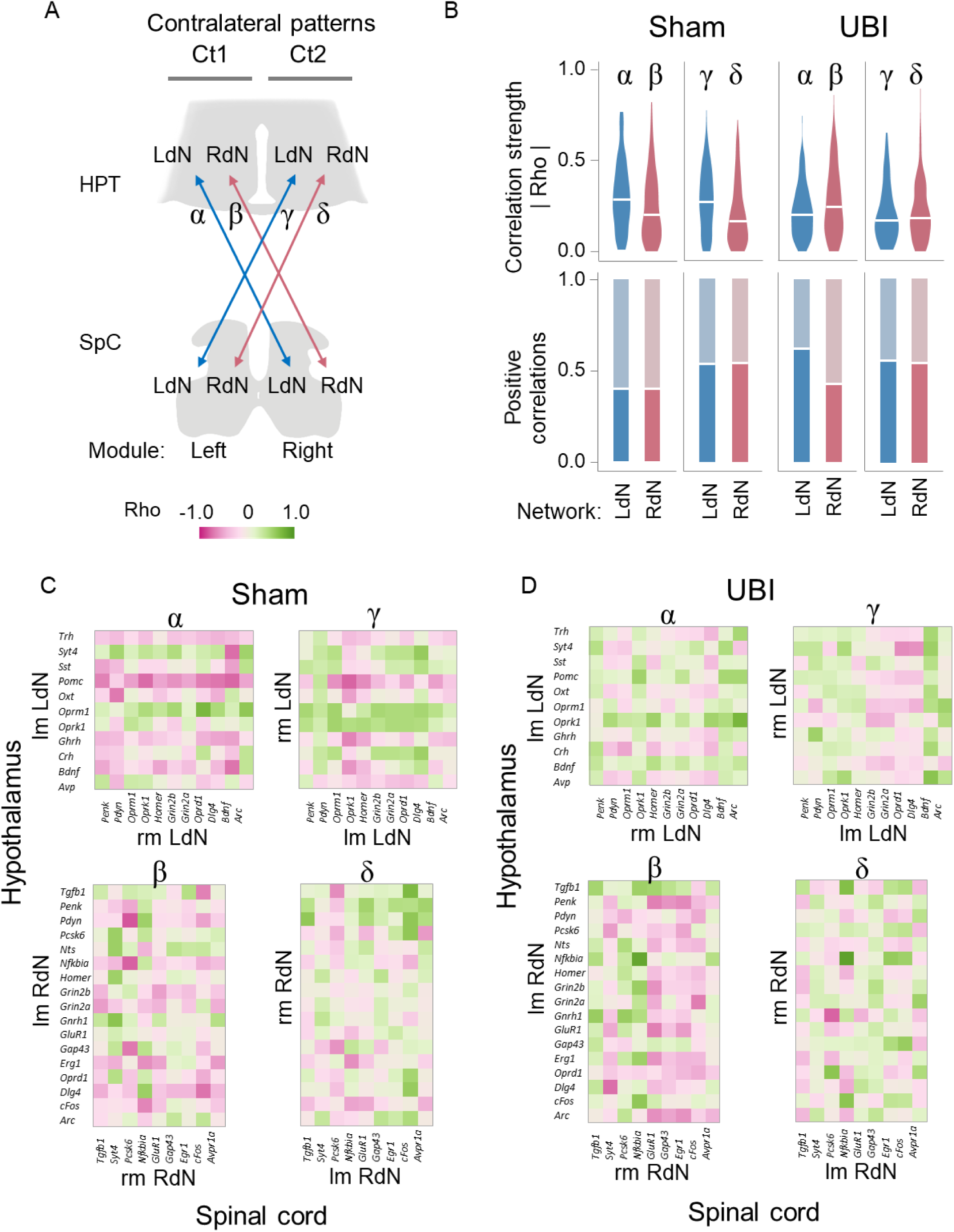
Coordination of the contralateral expression patterns of the left and right gene networks between the hypothalamus and lumbar spinal cord. The effects of UBI. Experimental details and design along with statistical analysis are described in legends to **Figures 4-6**. (**A**) Analyzed patterns of the contralateral pairwise gene-gene Spearman rank correlations between the left hypothalamus and right spinal cord (Ct1: α and β), and between right hypothalamus and left spinal cord (Ct2: γ and δ). (**B**) The coordination strength and the proportion of positive correlations for the Ct1 and Ct2 correlation patterns depicted in (**A**) in the sham surgery and UBI groups. The correlation patterns were compared between the LdN and RdN (α vs. β; γ vs. δ); each of them between the left and right modules (α vs. γ; β vs. δ), and all the patterns between UBI and sham surgery groups. (**C**,**D**) Heatmaps for Spearman’s rank coefficients for pairwise gene-gene correlations in sham surgery and UBI groups. **Source data:** The EXCEL source data file “Hypoth_SO_UBI.xlsx; SpinalC_SO_UBI_Ctrl_RD_DD.xlsx; Table III-S6 23 05 10.xlsx; raw_groups.xlsx”.

## REFERENCES

Adkins, D. L., Boychuk, J., Remple, M. S., & Kleim, J. A. (2006). Motor training induces experience specific patterns of plasticity across motor cortex and spinal cord. J Appl Physiol (1985), 101(6), 1776–1782. doi:10.1152/japplphysiol.00515.2006

Allen, H. N., Bobnar, H. J., & Kolber, B. J. (2021). Left and right hemispheric lateralization of the amygdala in pain. Prog Neurobiol, 196, 101891. doi:10.1016/j.pneurobio.2020.101891

Anderson, M. F., & Winterson, B. J. (1995). Properties of peripherally induced persistent hindlimb flexion in rat: involvement of N-methyl-D-aspartate receptors and capsaicin-sensitive afferents. Brain Res, 678(1-2), 140–150. doi:10.1016/0006-8993(95)00177-r

Antonucci, L. A., Di Carlo, P., Passiatore, R., Papalino, M., Monda, A., Amoroso, N., . . . Blasi, G. (2019). Thalamic connectivity measured with fMRI is associated with a polygenic index predicting thalamo-prefrontal gene co-expression. Brain Struct Funct, 224(3), 1331–1344. doi:10.1007/s00429-019-01843-7

Bakalkin, G. (2022). The left-right side-specific endocrine signaling in the effects of brain lesions: questioning of the neurological dogma. Cell Mol Life Sci, 79(11), 545. doi:10.1007/s00018-022-04576-9

Bakalkin, G., Iarygin, K., Kobylianskiĭ, A., Samovilova, N., & Klement’ev, B. (1981). Postural asymmetry induction by factors of the right and left hemispheres. Dokl Akad Nauk SSSR, 260(5), 1271– 1275.

Bakalkin, G., Iarygin, K. N., Trushina, E. D., Titov, M. I., & Smirnov, V. N. (1980). [Preferential development of flexion of the left or right hindlimb as a result of treatment with methionine enkephalin or leucine-enkephalin, respectively]. Dokl Akad Nauk SSSR, 252(3), 762–765. Retrieved from http://www.ncbi.nlm.nih.gov/pubmed/6250777

Bakalkin, G., & Kobylyansky, A. G. (1989). Opioids induce postural asymmetry in spinal rat: the side of the flexed limb depends upon the type of opioid agonist. Brain Res, 480(1-2), 277–289. doi:10.1016/0006-8993(89)90193-5

Bakalkin, G., Pivovarov, A., Kobylyansky, A. G., Yarygin, K. N., & Akparov, V. (1989). Ipsilateral responses induced by factors present in left and right hemispheres. Int Journal of Neuroscience, 47, 217–230.

Bakalkin, G., Pivovarov, A. S., Kobylyansky, A. G., Yarygin, K. N., & Akparov, V. (1989). Ipsilateral responses induced by factors present in left and right hemispheres. Int J Neurosci, 47(3-4), 217–230. doi:10.3109/00207458908987436

Bakalkin, G., Tsibezov, V. V., Sjutkin, E. A., Veselova, S. P., Novikov, I. D., & Krivosheev, O. G. (1984). Lateralization of LH-RH in rat hypothalamus. Brain Res, 296(2), 361–364. doi:10.1016/0006-8993(84)90074-x

Bakalkin, G. Y., Kobylyansky, A. G., Nagornaya, L. V., Yarygin, K. N., & Titov, M. I. (1986). Met enkephalin-induced release into the blood of a factor causing postural asymmetry. Peptides, 7(4), 551–556. Retrieved from http://www.ncbi.nlm.nih.gov/pubmed/3763433

Baude, M., Nielsen, J. B., & Gracies, J. M. (2018). The neurophysiology of deforming spastic paresis: A revised taxonomy. Ann Phys Rehabil Med. doi:10.1016/j.rehab.2018.10.004

Borson-Chazot, F., Jordan, D., Fevre-Montange, M., Kopp, N., Tourniaire, J., Rouzioux, J. M., . . . Mornex, R. (1986). TRH and LH-RH distribution in discrete nuclei of the human hypothalamus: evidence for a left prominence of TRH. Brain Res, 382(2), 433–436. doi:10.1016/0006-8993(86)91358-2

Boulton, D., Taylor, C. E., Green, S., & Macefield, V. G. (2021). The Role of Central Command in the Increase in Muscle Sympathetic Nerve Activity to Contracting Muscle During High Intensity Isometric Exercise. Front Neurosci, 15, 770072. doi:10.3389/fnins.2021.770072

Buisson, A., Lesne, S., Docagne, F., Ali, C., Nicole, O., MacKenzie, E. T., & Vivien, D. (2003). Transforming growth factor-beta and ischemic brain injury. Cell Mol Neurobiol, 23(4-5), 539–550. Retrieved from http://www.ncbi.nlm.nih.gov/pubmed/14514014

Burkner, P.C. (2021). Bayesian Item Response Modeling in R with brms and Stan. Journal of Statistical Software, 100(5), 1–54. doi:10.18637/jss.v100.i05

Bustin, S. A., Benes, V., Garson, J. A., Hellemans, J., Huggett, J., Kubista, M., . . . Wittwer, C. T. (2009). The MIQE guidelines: minimum information for publication of quantitative real-time PCR experiments. Clin Chem, 55(4), 611–622. doi:10.1373/clinchem.2008.112797

Canty A, R. B. (2022). boot: Bootstrap R (S-Plus) Functions. R package version 1.3–28.1.

Chazov, E. I., Bakalkin, G., Yarigin, K. N., Trushina, E. D., Titov, M. I., & Smirnov, V. N. (1981). Enkephalins induce asymmetrical effects on posture in the rat. Experientia, 37(8), 887–889. Retrieved from http://www.ncbi.nlm.nih.gov/pubmed/7286146

Chepurnov, S. A. (1994). [Role of neuropeptides in the lateralization of brain (as exemplified by behavior in radial maze]. Vestn Ross Akad Med Nauk(2), 36–40. Retrieved from https://www.ncbi.nlm.nih.gov/pubmed/7513579

Concha, M. L., Bianco, I. H., & Wilson, S. W. (2012). Encoding asymmetry within neural circuits. Nat Rev Neurosci, 13(12), 832–843. doi:10.1038/nrn3371

Cruz, M. E., Flores, A., & Dominguez, R. (2014). The cholinergic system of the preoptic-anterior hypothalamic areas regulates the ovarian follicular population in an asymmetric way. Endocrine, 47(3), 913–922. doi:10.1007/s12020-014-0266-2

de Kovel, C. G. F., Lisgo, S., Karlebach, G., Ju, J., Cheng, G., Fisher, S. E., & Francks, C. (2017). Left-Right Asymmetry of Maturation Rates in Human Embryonic Neural Development. Biol Psychiatry, 82(3), 204–212. doi:10.1016/j.biopsych.2017.01.016

de la Iglesia, H. O., Meyer, J., & Schwartz, W. J. (2003). Lateralization of circadian pacemaker output: Activation of left- and right-sided luteinizing hormone-releasing hormone neurons involves a neural rather than a humoral pathway. J Neurosci, 23(19), 7412–7414. doi:10.1523/JNEUROSCI.23-19-07412.2003

Deliagina, T. G., Orlovsky, G. N., Selverston, A. I., & Arshavsky, Y. I. (2000). Asymmetrical effect of GABA on the postural orientation in Clione. J Neurophysiol, 84(3), 1673–1676. doi:10.1152/jn.2000.84.3.1673

Dewald, J. P., Beer, R. F., Given, J. D., McGuire, J. R., & Rymer, W. Z. (1999). Reorganization of flexion reflexes in the upper extremity of hemiparetic subjects. Muscle Nerve, 22(9), 1209–1221. doi:10.1002/(sici)1097-4598(199909)22:9<1209::aid-mus7>3.0.co;2-b

Dobrin, R., Zhu, J., Molony, C., Argman, C., Parrish, M. L., Carlson, S., . . . Schadt, E. E. (2009). Multi tissue coexpression networks reveal unexpected subnetworks associated with disease. Genome Biol, 10(5), R55. doi:10.1186/gb-2009-10-5-r55

Dolan, S., Hastie, P., Crossan, C., & Nolan, A. M. (2011). Co-induction of cyclooxygenase-2 [correction of cyclooxyenase-2] and early growth response gene (Egr-1) in spinal cord in a clinical model of persistent inflammation and hyperalgesia. Mol Pain, 7, 91. doi:10.1186/1744-8069-7-91

Duboc, V., Dufourcq, P., Blader, P., & Roussigne, M. (2015). Asymmetry of the Brain: Development and Implications. Annu Rev Genet, 49, 647–672. doi:10.1146/annurev-genet-112414-055322

Efimova, E. V., Chepurnova, N. E., & Chepurnov, S. A. (1989). [Thyroliberin-induced motor asymmetry as an indicator of the individual behavioral strategy of rats in a radial labyrinth]. Nauchnye Doki Vyss Shkoly Biol Nauki(3), 23–27. Retrieved from https://www.ncbi.nlm.nih.gov/pubmed/2500987

Epstein, I., & Finkbeiner, S. (2018). The Arc of cognition: Signaling cascades regulating Arc and implications for cognitive function and disease. Semin Cell Dev Biol, 77, 63–72. doi:10.1016/j.semcdb.2017.09.023

Erola, P., Bjorkegren, J. L. M., & Michoel, T. (2020). Model-based clustering of multi-tissue gene expression data. Bioinformatics, 36(6), 1807–1813. doi:10.1093/bioinformatics/btz805

Farmer, D. G. S., Pracejus, N., Dempsey, B., Turner, A., Bokiniec, P., Paton, J. F. R., . . . McMullan, S. (2019). On the presence and functional significance of sympathetic premotor neurons with collateralized spinal axons in the rat. J Physiol, 597(13), 3407–3423. doi:10.1113/JP277661

Fu, Y., Liu, Q., Anrather, J., & Shi, F. D. (2015). Immune interventions in stroke. Nat Rev Neurol, 11(9), 524–535. doi:10.1038/nrneurol.2015.144

Gelman, A. (2019). Prior Choice Recommendations. Stan-dev/stan. Retrieved from https://github.com/stan-dev/stan/wiki/Prior-Choice-Recommendations

Gerring, Z. F., Gamazon, E. R., Derks, E. M., & Major Depressive Disorder Working Group of the Psychiatric Genomics, C. (2019). A gene co-expression network-based analysis of multiple brain tissues reveals novel genes and molecular pathways underlying major depression. PLoS Genet, 15(7), e1008245. doi:10.1371/journal.pgen.1008245

Goldberger, M. E. (1988). Partial and complete deafferentation of cat hindlimb: the contribution of behavioral substitution to recovery of motor function. Exp Brain Res, 73(2), 343–353. doi:10.1007/BF00248226

Gracies, J. M. (2005). Pathophysiology of spastic paresis. I: Paresis and soft tissue changes. Muscle Nerve, 31(5), 535–551. doi:10.1002/mus.20284

Grasselli, G., & Strata, P. (2013). Structural plasticity of climbing fibers and the growth-associated protein GAP-43. Front Neural Circuits, 7, 25. doi:10.3389/fncir.2013.00025

Gunturkun, O., Strockens, F., & Ocklenburg, S. (2020). Brain Lateralization: A Comparative Perspective. Physiol Rev, 100(3), 1019–1063. doi:10.1152/physrev.00006.2019

Hall, R. D., & Lindholm, E. (1974). Organization of motor and somatosensory neocortex in the albino rat. Brain Res, 66, 23–38.

Harris, K. P., Zhang, Y. V., Piccioli, Z. D., Perrimon, N., & Littleton, J. T. (2016). The postsynaptic t-SNARE Syntaxin 4 controls traffic of Neuroligin 1 and Synaptotagmin 4 to regulate retrograde signaling. Elife, 5. doi:10.7554/eLife.13881

Hayashi, M., Ueyama, T., Nemoto, K., Tamaki, T., & Senba, E. (2000). Sequential mRNA expression for immediate early genes, cytokines, and neurotrophins in spinal cord injury. J Neurotrauma, 17(3), 203–218. doi:10.1089/neu.2000.17.203

Horan, P., Taylor, J., Yamamura, H. I., & Porreca, F. (1992). Extremely long-lasting antagonistic actions of nor-binaltorphimine (nor-BNI) in the mouse tail-flick test. J Pharmacol Exp Ther, 260(3), 1237–1243. Retrieved from https://www.ncbi.nlm.nih.gov/pubmed/1312164

Hotta, H., Iimura, K., Watanabe, N., & Shigemoto, K. (2021). Maintenance of contractile force of the hind limb muscles by the somato-lumbar sympathetic reflexes. J Physiol Sci, 71(1), 15. doi:10.1186/s12576-021-00799-w

Hultborn, H., & Malmsten, J. (1983a). Changes in segmental reflexes following chronic spinal cord hemisection in the cat. I. Increased monosynaptic and polysynaptic ventral root discharges. Acta Physiol Scand, 119(4), 405–422. doi:10.1111/j.1748-1716.1983.tb07357.x

Hultborn, H., & Malmsten, J. (1983b). Changes in segmental reflexes following chronic spinal cord hemisection in the cat. II. Conditioned monosynaptic test reflexes. Acta Physiol Scand, 119(4), 423–433. doi:10.1111/j.1748-1716.1983.tb07358.x

Hussain, Z. M., Fitting, S., Watanabe, H., Usynin, I., Yakovleva, T., Knapp, P. E., . . . Bakalkin, G. (2012). Lateralized response of dynorphin a peptide levels after traumatic brain injury. J Neurotrauma, 29(9), 1785–1793. doi:10.1089/neu.2011.2286

Joynes, R. L., Janjua, K., & Grau, J. W. (2004). Instrumental learning within the spinal cord: VI. The NMDA receptor antagonist, AP5, disrupts the acquisition and maintenance of an acquired flexion response. Behav Brain Res, 154(2), 431–438. doi:10.1016/j.bbr.2004.03.030

Kantonen, T., Karjalainen, T., Isojarvi, J., Nuutila, P., Tuisku, J., Rinne, J., . . . Nummenmaa, L. (2020). Interindividual variability and lateralization of mu-opioid receptors in the human brain. Neuroimage, 217, 116922. doi:10.1016/j.neuroimage.2020.116922

Kawakami, R., Shinohara, Y., Kato, Y., Sugiyama, H., Shigemoto, R., & Ito, I. (2003). Asymmetrical allocation of NMDA receptor epsilon2 subunits in hippocampal circuitry. Science, 300(5621), 990–994. doi:10.1126/science.1082609

Klement’ev, B. I., Molokoedov, A. S., Bushuev, V. N., Danilovskii, M. A., & Sepetov, N. F. (1986). [Isolation of the postural asymmetry factor following right hemisection of the spinal cord]. Dokl Akad Nauk SSSR, 291(3), 737–741. Retrieved from http://www.ncbi.nlm.nih.gov/pubmed/3803179

Kononenko, O., Galatenko, V., Andersson, M., Bazov, I., Watanabe, H., Zhou, X. W., . . . Bakalkin, G. (2017). Intra- and interregional coregulation of opioid genes: broken symmetry in spinal circuits. FASEB J, 31(5), 1953–1963. doi:10.1096/fj.201601039R

Kononenko, O., Mityakina, I., Galatenko, V., Watanabe, H., Bazov, I., Gerashchenko, A., . . . Bakalkin, G. (2018). Differential effects of left and right neuropathy on opioid gene expression in lumbar spinal cord. Brain Res, 1695, 78–83. doi:10.1016/j.brainres.2018.05.043

Kruschke, J. (2015). Doing Bayesian Data Analysis: Academic Press.

Kryzhanovskii, G. N., Lutsenko, V. K., Karganov, M., & Beliaev, S. V. (1984). [Lateralization of peptide distribution in the brain and the asymmetry of motor control]. Patol Fiziol Eksp Ter(3), 68–71. Retrieved from http://www.ncbi.nlm.nih.gov/pubmed/6332294

Larsson, M., & Broman, J. (2008). Translocation of GluR1-containing AMPA receptors to a spinal nociceptive synapse during acute noxious stimulation. J Neurosci, 28(28), 7084–7090. doi:10.1523/JNEUROSCI.5749-07.2008

Lavrov, I., Courtine, G., Dy, C. J., van den Brand, R., Fong, A. J., Gerasimenko, Y., . . . Edgerton, V. R. (2008). Facilitation of stepping with epidural stimulation in spinal rats: role of sensory input. J Neurosci, 28(31), 7774–7780. doi:10.1523/JNEUROSCI.1069-08.2008

Lee, T. K., Lois, J. H., Troupe, J. H., Wilson, T. D., & Yates, B. J. (2007). Transneuronal tracing of neural pathways that regulate hindlimb muscle blood flow. Am J Physiol Regul Integr Comp Physiol, 292(4), R1532–1541. doi:10.1152/ajpregu.00633.2006

Lemon, R. N. (2008). Descending pathways in motor control. Annu Rev Neurosci, 31, 195–218. doi:10.1146/annurev.neuro.31.060407.125547

Lenth, R. (2023). emmeans: Estimated Marginal Means, aka Least-Squares Means. R package version 1.8.5. Retrieved from https://cran.r-project.org/package=emmeans

Lorentzen, J., Pradines, M., Gracies, J. M., & Bo Nielsen, J. (2018). On Denny-Brown’s ’spastic dystonia’ - What is it and what causes it? Clin Neurophysiol, 129(1), 89–94. doi:10.1016/j.clinph.2017.10.023

Louis, E. D. (1994). Contralateral control: evolving concepts of the brain-body relationship from Hippocrates to Morgagni. Neurology, 44(12), 2398–2400. doi:10.1212/wnl.44.12.2398

Lukoyanov, N., Watanabe, H., Carvalho, L. S., Kononenko, O., Sarkisyan, D., Zhang, M., . . . Bakalkin, G. (2021). Left-right side-specific endocrine signaling complements neural pathways to mediate acute asymmetric effects of brain injury. Elife, 10. doi:10.7554/eLife.65247

MacNeilage, P. F., Rogers, L. J., & Vallortigara, G. (2009). Origins of the left & right brain. Sci Am, 301(1), 60–67. doi:10.1038/scientificamerican0709-60

Malmsten, J. (1983). Time course of segmental reflex changes after chronic spinal cord hemisection in the rat. Acta Physiol Scand, 119(4), 435–443. doi:10.1111/j.1748-1716.1983.tb07359.x

Marinelli, L., Curra, A., Trompetto, C., Capello, E., Serrati, C., Fattapposta, F., . . . Bandini, F. (2017). Spasticity and spastic dystonia: the two faces of velocity-dependent hypertonia. J Electromyogr Kinesiol, 37, 84–89. doi:10.1016/j.jelekin.2017.09.005

Marlin, B. J., Mitre, M., D’Amour J, A., Chao, M. V., & Froemke, R. C. (2015). Oxytocin enables maternal behaviour by balancing cortical inhibition. Nature, 520(7548), 499–504. doi:10.1038/nature14402

Marsala, M., Hefferan, M. P., Kakinohana, O., Nakamura, S., Marsala, J., & Tomori, Z. (2005). Measurement of peripheral muscle resistance in rats with chronic ischemia-induced paraplegia or morphine-induced rigidity using a semi-automated computer-controlled muscle resistance meter. J Neurotrauma, 22(11), 1348–1361. doi:10.1089/neu.2005.22.1348

McCall, A. A., Miller, D. M., & Yates, B. J. (2017). Descending Influences on Vestibulospinal and Vestibulosympathetic Reflexes. Front Neurol, 8, 112. doi:10.3389/fneur.2017.00112

McElreath, R. (2019). Statistical Rethinking. A Bayesian Course with Examples in R and Stan (& PyMC3 & brms & Julia too): Chapman and Hall/CRC.

Moran, J. L., Cruz, M. E., & Dominquez, R. (1994). Differences in the ovulatory response to unilateral lesions in the preoptic or anterior hypothalamic area performed on each day of the estrous cycle of adult rats. Brain Res Bull, 33(6), 663–668. doi:10.1016/0361-9230(94)90230-5

Nathan, P. W., Smith, M. C., & Deacon, P. (1990). The corticospinal tracts in man. Course and location of fibres at different segmental levels. Brain, 113 ( *Pt 2*), 303–324. doi:10.1093/brain/113.2.303

Nation, K. M., De Felice, M., Hernandez, P. I., Dodick, D. W., Neugebauer, V., Navratilova, E., & Porreca, F. (2018). Lateralized kappa opioid receptor signaling from the amygdala central nucleus promotes stress-induced functional pain. Pain, 159(5), 919–928. doi:10.1097/j.pain.0000000000001167

Ng, S. Y., & Lee, A. Y. W. (2019). Traumatic Brain Injuries: Pathophysiology and Potential Therapeutic Targets. Front Cell Neurosci, 13, 528. doi:10.3389/fncel.2019.00528

Nizhnikov, M. E., Pautassi, R. M., Truxell, E., & Spear, N. E. (2009). Opioid antagonists block the acquisition of ethanol-mediated conditioned tactile preference in infant rats. Alcohol, 43(5), 347–358. doi:10.1016/j.alcohol.2009.06.001

Noguchi, T., Ohta, S., Kakinoki, R., Kaizawa, Y., & Matsuda, S. (2013). A new cervical nerve root avulsion model using a posterior extra-vertebral approach in rats. J Brachial Plex Peripher Nerve Inj, 8(1), 8. doi:10.1186/1749-7221-8-8

Nordez, A., Casari, P., & Cornu, C. (2008). Effects of stretching velocity on passive resistance developed by the knee musculo-articular complex: contributions of frictional and viscoelastic behaviours. Eur J Appl Physiol, 103(2), 243–250. doi:10.1007/s00421-008-0695-9

Nordez, A., Casari, P., Mariot, J. P., & Cornu, C. (2009). Modeling of the passive mechanical properties of the musculo-articular complex: acute effects of cyclic and static stretching. J Biomech, 42(6), 767–773. doi:10.1016/j.jbiomech.2008.12.019

Norris, J. N., Perez-Acosta, A. M., Ortega, L. A., & Papini, M. R. (2009). Naloxone facilitates appetitive extinction and eliminates escape from frustration. Pharmacol Biochem Behav, 94(1), 81–87. doi:10.1016/j.pbb.2009.07.012

O’Mahony, A., Raber, J., Montano, M., Foehr, E., Han, V., Lu, S. M., . . . Greene, W. C. (2006). NF-kappaB/Rel regulates inhibitory and excitatory neuronal function and synaptic plasticity. Mol Cell Biol, 26(19), 7283–7298. doi:10.1128/MCB.00510-06

Ocklenburg, S., Schmitz, J., Moinfar, Z., Moser, D., Klose, R., Lor, S., . . . Gunturkun, O. (2017). Epigenetic regulation of lateralized fetal spinal gene expression underlies hemispheric asymmetries. Elife, 6. doi:10.7554/eLife.22784

Patkar, K. A., Wu, J., Ganno, M. L., Singh, H. D., Ross, N. C., Rasakham, K., . . . McLaughlin, J. P. (2013). Physical presence of nor-binaltorphimine in mouse brain over 21 days after a single administration corresponds to its long-lasting antagonistic effect on kappa-opioid receptors. J Pharmacol Exp Ther, 346(3), 545–554. doi:10.1124/jpet.113.206086

Paxinos, G., & Watson, C. (2007). The Rat Brain in Stereotaxic Coordinates Academic Press

Petrillo, P., Angelici, O., Bingham, S., Ficalora, G., Garnier, M., Zaratin, P. F., . . . Scheideler, M. A. (2003). Evidence for a selective role of the delta-opioid agonist [8R-(4bS*,8aalpha,8abeta, 12bbeta)]7,10-Dimethyl-1-methoxy-11-(2-methylpropyl)oxycarbonyl 5,6,7,8,12,12b-hexahydro-(9H)-4,8-methanobenzofuro[3,2-e]pyrrolo[2,3-g]isoquinoli ne hydrochloride (SB-235863) in blocking hyperalgesia associated with inflammatory and neuropathic pain responses. J Pharmacol Exp Ther, 307(3), 1079–1089. doi:10.1124/jpet.103.055590

Phelps, C. E., Navratilova, E., Dickenson, A. H., Porreca, F., & Bannister, K. (2019). Kappa opioid signaling in the right central amygdala causes hind paw specific loss of diffuse noxious inhibitory controls in experimental neuropathic pain. Pain, 160(7), 1614–1621. doi:10.1097/j.pain.0000000000001553

Purves, D., Augustine, G. J., & Fitzpatrick, D. (2001). Neuroscience. 2nd edition. Sunderland: Oxford University Press.

Roper, J., O’Carroll, A. M., Young, W., 3rd, & Lolait, S. (2011). The vasopressin Avpr1b receptor: molecular and pharmacological studies. Stress, 14(1), 98–115. doi:10.3109/10253890.2010.512376

Rutten, K., Schroder, W., Christoph, T., Koch, T., & Tzschentke, T. M. (2018). Selectivity profiling of NOP, MOP, DOP and KOP receptor antagonists in the rat spinal nerve ligation model of mononeuropathic pain. Eur J Pharmacol, 827, 41–48. doi:10.1016/j.ejphar.2018.03.008

R Core Team (2022). R: A language and environment for statistical computing. R Foundation for Statistical Computing, Vienna, Austria. URL https://www.R-project.org/

Santibanez, J. F., Quintanilla, M., & Bernabeu, C. (2011). TGF-beta/TGF-beta receptor system and its role in physiological and pathological conditions. Clin Sci (Lond), 121(6), 233–251. doi:10.1042/CS20110086

Schutta, H. S., Abu-Amero, K. K., & Bosley, T. M. (2010). Exceptions to the Valsalva doctrine. Neurology, 74(4), 329–335. doi:10.1212/WNL.0b013e3181cbcd84

Serrao, M., Ranavolo, A., Andersen, O. K., Don, R., Draicchio, F., Conte, C., . . . Pierelli, F. (2012). Reorganization of multi-muscle and joint withdrawal reflex during arm movements in post stroke hemiparetic patients. Clin Neurophysiol, 123(3), 527–540. doi:10.1016/j.clinph.2011.07.031

Sheean, G., & McGuire, J. R. (2009). Spastic hypertonia and movement disorders: pathophysiology, clinical presentation, and quantification. PM R, 1(9), 827–833. doi:10.1016/j.pmrj.2009.08.002

Simon, F. (2018). On the origin of the term decussatio pyramidum. J Hist Neurosci, 27(1), 101–105. doi:10.1080/0964704X.2017.1285512

Smith, C. C., Paton, J. F. R., Chakrabarty, S., & Ichiyama, R. M. (2017). Descending Systems Direct Development of Key Spinal Motor Circuits. J Neurosci, 37(26), 6372–6387. doi:10.1523/JNEUROSCI.0149-17.2017

Soriano, J. R., Daniels, N., Prinsen, J., & Alaerts, K. (2020). Intranasal oxytocin enhances approach related EEG frontal alpha asymmetry during engagement of direct eye contact. Brain Commun, 2(2), fcaa093. doi:10.1093/braincomms/fcaa093

Stan Development Teamw (2022). RStan: the R interface to Stan. R package version 2.21.7. https://mc-stan.org/

Spaich, E. G., Hinge, H. H., Arendt-Nielsen, L., & Andersen, O. K. (2006). Modulation of the withdrawal reflex during hemiplegic gait: effect of stimulation site and gait phase. Clin Neurophysiol, 117(11), 2482–2495. doi:10.1016/j.clinph.2006.07.139

Takahashi, Y., & Nakajima, Y. (1996). Dermatomes in the rat limbs as determined by antidromic stimulation of sensory C-fibers in spinal nerves. Pain, 67(1), 197–202. doi:10.1016/0304-3959(96)03116-8

Tappe, A., Klugmann, M., Luo, C., Hirlinger, D., Agarwal, N., Benrath, J., . . . Kuner, R. (2006). Synaptic scaffolding protein Homer1a protects against chronic inflammatory pain. Nat Med, 12(6), 677–681. doi:10.1038/nm1406

Taylor, S. C., Nadeau, K., Abbasi, M., Lachance, C., Nguyen, M., & Fenrich, J. (2019). The Ultimate qPCR Experiment: Producing Publication Quality, Reproducible Data the First Time. Trends Biotechnol, 37(7), 761–774. doi:10.1016/j.tibtech.2018.12.002

Team, R. C. (2022). R: A language and environment for statistical computing. R Foundation for Statistical (Version R 4.2). Vienna, Austria. Retrieved from https://www.r-project.org/

Team, S. D. (2023). Stan Modeling Language Users Guide and Reference Manual, . Retrieved from https://mc-stan.org. https://mc-stan.org

Thulin, M. (2021). Modern Statistics with R.

Vandesompele, J., De Preter, K., Pattyn, F., Poppe, B., Van Roy, N., De Paepe, A., & Speleman, F. (2002). Accurate normalization of real-time quantitative RT-PCR data by geometric averaging of multiple internal control genes. Genome Biol, 3(7), RESEARCH0034. doi:10.1186/gb-2002-3-7-research0034

Vartanian, G., Shatik, S., Tokarev, A., & Klement’ev, B. (1989). The activity of postural asymmetry factors in symmetrical sections of the rat spinal cord. Biull Eksp Biol Med., 107(4), 404–407.

Vavrek, R., Girgis, J., Tetzlaff, W., Hiebert, G. W., & Fouad, K. (2006). BDNF promotes connections of corticospinal neurons onto spared descending interneurons in spinal cord injured rats. Brain, 129(Pt 6), 1534–1545. doi:10.1093/brain/awl087

Vulliemoz, S., Raineteau, O., & Jabaudon, D. (2005). Reaching beyond the midline: why are human brains cross wired? Lancet Neurol, 4(2), 87–99. doi:10.1016/S1474-4422(05)00990-7

Watanabe, H., Fitting, S., Hussain, M. Z., Kononenko, O., Iatsyshyna, A., Yoshitake, T., . . . Bakalkin, G. (2015). Asymmetry of the endogenous opioid system in the human anterior cingulate: a putative molecular basis for lateralization of emotions and pain. Cereb Cortex, 25(1), 97–108. doi:10.1093/cercor/bht204

Watanabe, H., Nosova, O., Sarkisyan, D., Andersen, M. S., Carvalho, L., Galatenko, V., . . . Bakalkin, G. (2021). Left-right side-specific neuropeptide mechanism mediates contralateral responses to a unilateral brain injury eNeuro.

Watanabe, H., Nosova, O., Sarkisyan, D., Andersen, M. S., Zhang, M., Rorick-Kehn, L., . . . Bakalkin, G. (2020). Ipsilesional versus contralesional postural deficits induced by unilateral brain trauma: a side reversal by opioid mechanism. Brain Communications, 2(2). doi:10.1093/braincomms/fcaa208

Wilson, L., Stewart, W., Dams-O’Connor, K., Diaz-Arrastia, R., Horton, L., Menon, D. K., & Polinder, S. (2017). The chronic and evolving neurological consequences of traumatic brain injury. Lancet Neurol, 16(10), 813–825. doi:10.1016/S1474-4422(17)30279-X

Wolpaw, J. R. (2012). Harnessing neuroplasticity for clinical applications. Brain, 135(Pt 4), e215; author reply e216. doi:10.1093/brain/aws017

Won, S., Incontro, S., Nicoll, R. A., & Roche, K. W. (2016). PSD-95 stabilizes NMDA receptors by inducing the degradation of STEP61. Proc Natl Acad Sci U S A, 113(32), E4736–4744. doi:10.1073/pnas.1609702113

You, H. J., Morch, C. D., & Arendt-Nielsen, L. (2004). Electrophysiological characterization of facilitated spinal withdrawal reflex to repetitive electrical stimuli and its modulation by central glutamate receptor in spinal anesthetized rats. Brain Res, 1009(1-2), 110–119. doi:10.1016/j.brainres.2004.02.053

Zhang, M., Watanabe, H., Sarkisyan, D., Andersen, M. S., Nosova, O., Galatenko, V., . . . Bakalkin, G. (2020). Hindlimb motor responses to unilateral brain injury: spinal cord encoding and left-right asymmetry. Brain Communications, 2(1). doi:10.1093/braincomms/fcaa055

Zink, C. F., Kempf, L., Hakimi, S., Rainey, C. A., Stein, J. L., & Meyer-Lindenberg, A. (2011). Vasopressin modulates social recognition-related activity in the left temporoparietal junction in humans. Transl Psychiatry, 1, e3. doi:10.1038/tp.2011.2

